# Large-scale benchmarking of circRNA detection tools reveals large differences in sensitivity but not in precision

**DOI:** 10.1101/2022.12.06.519083

**Authors:** Marieke Vromman, Jasper Anckaert, Stefania Bortoluzzi, Alessia Buratin, Chia-Ying Chen, Qinjie Chu, Trees-Juen Chuang, Roozbeh Dehghannasiri, Christoph Dieterich, Xin Dong, Paul Flicek, Enrico Gaffo, Wanjun Gu, Chunjiang He, Steve Hoffmann, Osagie Izuogu, Michael S. Jackson, Tobias Jakobi, Eric C. Lai, Justine Nuytens, Julia Salzman, Mauro Santibanez-Koref, Peter Stadler, Olivier Thas, Eveline Vanden Eynde, Kimberly Verniers, Guoxia Wen, Jakub Westholm, Li Yang, Chu-Yu Ye, Nurten Yigit, Guo-Hua Yuan, Jinyang Zhang, Fangqing Zhao, Jo Vandesompele, Pieter-Jan Volders

**Affiliations:** OncoRNALab, Cancer Research Institute Ghent (CRIG), Department of Biomolecular Medicine, Ghent University, Belgium; Department of Molecular Medicine, University of Padova, Italy; Genomics Research Center, Academia Sinica, Taiwan; Institute of Crop Science & Institute of Bioinformatics, Zhejiang University, China; Department of Biomedical Data Science and of Biochemistry, Stanford University, USA; Klaus Tschira Institute for Integrative Computational Cardiology, Department of Internal Medicine III, University Hospital Heidelberg, German Center for Cardiovascular Research (DZHK), Germany; School of Basic Medical Science, Department of Medical Genetics, Wuhan University, China; EMBL-EBI, UK; Collaborative Innovation Center of Jiangsu Province of Cancer Prevention and Treatment of Chinese Medicine, School of Artificial Intelligence and Information Technology, Nanjing University of Chinese Medicine, China; Computational Biology Group, Leibniz Institute on Aging - Fritz Lipmann Institute (FLI), Jena, Germany; Biosciences Institute, Faculty of Medical Sciences, Newcastle University, UK; Translational Cardiovascular Research Center, University of Arizon - College of Medicine Phoenix, USA; Developmental Biology Program, Sloan Kettering Institute, New York, USA; Bioinformatics Group, Department of Computer Science, and Interdisciplinary Center for Bioinformatics, Universität Leipzig, Germany; Data Science Institute, I-Biostat, Hasselt University, Belgium; State Key Laboratory of Bioelectronics, School of Biological Sciences and Medical Engineering, Southeast University, China; Dept of Biochemistry and Biophysics, National Bioinformatics Infrastructure Sweden, Science for Life Laboratory, Stockholm University, Sweden; Center for Molecular Medicine, Children’s Hospital, Fudan University and Shanghai Key Laboratory of Medical Epigenetics, International Laboratory of Medical Epigenetics and Metabolism, Ministry of Science and Technology, Institutes of Biomedical Sciences, Fudan University, China; CAS Key Laboratory of Computational Biology, Shanghai Institute of Nutrition and Health, University of Chinese Academy of Sciences, Chinese Academy of Sciences, China; Beijing Institutes of Life Science, Chinese Academy of Sciences, China

## Abstract

The detection of circular RNA molecules (circRNAs) is typically based on short-read RNA sequencing data processed by computational detection tools. During the last decade, a plethora of such tools have been developed, but a systematic comparison with orthogonal validation is missing. Here, we set up a circRNA detection tool benchmarking study, in which 16 tools were used and detected over 315,000 unique circRNAs in three deeply sequenced human cell types. Next, 1,516 predicted circRNAs were empirically validated using three orthogonal methods. Generally, tool-specific precision values are high and similar (median of 98.8%, 96.3%, and 95.5% for qPCR, RNase R, and amplicon sequencing, respectively) whereas the sensitivity and number of predicted circRNAs (ranging from 1,372 to 58,032) are the most significant tool differentiators. Furthermore, we demonstrate the complementarity of tools through the increase in detection sensitivity by considering the union of highly-precise tool combinations while keeping the number of false discoveries low. Finally, based on the benchmarking results, recommendations are put forward for circRNA detection and validation.

## Introduction

Circular RNAs (circRNAs) are a class of non-coding RNA molecules numerously present in humans and other eukaryotic species. For a long time, circRNAs were regarded as unimportant byproducts of splicing. However, since the advancement of RNA sequencing technologies and the development of circRNA detection bioinformatics pipelines, there has been a significant increase in circRNA research, with a compound annual growth rate of scientific publications of 58% over the last five years (Figure 1A) (1).

**Figure 1.**
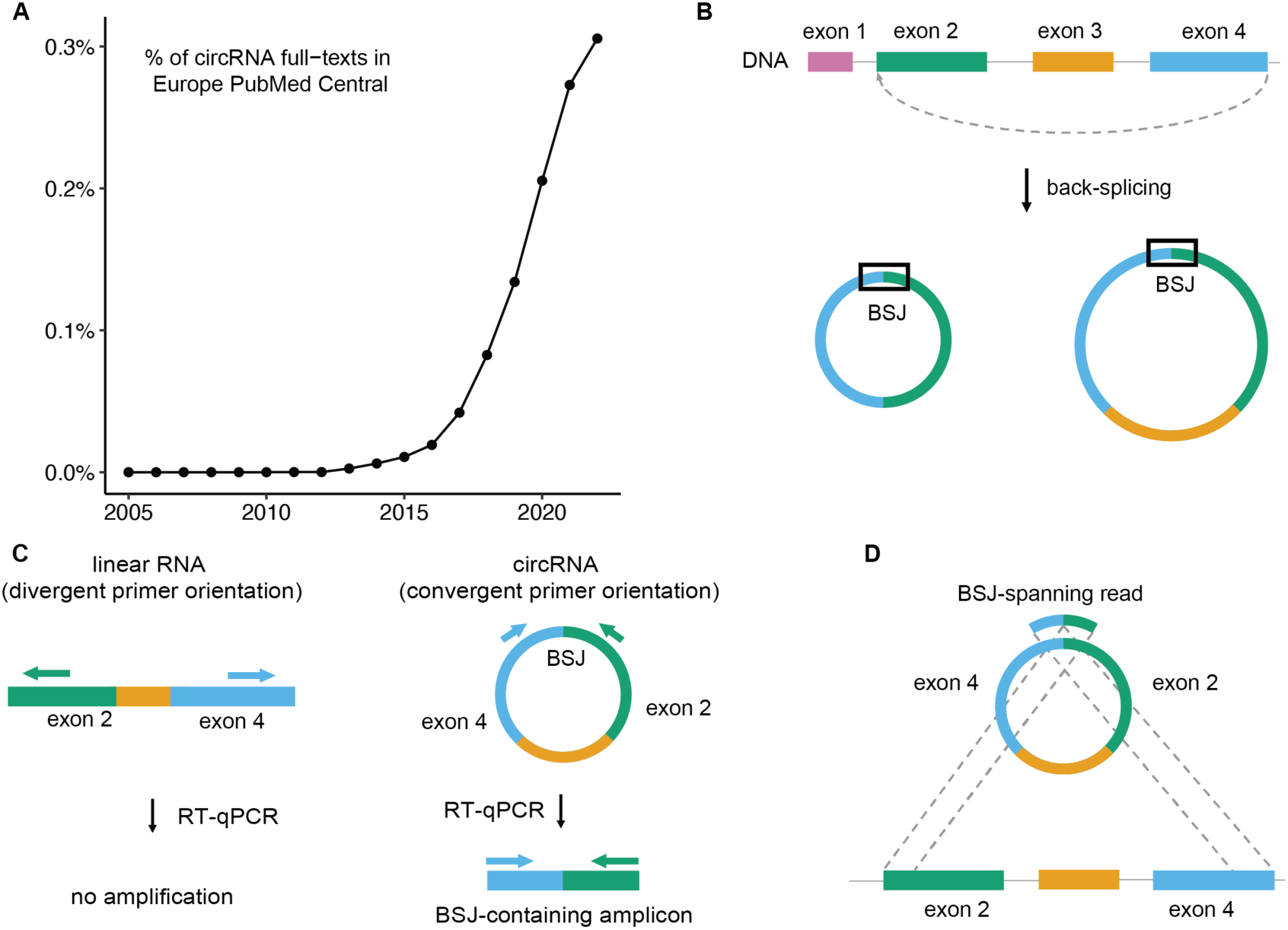
CircRNA scientific relevance, structure, and detection. **A.** Over the last decade, circRNA research has increased rapidly, as illustrated by the proportional growth of publications mentioning circRNA in Europe PubMed Central. **B.** CircRNAs are formed through back-splicing, which results in a circular molecule with a back-spliced junction (BSJ). Black boxes highlight the BSJ in the circRNA isoforms. **C.** CircRNAs can be detected with RT-qPCR using a BSJ-specific primer pair. The primer pair can only bind in a divergent manner (facing away from each other) to linear RNA, where no amplification will be possible, yet binds the circRNA in a convergent manner (facing towards each other), amplifying the BSJ sequence. **D.** Large-scale circRNA detection is typically performed using total RNA sequencing datasets and specialized computational tools. These tools identify BSJ-spanning reads, which map divergently (in reverse order) on the linear reference genome.

Although an *in vivo* function for most circRNAs remains unknown and functional analyses are typically restricted to *in vitro* experiments, some circRNAs have been linked to specific diseases, including cancer. CircRNAs have also been reported to be more stable than linear transcripts due to the absence of a free 5’ or 3’ end to be recognized by exonucleases (1). In line with this, a higher fraction of circRNA relative to linear RNA has been observed in a wide range of human biofluids, which makes them interesting biomarker candidates, with the potential to be used for minimally-invasive tests for diagnosis or response monitoring (2). Wang *et al.* reviewed 112 differentially expressed circRNAs in various biofluids from patients with different cancer types (3). Furthermore, 15 clinical trials incorporating circRNAs as disease biomarkers have been initiated (ClinicalTrials.gov, accessed on 20/10/2022).

Eukaryotic circRNAs are formed through a process called back-splicing, where the 5’ end of an RNA molecule forms a covalent bond with its own 3’ end, forming a circular molecule with a characteristic back-spliced junction (BSJ) sequence (Figure 1B) (1). CircRNAs comprise of one or multiple exons, and analogous to linear RNA, there is alternative splicing of circRNAs, where circRNAs with the same BSJ sequence may have a different exon (and/or intron) composition (1).

In a targeted manner, circRNAs can be quantified with RT-qPCR (reverse transcription quantitative polymerase chain reaction) using BSJ-spanning primer pairs to amplify the region flanking the BSJ (Figure 1C). These primer pairs are divergent (facing away from each other) when hybridizing to the linear host transcripts and can therefore only amplify the circRNA (4). However, false positives resulting from alignment ambiguity, repeat sequences, trans-splicing, or RT template-switching artifacts have been described (5, 6). In all these cases, a linear RNA molecule is formed with the same exon orientation and, therefore, the same sequence as the circRNA BSJ. To prevent false positive circRNA identifications, linear RNA is often digested with the exonuclease ribonuclease R (RNase R) followed by RT-qPCR. RNase R typically degrades linear RNA, whereas circRNAs are generally not affected. Of note, it has been suggested that long circRNAs may be somewhat sensitive to RNase R degradation and various challenges of validating circRNAs have been recognized (7, 8).

In general, high-throughput or exploratory circRNA detection is performed using bioinformatics approaches that analyze total RNA sequencing data. For this, the RNA sequencing reads are first mapped against a reference genome. The unmapped reads are subsequently used to identify BSJ-spanning reads that map divergently (in reverse order) on the linear genome (Figure 1D).

Over the last decade, numerous computational circRNA detection tools have been developed and tested. Whereas multiple sets of circRNA detection tools using a bioinformatics approach have been compared (often when a novel tool is published) (9–16), a systematic and comprehensive evaluation of many circRNA detection tools using an orthogonal validation method is still missing. In our benchmarking study, we aimed to evaluate all currently available circRNA detection tools with an orthogonal approach using RT-qPCR, RNase R, and amplicon sequencing (Figure 2A). Our study highlights that although the precision of the tools is generally excellent, their sensitivities are highly variable.

**Figure 2.**
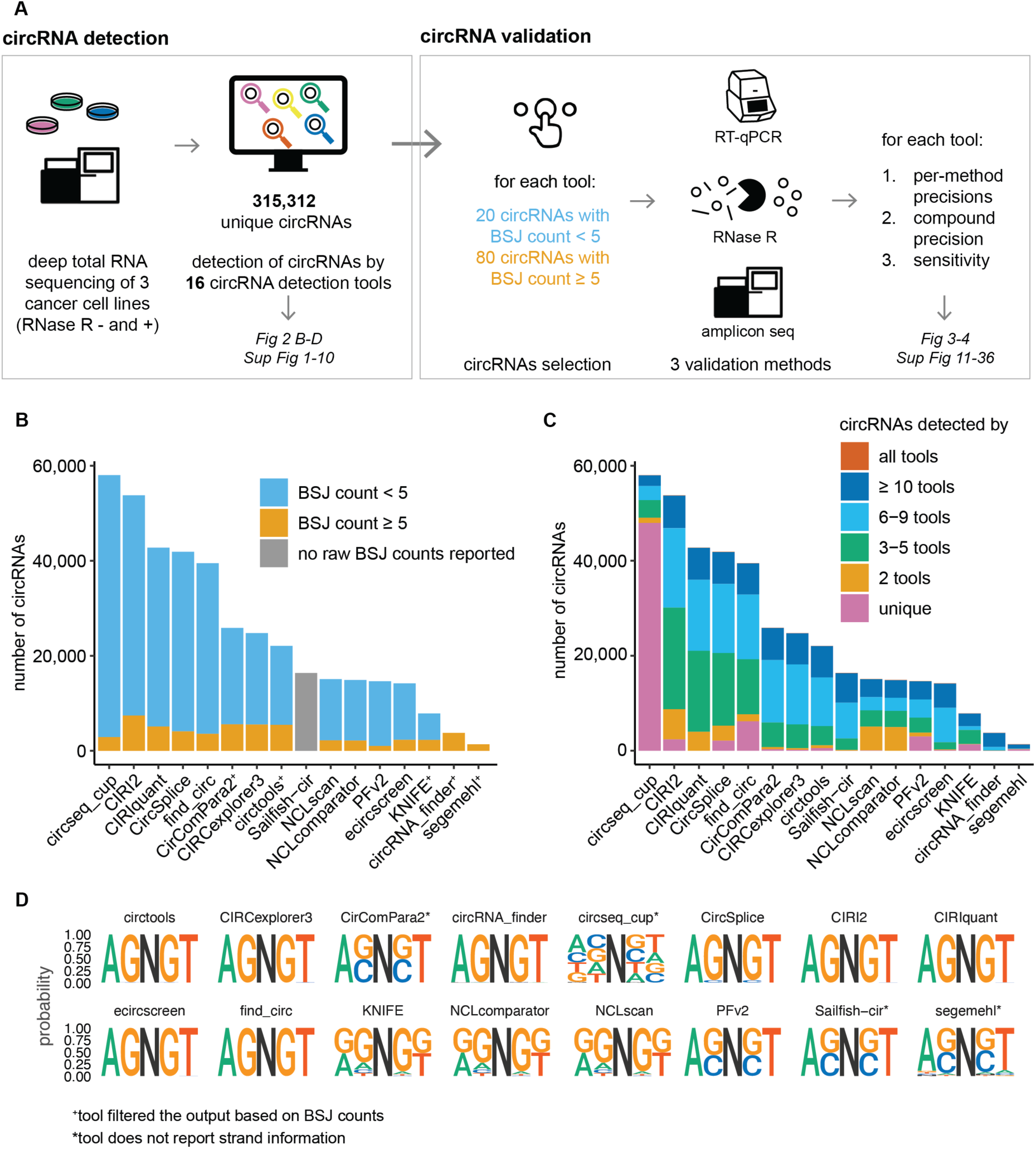
CircRNA detection tools predict a wide variety of circRNAs **A.** This study consists of a circRNA detection phase and a circRNA validation phase. For the former, 16 circRNA detection tools were used to predict circRNAs in three deeply sequenced cancer cell lines. For the latter, a set of circRNAs was selected per tool and validated using three orthogonal methods, generating tool-specific precision values for each method. This was also used to compute compound precision and both types of sensitivity values for each circRNA detection tool. **B.** The number of reported circRNAs differs greatly between tools (shown for HLF cells, similar results for the other cell lines are shown in Supplementary Figure 1). The tools are ordered according to the total number of predicted circRNAs. The vast majority of circRNAs are predicted with a BSJ count below 5 (in blue). Two tools, circRNA_finder, and segemehl, filtered their results to report only circRNAs with a BSJ count of at least 5 (in orange). **C.** The majority of circRNAs (49.9%) are detected by only one tool. Circseq_cup reports the largest set of unique circRNAs (shown for HLF cells, similar results for the other cell lines are shown in Supplementary Figure 4). A small set of 55 circRNAs is detected by all tools (column n_db in Supplementary Table 2). **D.** CircRNA splice sites differ among circRNA detection tools. Most commonly, the canonical AGNGT pattern is observed, with AG being the splice acceptor, N the circRNA, and GT the splice donor. Circseq_cup, CirComPara2, Sailfish-cir, and segemehl do not report strand information. To be able to retrieve a splicing sequence for the circRNAs from these tools, it was assumed the circRNA originated from the positive strand. This led to the ACNCT pattern (reverse complement of AGNGT), most probably from circRNAs that were assigned to the positive strand incorrectly. Lastly, there are some tools displaying the GGNGG pattern.

## Results

### CircRNA detection tools predict a wide variety of circRNAs

#### CircRNA detection tools differ in detection strategies and filtering

For this study, 16 different circRNA detection tools were included: CIRCexplorer3 (17), CirComPara2 (11), circRNA_finder (18), circseq_cup (19), CircSplice (20), circtools (21), CIRI2 (22), CIRIquant (23), ecircscreen (unpublished tool), find_circ (24), KNIFE (15), NCLscan (25), NCLcomparator (26), PFv2 (27), Sailfish-cir (28), and segemehl (29) (Table 1, Supplementary Table 1). CircRNA detection tools differ in their circRNA detection approach (including strand assignment), reliance on linear annotation, and filtering methods. CircRNAs can be detected from RNA sequencing data using the pseudo-reference-based approach (also called the candidate-based approach) or the fragmented-based approach (also called the segmented-read-based approach) (12, 14). The former approach uses a reference list of potential BSJ sequences, often based on all possible combinations of known annotated exons within a gene. This approach is therefore limited to species with annotated genomes and to previously annotated genes and will only detect circRNAs that use the same splicing sites as the linear RNAs. The latter approach splits unmapped sequencing reads into shorter sequences and remaps these against the reference genome. Lastly, integrative tools, such as CirComPara2 and ecircscreen, combine the results of multiple tools.

**Table 1.** CircRNA detection tools with their circRNA detection approach, strand assignment approach, reliance on linear annotation, and filtering approach. ^x^ Integrative tools combine the results of multiple circRNA detection tools. This includes CirComPara2 (combining CIRCexplorer2 v2.3.8 (on any of BWA v0.7.15, TopHat2 v2.1.0, STAR v2.6.1e, and Segemehl v0.3.4), CIRI2 v2.0.6, DCC v0.4.8, and find_circ v1.2, and the filtering all circRNAs detected by at least two methods) and ecircscreen (combining CIRI2 v2.0.6, circRNA_finder v1.2, PFv2 v2.0.0, find_circ v1.2, and CIRCexplorer v1.1.10, and then filtering all circRNA detected by at least three methods). ^+^ Some tools did not report strand information for this study, but the (updated) circRNA tool might report circRNA strand information. * The BSJ count, and minimum and maximum circRNA length filters, are the filters used for this specific study. The user can choose these parameters freely. Of note, the minimum and maximum length filters are based on the estimated circRNA length with introns, calculated by subtracting the start position from the end position of the BSJ. ^-^ Inferred based on publication and available code.

| **tool** | **approach** | **circRNAs detected in ...** | **strand assignment\*** | **splicing** | **BSJ count filter\*** | **min circRNA length\*** | **max circRNA length\*** |
| --- | --- | --- | --- | --- | --- | --- | --- |
| CIRCexplorer3 | segmented read-based | entire genome | based on linear annotation | AGNGT | none | none | none |
| CirComPara2 | integrative<sup>x</sup> | entire genome | no strand reported | AGNGT, ACNCT | ≥ 2 | 299 | 2,304,996 |
| circRNA_finder | segmented read-based | entire genome | based on mapping to genome | AGNGT | ≥ 5 | 200 | 100,000 |
| circseq_cup | based on segemehl, with full-length circRNA assembly | entire genome | no strand reported | non-canonical | none | none | 5,000 |
| CircSplice | segmented read-based | known splice sites | based on linear annotation | AGNGT, ACNCT | none | 78 | none |
| circtools | segmented read-based | entire genome | based on mapping to genome | AGNGT | ≥ 2 in ≥ 2 samples | 31 | 1,000,000 |
| CIRI2 | segmented read-based | entire genome | based on GT-AG splice sites | AGNGT | none | 135 | 200,000 |
| CIRIquant | based on CIRI2, with improved quantification | entire genome | based on GT-AG splice sites | AGNGT | none | 135 | 200,000 |
| ecircscreen | integrative<sup>x</sup> | entire genome | based on consensus from tools | AGNGT | none | none | none |
| find_circ | segmented read-based | entire genome<sup>-</sup> | based on mapping to genome<sup>-</sup> | AGNGT | none | none | 100,000 |
| KNIFE | candidate-based | entire genome | based on linear annotation | non-canonical | ≥ 2 | none | 1,000,000 |
| NCLscan | candidate-based | known splice sites | based on linear annotation | non-canonical | none | 100 | none |
| NCLcomparator | filtered results of NCLscan | known splice sites | based on linear annotation | non-canonical | none | 100 | none |
| PFv2 | segmented read-based | entire genome | based on mapping to genome | AGNGT, ACNCT | none | 50 | 1,000,000 |
| Sailfish-cir | based on CIRI2 v2.0.6 | entire genome | no strand reported | AGNGT, ACNCT | no BSJ counts reported | 135 | 200,000 |
| segemehl | segmented read-based | entire genome | no strand reported | non-canonical | ≥ 5 | none | 200,000 |

#### The number of detected circRNAs differs greatly between tools

A total of 315,312 unique circRNA predictions (corresponding to 1,137,099 unique circRNA/strand/tool/sample tuples) were detected using 16 different tools based on deeply sequenced total RNA from three human cancer cell lines (Supplementary Table 2, because of large file size, only available on https://github.com/OncoRNALab/circRNA_benchmarking). The circRNA detection tools were run by their developers (details in Methods and Supplementary Notes). There is a striking almost 40-fold difference between the tool with the highest number of predicted circRNAs (circseq_cup with 58,032 circRNAs) and the tool with the lowest number of predicted circRNAs (segemehl with 1,372 circRNAs) for one of the cell lines (Figure 2B shows results for HLF cells, similar results for the other cell lines are shown in Supplementary Figure 1).

#### Most circRNAs are characterized by low BSJ counts

CircRNA abundance is reflected by the BSJ count, which is the number of reads uniquely assigned to a given circRNA. The majority of circRNAs (86.6%) are detected with a BSJ count below 5 (Figure 2B), with only 46.1% of the detected circRNAs being observed with at least 2 BSJ counts (detailed distribution in Supplementary Figure 2). To increase confidence, circRNA_finder and segemehl filtered their results to report only circRNAs with a BSJ count of at least 5, and CirComPara2 and KNIFE filtered for circRNAs with a BSJ count of at least 2. Circtools filtered circRNAs with at least 2 counts in at least 2 samples. Of note, Sailfish-cir does not report raw BSJ counts, but transcripts per million (TPM) instead. The similarity of circRNA BSJ counts between tool pairs is reasonable, according to correlation analysis (linear model with median r^2^ = 0.71, median slope = 0.70, p-value < 0.001, Supplementary Figure 3).

#### CircRNA detection tools predict different sets of circRNAs

Half of all circRNAs in this study (49.9%) are reported only by one tool, which is largely due to circseq_cup’s high number of uniquely predicted circRNAs (Figure 2C for HLF cells, similar results for the other cell lines are shown in Supplementary Figure 4). The overlap of circRNA predictions among different tools is visualized in a heat map for each cell line in Supplementary Figure 5. Out of 16 circRNA detection tools, 8 exclusively report circRNAs flanked by canonical splice sites (with an AGNGT pattern, where AG is the splice acceptor, N represents the circRNA sequence in between, and GT is the splice donor) (Figure 2D). CirComPara2, circseq_cup, Sailfish-cir, and segemehl do not report circRNA strand orientation, explaining most of the ACNCT patterns. Two-thirds of all predicted circRNAs in this study (68.5%) are novel compared to a set of previously reported circRNAs extracted from 13 published circRNA databases (Circ2Disease, circad, CircAtlas, circbank, circBase, CIRCpediav2, CircR2disease, CircRiC, circRNADb, CSCD, exoRBase, MiOncoCirc, and TSCD) (Supplementary Figure 6) (30). Of note, approximately half of these novel circRNA candidates originate solely from circseq_cup. Looking at the tools individually, circseq_cup, KNIFE, NCLscan, and NCLcomparator report a higher number of novel circRNAs (87.8%, 53.9%, 53.4%, 53.3%, respectively) compared to the other tools (median 19.7%, interquartile range (IQR) 4.9-34.8%). Tools were further compared based on the predicted circRNA length, strand information, correspondence to linear annotation, and predicted exon composition (Supplementary Data 1-4, Supplementary Figure 7-10). No notable differences were observed among tools, except for CIRI2 and PFv2 having a higher number of circRNAs for which no canonical linear annotation match was found compared to the other circRNA detection tools. Across all tools, 53.7% of circRNAs uniquely match one canonical linear transcript, 10.3% match more than one canonical transcript, and 35.9% do not match any canonical transcript. CircRNAs were found for 17,461 different human genes, demonstrating the pervasive nature of back-splicing.

### CircRNA validation with empirical methods

#### CircRNA primer design inherently introduces a selection bias

Based on previous experiments (Supplementary Data 5, Supplementary Figure 11), for each tool, we aimed to select 80 random *high-abundance* circRNAs with a BSJ count of at least 5 and 20 random *low-abundance* circRNAs with a BSJ count below 5. Of note, circRNA primer design inherently introduces a bias (Supplementary Data 6, Supplementary Figure 12). A selection of 1,560 circRNAs was obtained (Supplementary Table 3, BSJ count distribution in Supplementary Figure 13). As some circRNAs were selected more than once (by chance, for different tools or in different cell lines), the total number of unique circRNAs/sample pairs was 1,516, from here on termed ‘selected circRNAs’ (detailed description in Supplementary Figure 14).

#### High BSJ detection precision using RT-qPCR validation

Of the 1,516 selected circRNAs, 1,479 (97.6%) could be validated with RT-qPCR, *i.e.*, the primer pair flanking the BSJ site resulted in a detectable amplicon. For the *low-abundance* circRNAs there is some variation in the tool-specific precision values (median 95.0%, range 80.0-100%), which is expected. *High-abundance* circRNAs have high RT-qPCR precision values for most tools (median 98.8%, range 90.0-100%) (Figure 3A; the cumulative plot of the RT-qPCR precision in function of the BSJ count is shown in Supplementary Figure 15). It is important to note that RT-qPCR-based validation is the net result of a successful primer pair and the actual presence of a sufficiently abundant circRNA in the amount of RNA tested.

**Figure 3.**
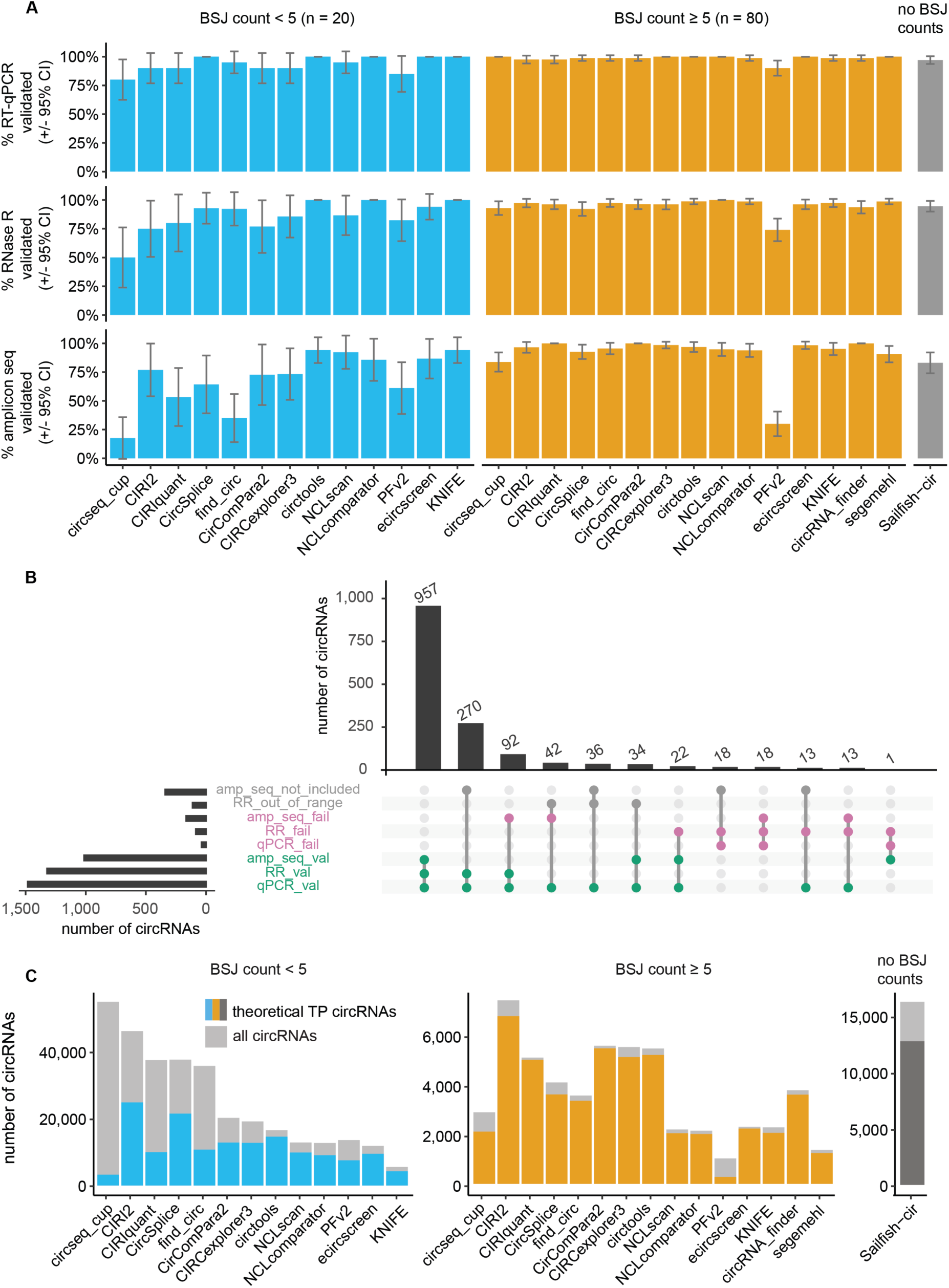
The precision of circRNA detection tools is generally high and similar, whereas tools largely differ with respect to the number of predicted circRNAs. The plots are separated based on circRNA BSJ count below 5 (*low-abundance*, in blue) or a BSJ count of at least 5 (*high-abundance*, in orange). Sailfish-cir reports TPM (transcripts per million) values instead of BSJ counts, and is therefore depicted separately. **A.** CircRNAs were validated using three different techniques: RT-qPCR detection, resistance to degradation by RNase R, and amplicon sequencing. *Low-abundance* circRNAs are in general more difficult to validate. Of note, the precision values for *low-abundance* circRNAs are based on a limited set of circRNAs. *High-abundance* circRNAs have good precision values for most tools and most validation methods. The error bars represent the 95% confidence intervals (CI). **B.** The vast majority of circRNAs obtain the same verdict based on the three different validation methods. However, some circRNAs have conflicting results. For example, there are 13 circRNAs that are detectable by RT-qPCR but also are degraded upon RNase R treatment and for which the primers seem to amplify the wrong product. **C.** The compound precision value is used to compute the theoretical number of true positive circRNAs by multiplying it with the original number of circRNAs detected by that tool (*i.e.,* the extrapolated sensitivity) (shown for HLF, similar results for the other cell lines are shown in Supplementary Figure 25).

#### One in sixteen predicted circRNAs fail validation upon RNase R treatment

RNase R was used as a second, more stringent validation approach. RNase R selectively degrades linear transcripts, ensuring the RT-qPCR primers amplify a circular molecule. For 112 out of 1,516 selected circRNAs (7.4%), RNase R treatment could not be evaluated, as their abundance in the untreated sample was too low, leaving no room to confirm RNase R degradation in the event of a false positive circRNA (hence labeled as NAs). In the remaining set of 1404 predicted circRNAs, 1319 circRNAs (93.9%) could be successfully validated using RT-qPCR on RNase R treated RNA. For most tools, high RNase R precision values were observed for *high-abundance* circRNAs (median 96.3%, range 74.0-100%). PFv2 displays the lowest precision (74.0%). For *low-abundance* circRNAs, lower precision values were observed (median 86.7%, range 50.0-100%) (Figure 3A; the cumulative plot of the RNase R precision in function of the BSJ count is shown in Supplementary Figure 15). Of note, the number of circRNAs per tool in this bin is lower than the original 20 that were selected, as more circRNAs were excluded due to too low abundancy (resulting in only 10-18 circRNAs per tool, with a median of 14 circRNAs). A comparison with matched RNase R treated and untreated sequencing data is described in Supplementary Data 7 and shows that the RNase R precision calculated from sequencing results is mostly high and similar among tools, with PFv2 having the lowest precision value (Supplementary Figures 16-19, Supplementary Tables 4 and 5, because of large file size, only available on https://github.com/OncoRNALab/circRNA_benchmarking).

#### Amplicon sequencing is the most stringent orthogonal validation method

The RT-qPCR amplicons of the untreated RNA were sequenced for further validation of the circRNAs. A random subset of circRNAs (337/1,516, 22.2%) was not included in the amplicon sequencing experiment (hence labeled as NAs). For the remaining 1,179 circRNAs, 1,014 circRNAs (86.0%) could be readily validated with amplicon sequencing, i.e., the majority of reads aligned to the expected BSJ sequence. Most tools have similar amplicon sequencing precision values for *high-abundance* circRNAs (median 95.5%, range 30.0-100%), with PFv2 displaying a very low (30.0%) amplicon sequencing precision value. Of note, as PFv2 was developed to retain repeat sequences, it is expected to result in more false positives. The most obvious are caused by linear read-through between exons in neighboring tandemly repeated gene clusters/interspersed repeats, and these tend to be abundant. For *low-abundance* circRNAs, performance is more diverse, with generally lower amplicon sequencing precision (median 73.3%, range 17.6-94.1%) (Figure 3A; the cumulative plot of the amplicon sequencing on-target amplification rate and the cumulative plot of the amplicon sequencing precision in function of the BSJ count are shown in Supplementary Figures 20 and 15, respectively).

#### Different validation methods should be used concurrently to compensate for their inherent limitations

Although the three validation strategies were used independently, it is interesting to evaluate to what extent they support each other (Figure 3B; simple overview in Supplementary Figure 21). Considering 1,103 circRNAs for which all three validation results are available, 957 circRNAs (86.8%) pass all validation methods, 128 circRNAs (11.6%) fail one or two of the validation methods, and 18 circRNAs (1.6%) fail all three validation methods. These observations show that orthogonal validation with more than one empirical approach is important. It is beyond the scope of this study to investigate why there are some discrepancies among the validation results (some hypotheses are considered in the Discussion). First, they are rare (for most circRNAs, the different methods completely agree). Second, the same methods are used to compare the tools, whereby no tool should be favored over the other.

The three orthogonal validation methods were combined to label each circRNA as a true or false positive result and the compound precision was calculated for each tool. Similar to the separate precision values, the compound precision is high and similar for most tools when looking at *high-abundance* circRNAs (median 93.1%, range 27.1-98.3%, IQR 90.5-95.3%), and lower and more variable for *low-abundance* circRNAs (median 63.6%, range 5.9-88.2%, IQR 53.8-76.5%) (Supplementary Figures 15 and 22).

#### CircRNA detection tools differ greatly in sensitivity

Tool sensitivity was evaluated using two different methods. First, sensitivity was calculated based on the total number of true positive circRNAs (n = 957) (Supplementary Figures 23 and 24). Of note, this sensitivity metric should be used with caution as it is based on a biased set of circRNAs (see Methods). Second, the theoretical number of true positive circRNAs for each tool was computed by multiplying the total number of detected circRNAs and the compound precision (*i.e.,* the extrapolated sensitivity) (Figure 3C for HLF cells, similar results for the other cell lines are shown in Supplementary Figure 25). There is a significant positive correlation between both sensitivity values (Spearman rank correlation of 0.84 with p-value < 0.001, S = 58, for *low-abundance* circRNAs, and a Spearman rank correlation of 0.80 with p-value < 0.001, S = 113, for *high-abundance* circRNAs). Both methods show great variability in tool sensitivity, with a median sensitivity of 75.1% (range 29.7-87.1%) for *low-abundance* circRNAs and 65.7% (range 18.7-87.4%) for *high-abundance* circRNAs. To visualize the relationship between sensitivity and compound precision, a precision-recall (sensitivity) dot plot for all tools is shown in Supplementary Figure 26.

All metrics described above ((compound) precisions and sensitivity) and the tool ranking for each metric are available in Supplementary Table 6. The user can easily filter and order the circRNA detection tools based on their preferences. Reproducibility evaluations were performed and are described in Supplementary Data 8-10 (Supplementary Figures 27-31).

#### Evaluation of circRNA detection tool precision in function of circRNA annotation

To compare precision values in function of circRNA annotation, we restrict the analyses to *high-abundance* circRNAs with information for all validation techniques. Furthermore, a strict validation definition was used, where all circRNAs failing for at least one technique were classified as unvalidated. CircRNAs previously described in databases have higher chances of getting validated (Chi-squared = 181.0, degrees of freedom (df) = 1, p-value < 0.001, odds ratio (OR) = 13.1). Nevertheless, false positive circRNAs according to our data are still present in multiple published databases (Supplementary Figure 32). For example, false positive circRNA chr6:47526627-47554766 (hg38, 0-based) is present in CircAtlas (as hsa-CD2AP_0048) and in exoRBase (as exo_circ_65199). A difference in validation rate in function of the splicing pattern was observed, with better validation of circRNAs surrounded by canonical splice sites (Chi-squared = 45.4, df = 1, p-value < 0.001, OR = 5.0). Similarly, circRNAs that originate from a region with an annotated linear transcript have higher validation rates (Chi-squared = 185.8, df = 1, p-value < 0.001, OR = 17.1). Surprisingly, single-exon circRNAs displayed significantly lower validation rates than multi-exon circRNAs (Chi-squared = 20.0, df = 1, p-value < 0.001, OR = 3.8). Lastly, while tools with a ‘candidate-based’ approach seem more precise than tools using the ‘segmented read-based’ approach (Chi-squared = 9.4, df = 1, p-value = 0.0022, OR = 2.6), we cannot be sure that these results are not confounded by other algorithmic differences.

#### Evaluation of circRNA detection tool sensitivity in function of circRNA annotation

There is a significantly higher sensitivity for tools reporting circRNAs surrounded by canonical splice sites, resulting in a median difference in sensitivity of 38.5% (two-sided Mann-Whitney U = 55, p-value = 0.0022, large effect size of 0.78, 95% CI [0.56 - 0.85], n1 = 11, n2 = 5, only *high-abundance* circRNAs). However, no link could be found between sensitivity and tool approach, use of linear annotation, strand annotation method, or BSJ count filtering.

### Evaluation of tool combinations to improve performance

For the combination of two or more tools, both the intersection and the union have been proposed (11) (Supplementary Tables 7 and 8). While not evaluated here, the increased time and resource consumption should also be taken into account when considering the use of multiple tools. A list of the top-performing combinations is available in Supplementary Table 9 and can be used as a reference.

#### A circRNA predicted by two detection tools is not necessarily a true positive result

Figure 4A shows that circRNAs uniquely detected by a single tool generally have lower precision values. In line with this, circRNAs detected by at least two tools have a higher chance of getting validated (Chi-squared = 333.1, df = 1, p-value < 0.001, OR = 53.8). On the other hand, out of 1,380 unique circRNAs detected by at least two tools, 7 circRNAs (0.5%) failed all three validation methods, and 137 (9.9%) failed at least one of the validation methods, illustrating that the practice of using the intersection is not a guarantee to avoid false positive results.

**Figure 4.**
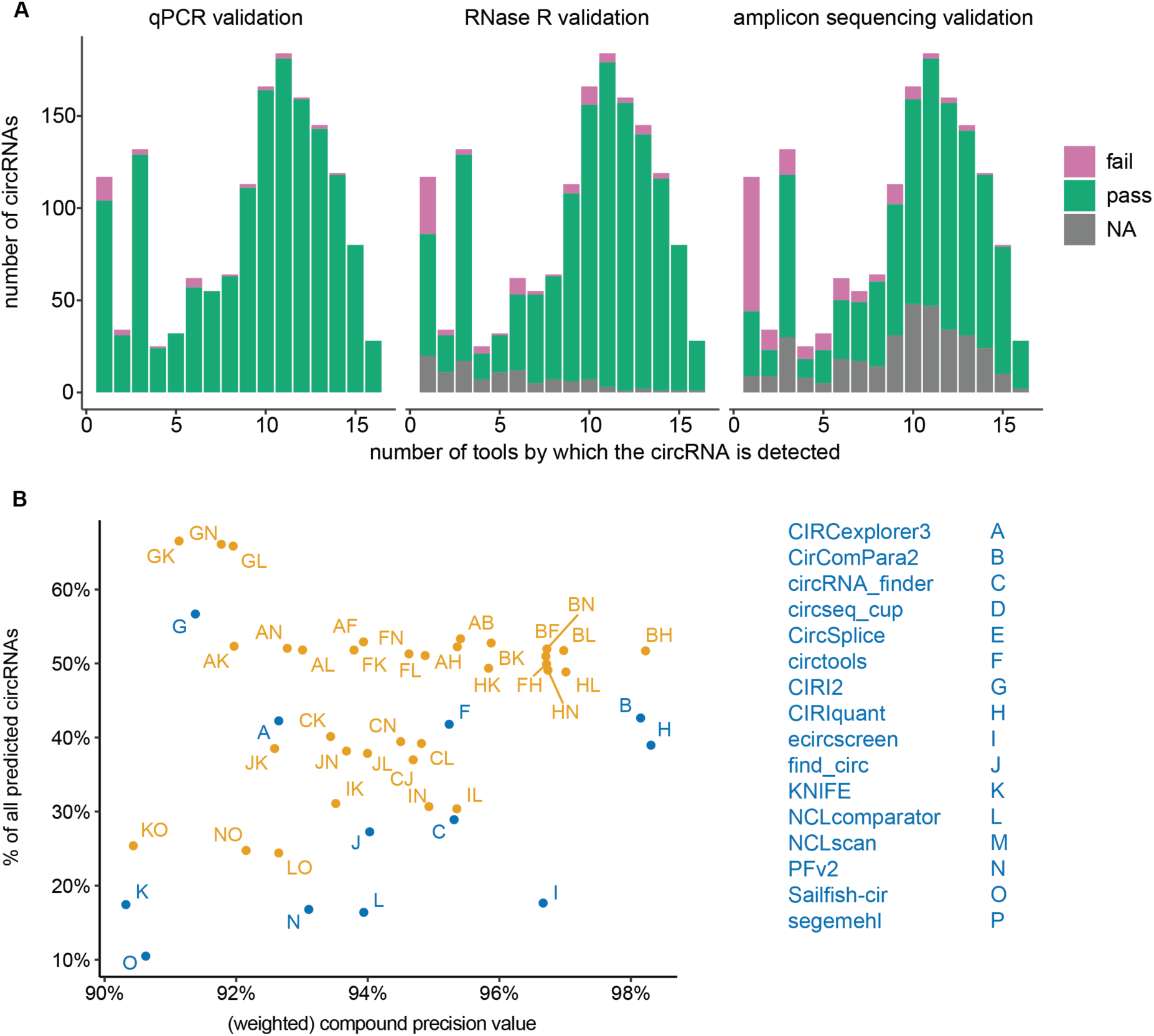
The intersection or union of two circRNA detection tools decreases the number of false positives, or increases the overall number of detected circRNAs, respectively. **A.** CircRNAs detected by multiple tools generally have higher precision values. However, the often-used practice of using the intersection of two tools is not necessarily a guarantee for avoiding false positive results. **B.** By considering the union of two circRNA detection tools, the number of circRNAs can be significantly increased, whilst keeping the number of false positive predictions low (shown for the HLF cell line, similar results for the other two cell lines are shown in Supplementary Figure 33). For the y-axis, the percentage of detected circRNAs is calculated by dividing the number of circRNA detected by that tool combination by the total number of predicted circRNAs for that sample taking the union of all tools (13,087 circRNAs for the HLF sample). For this analysis, the compound precision value of *high-abundance* circRNAs was used. Some circRNA detection tools are integrative and combine the results of multiple other tools. It is therefore assumed an integrative tool would have large similarities with its integrated tools. However, a difference in tool version and filtering can still give a different set of circRNAs. For example, CirComPara2 is an integrative tool that combines CIRCexplorer2, CIRI2, DCC, and find_circ, nevertheless, the combination of CirComPara2 and CIRCexplorer3 still gives a significant increase in detected circRNAs (corresponding to 10% of all circRNA predictions for that cell line).

#### The union of highly-precise circRNA detection tools substantially increases the number of true positive circRNAs

To maximize detection sensitivity and maintain precision, we evaluated the union of pairs or triples of circRNA detection tools. Generating all possible combinations of the better tools with individual compound precision ≥ 90% for *high-abundance* circRNAs (n = 12 tools) consistently results in higher detection sensitivity while maintaining a high weighted precision value. The median increase in the number of detected circRNAs is 37.0% (IQR 16.5-129.7%) and 79.6% (IQR 33.1-215.7%) for combinations of 2 or 3 tools, respectively. In other words, when combining very precise tools, the number of false positives does not counteract the gain in additional true positives. A subset of tool combinations with an increase of at least 1,000 circRNAs is shown in Figure 4B (shown for HLF cells, similar results for the other cell lines are shown in Supplementary Figure 33; the combo of three tools is shown in Supplementary Figure 34). One obvious consideration when selecting two different tools is their circRNA detection approach, their reliance on linear annotation, and their filtering methods. For example, when combining two tools with a different detection approach (pseudo-reference-based and fragmented-based approach), the median increase in the number of detected circRNAs is 61.1%, compared to 35.4% for two tools with the same detection approach (two-sided Mann-Whitney U = 58554.5, p-value < 0.001, small effect size of 0.16, 95% CI [0.08 - 0.23], n1 = 336, n2 = 294). Similarly, when combining two tools with the same splice site settings (both canonical or both non-canonical), the median increase in the number of detected circRNAs (32.6%) is significantly smaller compared to combining two tools with different splice site settings (one canonical and one non-canonical) (76.2%) (two-sided Mann-Whitney U = 65356.5, p-value < 0.001, small effect size of 0.28, 95% CI [0.21 - 0.35], n1 = 300, n2 = 330). A similar analysis for the combination of tools that either nor not rely on linear annotation was not significant.

## Discussion

Multimodal orthogonal validation of bioinformatics tools that predict circular RNAs from total RNA sequencing data is currently lacking. Hence, their precision and sensitivities are unknown and scientific data is confounded with false positive and false negative predictions. To accommodate this lacune, we set up a large-scale international collaborative circRNA detection tool benchmarking study (Figure 2A). First, a deeply sequenced total RNA sequencing dataset was processed by the developers of 16 different circRNA detection tools. Next, 3 empirical validation strategies were used to evaluate a random selection of 1,560 circRNAs representing each tool: 1) RT-qPCR to determine if the candidate circRNA BSJ (back-spliced junction) sequence was detectable; 2) RNase R treatment to confirm that the detected RNA was most likely circular and not linear; and 3) amplicon sequencing to confirm the circRNA BSJ sequence. Of note, both circRNA RT-qPCR and RNAse R validation protocols were extensively validated (4, 31).

The precision values are similarly high among tools (Figure 3A), especially when considering the subset of *high-abundance* circRNAs (with a BSJ count ≥ 5). In contrast, the number of predicted circRNAs and the sensitivity varies greatly among tools, in line with previous studies based on simulated data (11, 13, 14) (Supplementary Data 11, Supplementary Figures 35 and 36). The striking differences in sensitivity are in part dependent on the operator applying BSJ count filters.

The three validation methods each have their own strengths and biases, with conflicting results for several circRNAs (Figure 3B, discussed in detail in Supplementary Discussion 1). In total, 22 circRNAs are validated with qPCR and amplicon sequencing but are degraded by RNase R for at least 87.5% (*i.e.*, a difference of 3 cycles). A possible explanation could be that some *bona fide* circRNAs are susceptible to RNase R degradation (8) or that the primers amplify a mixture of circular and linear RNA. Another subset of 92 circRNAs pass RT-qPCR validation and RNase R validation but fail amplicon sequencing. These could be (repetitive) RNAs resistant to RNAse R due to secondary structure, either internal or through base pairing with orthologs (8). These examples underscore the importance of using different validation methods to compensate for their intrinsic limitations and to increase the validation status confidence (as previously suggested in (8)).

Although long-read sequencing has been implemented to study full-length circRNAs (32–35), the bulk of currently available data remains short-read sequencing. Therefore, this benchmarking study evaluated circRNA detection tools for short-read sequencing data, which typically report circRNAs by their BSJ position (chr, start, end, strand). However, it remains unknown if the detected BSJ corresponds to one circRNA, or multiple alternatively spliced circRNAs with different exon/intron compositions. Henceforth, the prediction precision values reported here might be influenced by more than one circRNA with the same BSJ. As this study is focused on circRNA detection in short-read sequencing data, the internal circRNA composition was not evaluated. Furthermore, no distinction can be made between circRNAs on the positive strand or negative strand using RT-qPCR and amplicon sequencing (9.4% of circRNAs were reported to originate from different strands according to different tools).

Based on a pilot study (Supplementary Data 5, Supplementary Figure 11), a cut-off was set at BSJ count 5, as circRNAs under this cut-off approached the qPCR limit to reliably detect RNase R based degradation of falsely predicted circRNAs. While very deep sequencing of a large RNA input amount was performed, it is beyond the scope of this study to evaluate if the BSJ count should be reconsidered in function of sequencing depth. However, as the majority of predicted circRNAs have a BSJ count below 5, we decided to include at least a subset of these *low-abundance* circRNAs to calculate the corresponding prediction precision. It is no surprise that the precision values for *low-abundance* circRNAs are significantly lower compared to *high-abundance* circRNAs (Chi-squared = 76.7, df = 1, p-value < 0.001, OR = 3.8). This difference is likely due to the detection limits of the applied validation strategies in conjunction with the sampling bias of *low-abundance* analytes, and not due to inherently more false positive predictions for circRNAs with a lower count. Of note, it can be presumed that weakly expressed circRNAs are less relevant for both functional studies and biomarker research.

Focusing on *high-abundance* circRNAs, interesting links were found between circRNA annotation and validation rates. As such, circRNAs had higher validation rates when they were detected by multiple tools, when they were previously reported in a circRNA database, when they were surrounded by canonical splice sites, and when they originate from a region with an annotated linear transcript. CircRNA detection tools with a ‘candidate-based’ approach are more precise than tools using the ‘segmented read-based’ approach, which is in line with the higher validation likelihood of circRNAs originating from known linear genes and surrounded by canonical splice sites.

Based on our study, we compiled a list of recommendations for circRNA detection and validation, and for the future development of circRNA detection tools and their performance evaluation (Table 2). Ideally, publicly available (spike-in) reference material (consisting of known synthetic circRNAs) should be used to benchmark existing and novel circRNA detection tools. However, such reference material is currently not available. As the main goal of this study was to perform a neutral assessment of circRNA detection tool sensitivity and precision, the developers of the tools were asked to run the tools themselves. Therefore, execution time, memory usage, and ease of use could not be compared and were not assessed here.

**Table 2.** CircRNA research recommendations

|  |  |
| --- | --- |
| **circRNA detection** | 1. An orthogonal validation method must be used to validate a predicted circRNA; qPCR validation on its own is not sufficient, at least qPCR + RNase R treatment or preferably qPCR + amplicon sequencing should be used.<br>2. Filtering based on a minimum BSJ count is recommended to increase the likelihood of successful empirical validation. |
| **circRNA validation** | 3. For a precision-focused approach, the *intersection* of two tools with a high individual precision (for example $\geq 90\%$) should be used.<br>4. For a sensitivity-focused approach, the *union* of two tools with a high individual precision (for example $\geq 90\%$) should be used.<br>5. The choice of tools to be combined may be informed based on the tools' underlying principles (circRNA detection approach, reliance on linear annotation and canonical splicing, and filtering). |
| **circRNA tool development** | 6. Tools should report the originating strand information, the BSJ count evidence, and the chromosomal start and end position of the BSJ. |
| **circRNA tool validation** | 7. For evaluation of sensitivity, novel and updated tools are encouraged to use the empirically validated set of 957 true positive circRNAs.<br>8. For evaluation of precision, a random set of 100 predicted circRNAs should be validated with empirical methods. |

Furthermore, this study resulted in a circRNA resource containing > 315,000 circRNAs detected in three human cancer cell lines from different tissue origins and provides validation results for 1500 circRNAs that can be used as a reference for the development of new or improved circRNA detection tools. Finally, our study can also serve as an example framework for empirical validation of benchmarking results from other bioinformatics tools in the future.

## Online methods

### Study set-up

As this study includes executing and evaluating circRNA detection tools, the co-authors can be divided into two groups 1) a first independent group (with no circRNA detection tool of their own) which initiated and designed the study and performed all the wet-lab work and data analysis (the validation co-author group), and 2) a second group of tool developer co-authors, which detected circRNAs using their own circRNA detection tools according to their expertise (the circRNA prediction co-author group) (details in Author Contribution section). During the study, meetings and emails were used to share the results (initially in a blinded manner) and discuss the final manuscript with the circRNA prediction co-authors.

### Cell culture

Three cancer cell lines from different cellular origin were randomly chosen as biological replicates. Ethical approval was obtained for this study (#EC014-202, Ghent University Hospital) and the cell lines were purchased from the JCRB Cell Bank (HLF and NCI-H23) or ECCAC (SW480). SW480 cells were cultured at 37 °C, 0% CO_2_ in Leibovitz’s L-15 medium (#31415-029, ThermoFisher). HLF cells and NCI-H23 cells were cultured at 37 °C, 5% CO_2_ in DMEM, low glucose, GlutaMAX Supplement, pyruvate (#21,885,025, ThermoFisher) and RPMI 1640 Medium, HEPES (#52,400,041, ThermoFisher), respectively. 10% fetal bovine serum (FBS) (#F7524, Sigma) and 1% penicillin-streptomycin (10,000 U/mL) (#15,140,122, ThermoFisher) were added to all three media.

### RNA isolation

RNA was isolated from the cells using the miRNeasy Mini kit (#217004, Qiagen) according to the manufacturer’s instructions, including the optional on-column DNase treatment (#79254, Qiagen). For each cell line, a sufficient number of cells was cultured to be able to harvest a minimum of 330 µg RNA. The RNA concentration was measured spectrophotometrically using a NanoDrop instrument and the RNA integrity was evaluated using a Fragment Analyzer. For each cell line, the RNA was pooled and aliquoted (1000 ng RNA in 100 µL nuclease-free water per aliquot) and stored at –80 °C, making a uniform RNA collection to use for all downstream experiments.

### RNase R treatment, library preparation, and sequencing

For each cell line, two aliquots of 1000 ng input RNA (in 10 µl nuclease-free water) were used. First, ribosomal RNA (rRNA) was removed with the NEBNext rRNA Depletion Kit (#E6350X, New England Biolabs), following the manufacturer’s instructions. Next, RNAse R treatment was performed according to our previously described protocol (4). In summary, one aliquot of each cell line was treated with RNase R (#RNR07250 (250 U), Lucigen), and one aliquot of each cell line was treated as a buffer control. This was followed by a clean-up step using Vivacon 500, 10,000 MWCO Hydrosart columns (VN01H02, Sartorius). Subsequently, the NEBNext Ultra II Directional RNA Library Prep Kit for Illumina (#E7760L, New England Biolabs) was used in combination with the NEBNext Multiplex Oligos for Illumina (#E7600S, New England Biolabs) to index and prepare the samples for sequencing. The library preparation protocol was adjusted to obtain relatively long insert sizes (average size of 636 nucleotides measured using a Fragment Analyzer): RNA fragmentation of 7.5 minutes; first-strand cDNA synthesis elongation step of 50 minutes instead of 15 minutes. The last bead clean-up step was performed twice to completely remove all indexes from the samples. Finally, the samples were pooled equimolarly and sequenced on a NovaSeq 6000 instrument using a NovaSeq 6000 S1 Reagent Kit v1.5 (300 cycles) (#20028317, Illumina), resulting in approximately 300 million paired-end 150-nucleotides reads per sample. Raw FASTQ files are stored in the Sequence Read Archive (PRJNA789110: SRX13414572 (untreated HLF), SRX13414573 (untreated NCI-H23), SRX13414574 (untreated SW480), SRX13414575 (RNase R treated HLF), SRX13414576 (RNase R treated NCI-H23), SRX13414577 (RNase R treated SW480)).

### CircRNA detection

In November 2020, a comprehensive list of all published circRNA detection tools was compiled, and all developers were invited to collaborate. Upon consent, they were asked to detect circRNAs using their own circRNA detection tool as they seemed fit for the data that was provided. The circRNA detection steps for each tool are detailed in the Supplementary Notes. Often, the default parameters were used as most of the methods included in our benchmarking underwent continuous development during the last several years and their parameters have been optimized for standard RNA sequencing data (as is the case in this study). We were unable to get into contact with the authors of find_circ (24) and decided to run this tool ourselves, as it is one of the most frequently cited and broadly used circRNA detection tools. Unfortunately, other well-performing tools (according to (10, 14)), such as MapSplice (36), could not be included. More recent tools, such as Circall (13) and CYCLeR (37), have been published after the validation experiments were performed, and are therefore not included.

After collecting all circRNA detection results, a uniform list of circRNAs defined by their BSJ position (chr, start, end, strand) and the BSJ count for each tool was compiled (Hg38, 0- based).

### CircRNA selection and primer design

Guided by a pilot experiment assessing circRNA RT-qPCR detectability in function of abundance and RNA input amount (Supplementary Data 5, Supplementary Figure 11), for each tool, 80 *high-abundance* circRNAs (with a BSJ count of at least 5), and 20 *low-abundance* circRNAs (with a BSJ count below 5) were selected (as two separate count bins). Primer pairs were designed using our primer design tool CIRCprimerXL (31). All primer sequences are available in Supplementary Table 3. If no primer pair could be designed for a given circRNA, a substitution was randomly selected from the complete dataset, considering the BSJ count bin. In total, 1,560 circRNA/tool/cell line tuples were selected. As some circRNAs were selected more than once (for different tools) the total number of unique circRNA/cell line pairs is 1,516, and the number of unique circRNAs (not taking into account the strand) is 1,457 (Supplementary Figure 14). Additionally, most of the selected circRNAs are detected by multiple tools (for which they were not selected). For the precision calculations, only the 20 + 80 selected circRNAs for a specific tool were used to evaluate that tool to keep the number of observations equal for each tool, even though more of its predicted circRNAs might have been validated. However, for the sensitivity calculations, the complete set of circRNAs had to be used (*vide infra*).

### RNase R and RT-qPCR

The RNA aliquots derived from the three cell lines were used for the circRNA RT-qPCR validation. A total of 1,080, 900, and 780 µl RNA (100 ng/µL) was required to validate 579, 500, and 437 circRNAs in HLF, NCI-H23, and SW480 cells, respectively. RNase R treatment was performed according to our previously reported protocol (4), adapted for this large-scale experiment. In summary, one RNA aliquot of a given cell line was treated with RNase R (#RNR07250 (250 U), Lucigen) and another was treated as a buffer control, for a total of 92 RNase R treated replicates and 92 buffer control replicates (2 * 36 for HLF, 2 * 30 for NCI-H23, and 2 * 26 for SW480 RNA). All volumes were doubled during the buffer and RNase R reaction (total reaction volume of 20 µL). This was followed by a clean-up step using Vivacon 500, 10,000 MWCO Hydrosart columns (#VN01H02, Sartorius). Next, the RT reaction was performed on the 184 separate replicates using the iScript Advanced cDNA Synthesis Kit (#172-5038, Bio-Rad), according to the manufacturer’s instructions. After RT, the cDNA was diluted 1:2 and an aliquot (2.5 µL) was further diluted 1:4 to evaluate the success of the RNase R reaction for each individual replicate. For this, ACTB and a known circRNA (chr1:117402185-117420649) previously described (4) (primer sequences available in Supplementary Table 10) were measured with qPCR using 2.5 µL 2x SsoAdvanced Universal SYBR Green Supermix (#172-5274, Bio-Rad), 0.5 µL forward and reverse primer (5 nM), and 2 µl cDNA per well, with qPCR duplicates. Once the RNase R treatment was successfully validated, all cDNA replicates were pooled per cell line and treatment condition. The cDNA was diluted 1:5 in 2× SsoAdvanced Universal SYBR Green Supermix (#172-5274, Bio-Rad). All 1,560 circRNA primer pairs were ordered from IDT in 96-well plates at a concentration of 100 µM in nuclease-free water. All primers were diluted 1:160 to obtain a 0.625 µM concentration. In each well of a qPCR plate, 2 µl diluted primers and 3 µl cDNA-master mix combination were added, resulting in an equivalent of 25 ng input RNA per qPCR reaction. Each assay (circRNA) was measured 4 times to include qPCR duplicates and to measure the abundance in both an RNase R untreated and treated sample, resulting in a total of more than 6000 qPCR reactions. A pipetting robot (EVO100, TECAN L) was used to dilute the primers and fill the qPCR plates. The qPCR reactions were run on a CFX384 instrument (Bio-Rad). Cq calling was done using the Bio-Rad CFX Manager (version 3.1), with the ’regression’ settings. The plates were stored at -20 °C prior to amplicon sequencing.

### Amplicon sequencing

After RT-qPCR, ∼80% of the circRNAs were randomly included for amplicon sequencing. To make the sequencing library, the amplicons were pooled by combining 2 µL of the PCR reaction from one of the untreated qPCR duplicates, per cell line. Next, the 3 samples were cleaned using Vivacon 500, 10,000 MWCO Hydrosart columns (#VN01H02, Sartorius). The PCR product pools were analyzed using a TapeStation 4150 (Agilent) and the concentration was measured using a Qubit fluorometer (ThermoFisher). Next, the three pools were diluted in nuclease-free water to obtain 50 µl samples with a concentration of 20 ng/µl. Finally, the samples were prepared for sequencing using the NEBNext Ultra II DNA Library Prep Kit for Illumina (#E7645S, New England Biolabs) and NEBNext Multiplex Oligos for Illumina (Dual Index Primers Set 1) (#E7600S, New England Biolabs). To retain all amplicons, no size selection was performed after adaptor ligation, and 1.0x AMPure XP beads (#A63881, Beckman Coulter) in a 1:1 sample:beads ratio was used instead. After library preparation, the samples were pooled equimolarly. The pool was sequenced on a NextSeq 500 instrument using a Mid Output Kit v2.5 (150 cycles) (#20024904, Illumina), resulting in approximately 25-30 million paired-end 75-nucleotides reads per library.

### Data analysis

Data analysis was mostly done using R (38) (version 4.2.1) in RStudio (39) (version 2022.07.1, build 554). The following R packages were used: tidyverse (version 1.3.2), conflicted (version 1.1.0), ggrepel (version 0.9.1), ggseqlogo (version 0.1), europepmc (version 0.4.1), gplots (version 3.1.3), ggpubr (version 0.4.0), quantreg (version 5.94), rstatix (version 0.7.0) and UpSetR (version 1.4.0). For sequencing data analyses, including circRNA detection and amplicon sequencing analysis, the Ghent University high-performance cluster was used. For this, Python3 (version 3.6.8) (40), Bowtie2 (version 2.3.4.1) (41), fastahack (version 1.0.0), SAMtools (version 1.11) (42), and BEDTools (version 2.30.0) (43) were used. The human reference transcriptome was downloaded as a GTF file from Ensembl (44). All data analysis scripts are available at https://github.com/OncoRNALab/circRNA_benchmarking.

#### Amplicon sequencing data analysis

For the amplicon sequencing data analysis, first, a custom Python script matches the primer sequences with the first 16-mer of each read (forward and reversed) and generates a separate FASTQ file per primer pair, containing all reads starting with that primer sequence. The FASTQ reads are then clipped to remove the primer sequences. Next, all FASTQ files are mapped against the reference genome (Ensembl version GRCh38.101) supplemented with the theoretical BSJ amplicon sequences using Bowtie2 with default settings. Lastly, the Bowtie2 BAM files are converted to counts using another custom Python script and the percentage on-target amplification was calculated for each primer pair.

#### Filtering and determination of orthogonal precision values and sensitivity

Several strategies to filter the data prior to precision and sensitivity calculations were explored. For RT-qPCR, a circRNA was considered validated when at least one of the untreated RNA samples had a Cq value above 10. Multiple variations of this threshold and a potential upper Cq threshold were evaluated. For RNase R validation, a subset of circRNAs with at least one untreated replicate with a Cq value below 32 was selected to ensure that the enzymatic degradation of a false positive circRNA could be measured. Multiple approaches to deal with these very low abundant circRNAs were compared and no significant differences were observed (data not shown). A circRNA was considered validated upon RNase R treatment if the difference in Cq between the untreated and treated RNA sample was equal to or less than 3 cycles, based on a previous study (4). As there were two qPCR replicates available for each (un)treated sample, the ‘best-case scenario’ was used to calculate the difference in Cq by subtracting the maximum untreated Cq replicate from the minimum treated Cq replicate. A circRNA with both untreated replicates having a Cq value above 32 was labeled as NA. For amplicon sequencing, a circRNA was considered validated if the primer pair was found in at least 1000 reads and if at least 50% of these reads matched the expected amplicon upon mapping with Bowtie2. For a random subset of circRNAs, unintentionally no amplicon sequencing was performed; these were labeled as NA. A detailed description of the choice of performance metrics is available in Supplementary Data 12 and 13. To calculate precision values per tool, BSJ count bin, and validation method, the number of circRNAs that passed the validation was divided by the total number of circRNAs that were not NA for that validation method. We also determined a compound precision value by considering both qPCR, RNase R treatment, and amplicon sequencing. For this, each circRNA was labeled as a true positive (*i.e.,* validated by all three methods), as a false positive (*i.e.*, not validated by at least one of the methods), or as NA (*i.e.,* not included in the amplicon sequencing run). Based on this summarizing label, compound precision values were computed for each tool and BSJ count bin. The number of theoretically true positive circRNAs was calculated by multiplying the total number of circRNAs predicted by that tool for that sample with the compound precision value (*i.e.,* the extrapolated sensitivity). The sensitivity was also calculated as the percentage of circRNAs each tool detected from the validated set of true positive circRNAs (*i.e.,* the circRNAs labeled as true positives over all three methods). This metric should be used with caution as it is based on a biased selection of circRNAs due to the overlap among tools (Supplementary Data 12). To calculate the sensitivity per BSJ count group, the median BSJ count of each circRNA was used (as most circRNAs are detected by multiple tools and have therefore multiple BSJ count values).

#### Annotation of circRNAs

To obtain the circRNA splice site information, the BSJ-flanking nucleotides were extracted from the reference genome using fastahack (Ensembl version GRCh38.104). To compare BSJ positions with known linear annotation, BEDtools intersect was used with a list of canonical transcripts from Ensembl with their positions based on the corresponding Ensembl GTF file (Ensembl version GRCh38.103). When a circRNA mapped to multiple isoforms, the annotation was labeled as ‘ambiguous’ and the circRNA was not taken into account for further annotation-based calculations and figures. The annotation was used to compute the length of each circRNA excluding introns, and the number of exons per circRNA. CircRNAs smaller than their host gene exon were labeled ‘single-exon’ circRNAs. For the length of each circRNA including introns, the BSJ start position was simply subtracted from the BSJ end position. Furthermore, for each circRNA, annotation was added to indicate if the BSJ start and end positions match known exon boundaries. When comparing predicted circRNAs to circRNAs previously described in databases, strand information was discarded.

#### Combination of tools

To compare the circRNA tools, the union and intersection of all circRNAs predicted by each tool pair and triple were calculated. A weighted precision value was calculated for each combination of tools as follows: ((perc_compound_val_1 * total_n_1) + (perc_ compound _val_2 * total_n_2)) / (total_n_1 + total_n_2). For this, strand information was discarded, as 4 out of 16 tools did not report circRNA strands and would therefore have been excluded. These calculations were performed for each cell line separately. To determine the correspondence among tools, the Jaccard distance was calculated and heatmap clusters were generated. The tools were compared based on the mere presence or absence of a circRNA. Also, for the calculation of how many tools detected a given circRNA, circRNA strand information was discarded.

#### Statistical analyses

To evaluate the effect of circRNA characteristics on circRNA validation, the Chi-squared test was used (*chisq.test()* function in R). For every test, the set of used circRNAs was slightly different depending on the availability of annotation information. All tests had all expected values in the contingency table above 5, therefore no correction for small sample size was necessary. Seven different characteristics were tested, and no multiple testing correction was performed. To evaluate the effect of circRNA detection tool methods on sensitivity and to evaluate the effect of the combination of tools with different approaches, the two-sided Mann-Whitney U test was used (*rstatix::wilcox_test()* function in R). For correlation analysis between the sensitivity and the extrapolated sensitivity, the Spearman rank correlation was used (*cor.test(method = ’spearman’)* function in R). For correlation analysis between circRNA BSJ counts from different tools, or between circRNA BSJ counts and Cq values, or between Cq values in different cell lines, linear models were used (*lm()* function in R). To evaluate the contribution of the cell lines (in contrast to the tools) to the precisions and sensitivity values, an ANOVA test was used (*aov()* function in R).

## Data availability

We anticipate this study will serve as a future resource for the circRNA community. The information on all predicted circRNAs (n = 315,312), including the large extensively validated circRNA set (n = 1,516), along with the validation results are available in the GitHub repository https://github.com/OncoRNALab/circRNA_benchmarking and as Supplementary Tables. Raw FASTQ files are stored in the Sequence Read Archive (PRJNA789110: SRX13414572 (untreated HLF), SRX13414573 (untreated NCI-H23), SRX13414574 (untreated SW480), SRX13414575 (RNase R treated HLF), SRX13414576 (RNase R treated NCI-H23), SRX13414577 (RNase R treated SW480)).

## Code availability

All the scripts used to compute the metrics described in the study and generate the figures are available at https://github.com/OncoRNALab/circRNA_benchmarking.

## Acknowledgments

We thank Steve Lefever for his contribution to primer design in the early stage of this project. The computational resources (Stevin Supercomputer Infrastructure) and services used in this work were provided by the VSC (Flemish Supercomputer Center), funded by Ghent University, FWO, and the Flemish Government – department EWI. Stefania Bortoluzzi would like to acknowledge support of Fondazione AIRC per la Ricerca sul Cancro, Milan, Italy (Investigator Grant 2017 20052), Italian Ministry of Education, Universities, and Research (PRIN 2017 2017PPS2X4_003), EU within the MUR PNRR “National Center for Gene Therapy and Drugs based on RNA Technology” (Project no. CN00000041 CN3 RNA) and "HPC, big data and quantum computing" (CN1 HPC) and the Department of Molecular Medicine of the University of Padova. Trees-Juen Chuang would like to acknowledge the National Health Research Institutes, Taiwan (NHRI-EX110-11011B1). Christoph Dieterich would like to acknowledge the German Science Foundation (DI 1501/13-1). Paul Flicek would like to acknowledge Wellcome Trust (WT108749/Z/15/Z). Enrico Gaffo would like to acknowledge the Fondazione Umberto Veronesi, Milan, Italy (Fellowship 2020). Steve Hoffmann would like to acknowledge the German Federal Ministry of Education and Research (BMBF 031L0106D). Eric C. Lai would like to acknowledge the NIH (R01-NS083833) and MSK (Core Grant P30-CA008748). Peter Stadler would like to acknowledge the German Federal Ministry of Education and Research (BMBF 031L0164C). Jakub Westholm would like to acknowledge the Knut and Alice Wallenberg Foundation as part of the National Bioinformatics Infrastructure Sweden at SciLifeLab. Li Yang would like to acknowledge the National Natural Science Foundation of China (NSFC, 31925011) and the Ministry of Science and Technology of China (MoST, 2021YFA1300503). Chu-Yu Ye would like to acknowledge the National Science Foundation of China (31871589).

## Author contributions

Validation co-author group: Marieke Vromman: methodology, software, validation, formal analysis, investigation, data curation, writing - original draft, visualization, funding acquisition; Jasper Anckaert: software; Justine Nuytens: investigation; Olivier Thas: methodology; Eveline Vanden Eynde: investigation; Kimberly Verniers: investigation; Nurten Yigit: investigation; Jo Vandesompele: conceptualization, methodology, writing - review & editing, supervision, project administration, funding acquisition; Pieter-Jan Volders: conceptualization, methodology, writing - review & editing, supervision, project administration, funding acquisition.

CircRNA prediction co-author group (contribution: formal analysis and writing - review & editing): Stefania Bortoluzzi, Alessia Buratin, Chia-Ying Chen, Qinjie Chu, Trees-Juen Chuang, Roozbeh Dehghannasiri, Christoph Dieterich, Xin Dong, Paul Flicek, Enrico Gaffo, Wanjun Gu, Chunjiang He, Steve Hoffmann, Osagie Izuogu, Michael S. Jackson, Tobias Jakobi, Eric C. Lai, Julia Salzman, Mauro Santibanez-Koref, Peter Stadler, Guoxia Wen, Jakub Westholm, Li Yang, Chu-Yu Ye, Guo-Hua Yuan, Jinyang Zhang, Fangqing Zhao

