## Supplementary Information for "Large-scale benchmarking of circRNA detection tools reveals large differences in sensitivity but not in precision"

### Supplementary Notes

#### CircRNA detection

The developers of the following circRNA detection tools were invited: ACFS2, CircCode, CIRCexplorer2, circLGB, CirComPara, CircParser, CIRCPlus2, circRNA_finder, CircRNAwrap, circseq_cup, CircSplice, circtools, CIRI2, ecircscreen, Fcirc, find_circ, KNIFE, NCLscan, PTESFinder, Sailfish-cir, seekCRIT, segemehl, SRCP, STARChip, UROBORUS. Some tool developers declined for various reasons; others never responded to repeated invitations via email. In total, 16 different circRNA detection tools were included: CIRCexplorer3 (1), CirComPara2 (2), circRNA_finder (3), circseq_cup (4), CircSplice (5), circtools (6), CIRI2 (7), CIRIquant (8), ecircscreen (unpublished tool), find_circ (9), KNIFE (10), NCLscan (11), NCLcomparator (12), PFv2 (13), Sailfish-cir (14), and segemehl (15).
Each circRNA detection tool was run by the developer(s) of the tool as described below.

1. CIRCexplorer3/CLEAR

FastQC (https://www.bioinformatics.babraham.ac.uk/projects/fastqc/) was first applied (parameters: fastqc -o output_dir mate_fastq.gz) to examine the quality of three raw RNA-seq datasets used in this benchmark study (RNAs individually isolated from HLF, NCI-H23 or SW480 cell line). In our hands, all these three RNA-seq datasets have passed the quality control by FastQC with satisfied per base sequence quality (the lower quartile of quality score for each base was greater than 10, and the median of quality score for each base was greater than 25) and adapter content (the percentage of adapter sequences in reads was less than 5%), and thus we did not trim or remove any reads. Next, we applied CIRCexplorer3/CLEAR (1) to identify and quantify circular RNAs individually from three datasets (parameters: clear_quant -1 mate_1.fastq.gz -2 mate_2.fastq.gz -g hg38.fa -i hg38.hisat2_index -j hg38.bowtie_index -G GENCODE.v31.annotation.gtf -o output_dir). Of note, hg38.hisat2_index and hg38.bowtie_index were built with default parameters by HISAT2 and BowTie, respectively. After analyzing with CIRCexplorer3/CLEAR, two types of circular RNAs, circRNAs from back-splicing of exons and ciRNAs from spliced intron lariats, were recorded in the final output file individually for each dataset. All used bioinformatic tools and packages are listed here: Python (v2.7.15), FastQC (v0.11.8), CIRCexplorer3/CLEAR (v1.0), CIRCexplorer2 (v2.3.8), HISAT2 (v2.1.0), BowTie (v0.12.9), SAMtools (v1.9), TopHat2 (v2.0.13), StringTie (v2.1.2), pybedtools (v0.7.5) and pysam (v0.16.0.1).

Parameter choice: The source code of CLEAR/CIRCexplorer3 has been optimized to include the necessary parameters for read alignment and circRNA annotation, which are well encapsulated and do not need to be modified. The default parameters are the most optimal to run the pipeline on a standard RNA sequencing dataset.

2. CirComPara2

CirComPara2 v0.1 was run with default parameters except for the read pre-processing parameters, which were PREPROCESSOR = "trimmomatic" and PREPROCESSOR_PARAMS = "MAXINFO:40:0.5 LEADING:20 TRAILING:20 SLIDINGWINDOW:4:30 MINLEN:50 AVGQUAL:26". From the detected circRNAs, circRNAs with back-spliced ends less than 300 bp apart (min length) and farther than 2,304,996 bp (max length) were discarded. The maximum length was determined from the genomic size of the largest gene expressed in the data.

Parameter choice: During development, the parameters of all tools CirComPara2 integrates were tuned to perform well as an ensemble. However, CirComPara2 employs a "consensus" strategy, and those tunings do not grant each tool to give its "best performance" when considered alone. Nonetheless, we ended up using default parameters for most circRNA tools. Moreover, one asset CirComPara2 showed was that it performed well on average among multiple, heterogeneous independent test (real) data sets and genome annotation "completeness" conditions, so there is not much more left to be tuned in a standard experiment on human cells. Since the sequencing dataset used for this benchmarking is "standard" and of high quality, the only parameter that was set differently from the default in CirComPara2 was related to the read preprocessing step: we allowed a less stringent quality filtering of the reads, which should allow a bit higher sensitivity (NB: CirComPara2's default is very stringent as it discards reads with an average quality score below 30 Phred; for the benchmarking study, we allowed 26 Phred score, which is still high).

3. circRNA_finder

First, a reference genome for STAR was created, including a database of splice junctions based on Gencode gene annotation and a splice junction overhang of 100. Next the circRNA_finder script runStar.pl was run in default mode on each library. Finally, all results were processed using the circRNA_finder script postProcessStarAlignment.pl in default mode. The following versions were used: circRNA_finder v1.2, STAR v 2.7.2d.

Parameter choice: The only parameter that was adjusted was to require at least 5 BSJ junction spanning reads to call a circular RNA.

4. circseq_cup

CircRNAs were identified by circseq_cup (v1.0) using default parameters as described previously (4). Briefly, TopHat-Fusion was used to find candidate fusion junction sites. Then, genomic sequences between two junction sites were extracted and duplicated once to form pseudo-references, of which the middle two nucleotides were the potential back-splicing sites of circRNAs. Tophat (v2.1.0) was used to align unmapped reads collected in the first step to pseudo-references to confirm the back-splicing sites. Only reads that mapped across the middle two nucleotides (i.e. back-splicing sites) at least 10nt were collected for full-length assembly by Cap3. After filtering criteria described by Ye *et al.* (16), assembled sequences that were consistent with the reference sequences were considered as full-length sequences of candidate circRNAs.

Parameter choice: The most important parameter is "-l" which indicates the maximal length between the candidate's start and end position. The length of most circRNA sequences was under 1000 bp. So, we used the default value (5000 bp) for this parameter. Otherwise, the larger of this parameter, it may predict more false positive circRNA candidates.

5. CircSplice

CircSplice was performed with default parameters. Briefly, FastQC(v0.11.9) was used to examine the reads quality, and adapters and low-quality bases were removed by trim_galore(v0.6.6) with parameters -q 20 --phred33 --stringency 3 --length 20 -e 0.1 --paired, then the STAR aligner(v2.7.6a) was used to align reads to reference genome. Finally, CircSplice.pl was used to identify circRNAs.

6. circtools

After initial quality assessment, low quality regions and adapter sequences were removed with Flexbar (version 3.5). Residual rRNA reads were removed using Bowtie2 with an rRNA sequence-based index. Principal read mapping against the ENSEMBL reference genome build 100 was performed with the STAR RNA-seq aligner (version 2.7.5a, with command line arguments as specific at https://docs.circ.tools/en/latest/Detect.html). CircRNAs were detected with circtools version 1.2.0 using default settings. CircRNA reconstruction was performed using the circtools reconstruct module using settings for ENSEMBL-based genomes (see https://docs.circ.tools/en/latest/Reconstruct.html and (17)). The quality of sequencing data and mapping results was assessed with MultiQC (version 1.9) as well as with the circtools quickcheck module using default parameters.

Parameter choice: The circtools detect step was mainly run with default parameters, which have been found to be most suitable to produce optimal results. However, a few parameters were modified to achieve the best performance for the given dataset. The -ss flag was enabled due to the stranded nature of the dataset, moreover, circtools was run with -Pi, -mt1, and -mt2 flags since the library was generated in paired-end mode. In order to achieve high sensitivity on the one hand and not introduce false positive calls, the -Nr flag was set to "2,2", which requires BSJs to be detected in at least two samples with two BSJ covering reads. Additionally, a RepeatMasker file was provided via -R to filter out circRNAs from repetitive regions.

7. CIRI2 and CIRIquant

To run CIRI2, the human reference genome and gene annotation files were downloaded from the GENCODE (Release 29, GRCh38.p12) project. Firstly, the BWA and HISAT2 indexes of the reference genome were built using to perform sequence alignment. Then, sequencing reads were aligned to the reference genome using BWA mem (18) (v0.7.17) with the “-T 19” option, and back-spliced junction sites of circRNAs were predicted using CIRI2 (7, 19) (v2.0.6) with the “-0” option. For CIRIquant analysis, BWA (18) (v0.7.17), HISAT2 (20) (v2.1.0), StringTie (21) (v1.3.3b), and Samtools (v1.11) were used, and CIRIquant (8) (v.1.1) was used for quantification of circRNAs with the default parameters. No extra filtering step was performed.

Parameter choice: CIRI2 and CIRIquant programs were run with the default parameters. The key parameters (anchor length, mapping quality threshold, alignment length, etc.) in CIRI2 and CIRIquant have been chosen based on both simulation and experimental validation datasets to ensure the performance of our programs.

8. Ecircscreen

Input reads were first assessed using MultiQC (https://multiqc.info), trimming off errant adapter sequences and regions with quality scores lower than 15 using trim_galore (https://github.com/FelixKrueger/TrimGalore.git). Genome index files for GRCh38 were built using STAR (22) (v. 2.7.10a), BWA (18) (v. 0.7.17-r1188) and Bowtie2 (23) (v. 2.3.4). Reads were aligned to the genome using respective aligners: STAR (--alignIntronMax:1000000; -–outFilterMultimapNmax:10; -–outFilterMismatchNmax:2), Bowtie (--very-sensitive --score-min=C,-15,0 --mm -q) and BWA mem (-T 19). To mitigate aligner-specific biases, we developed a computational pipeline that pools predictions from five publicly available circRNA prediction tools (CIRI2, circRNA_finder, PFv2, find_circ, and CIRCexplorer) using the eHive pipeline management system (https://ensembl-hive.readthedocs.io/en/version-2.6/). Additional filters were applied to exclude: 1) predictions supported by fewer than three methods, 2) predictions overlapping segmental duplicated regions (24) in the genome, and 3) predictions with flanking regions of the BSJ overlapping retrotransposons by over 25%. Genomic region overlaps were assessed using intersectBed and annotateBed from BEDTools (v. 2.26.0 [https://github.com/arq5x/bedtools2.git]). Sequence constructs of candidate BSJ predictions were generated by concatenating 100bp of sequence flanking BSJ coordinates in a head-to-tail manner, consistent with back-splicing. Reads were remapped to the BSJ Bowtie2 index files and mapped reads extracted to generate read counts for each prediction in the final output.

Parameter choice: Ecircscreen aims to reduce aligner-specific biases, excluding predictions reported by fewer than 3 identification tools and predictions overlapping low-complexity regions of the genome. As default parameter values within ecircscreen are conservative, adjustments likely increase sensitivity and reduce specificity.

9. find_circ

Find_circ is the only tool that was run by the validation co-author group and not by the tool developers, as we did not receive feedback from the tool developers. For this, total RNA sequencing samples were first trimmed using cutadapt (version 1.18) to remove adaptors. Next, Biopython (version 1.72) was used to remove reads with low QC and clumpify was used to remove duplicate reads. Bowtie2 (version 2.3.4.1) was then used to align all reads against the reference genome, and SAMtools (version 1.7) was used to extract all unmapped reads. Next, find_circ is run on the unmapped reads by using their unmapped2anchors.py and find_circ.py scripts. The output was not filtered additionally.

10. KNIFE

KNIFE (https://github.com/lindaszabo/KNIFE) was run with default parameters, except for 13 for the minimum junction overlap for considering a read is aligned to a junction. Reads were aligned to the reference genome and databases of linear and back-splice junctions created based on GENCODE v31 annotations using Bowtie2.2.1. The Bowtie2 parameters were set as suggested for KNIFE (10). Each linear or back-splice junction in the database has 300 bps from two annotated exons. Shorter exons are N-padded so that all junctions have the same length. circRNAs with at least 2 reads and posterior probability (the estimated likelihood by KNIFE that a candidate circRNA is a true positive) of 0.9 or higher were called separately for each library.

Parameter choice: KNIFE was run with default parameters. The default parameters for KNIFE were chosen based on its extensive benchmarking and evaluation of a wide range of datasets as reported in its original paper (10).

11. NCLscan and NCLcomparator

CircRNAs of "NCLscan_results" were identified by NCLscan (version 1.6.5; https://github.com/TreesLab/NCLscan) (11) with default parameters on the basis of the human reference genome (GRCh38) and the Ensembl annotation (version 97). Here a NCLscan-identified event must simultaneously satisfy the following two rules: (i) at least one junction read supported the BSJ and spanned the BSJ by ≥ 10 bp on both sides of the BSJ, and (ii) the collection of the supporting junction and encompassing reads spanned the BSJ by ≥ 50 bp on both sides of the BSJ. The spanning range (i.e., ≥ 50 bp) was set because the read length of currently-available RNA-seq data was generally more than 50 bp. For accuracy, "NCLscan_post_results" further eliminated circRNAs of "NCLscan_results" that were potential false positives derived from ambiguous alignments (i.e., the circRNAs have an alternative co-linear explanation or multiple matches against the reference genome) using NCLcomparator (version 1.0.0; https://github.com/TreesLab/NCLcomparator) (12) with default parameters. In brief, the exonic circle sequences flanking a BSJ (± 100 nucleotides of the BSJ) were concatenated using BEDTools (version 2.25.0). The concatenated sequence was then aligned against the reference genome and the Ensembl-annotated transcripts using BLAT (version 36) with the default parameters. The concatenated sequence of the NCLscan_post BSJ event should not map to an alternative co-linear sequence with more than 80% identical to the concatenated sequence or multiple hits with a similar BLAT mapping score < 3. Both NCLscan and NCLcomparator were implemented under Bio-Linux (Ubuntu 18.04).

12. PFv2

Input sample reads were preprocessed as described above for ecircscreen, using MultiQC (https://multiqc.info) and trim_galore (https://github.com/FelixKrueger/TrimGalore.git). To enhance sensitivity and potentially detect circRNAs within intergenic, antisense, and unannotated regions of the genome, the original PTESfinder (25) methodology that relied on existing transcriptome annotations was modified for annotation-free circRNA identification in PFv2 (https://github.com/osagiei/pfv2). Briefly, the underlying aligner for identifying reads with short sequence anchors that map head-to-tail, Bowtie1, was replaced with STAR (v. 2.7.10a) to detect putative BSJ predictions. Next, sequence constructs of ~200bp in size were generated using coordinates of predictions defined by STAR, by joining sequence regions flanking coordinates in a manner consistent with back-splicing. Bowtie2 index files were generated for the sequence constructs and the predictions evaluated by re-mapping reads using Bowtie2 (v. 2.3.4). To exclude BSJ supporting reads potentially arising from known sources of artifacts, we applied three filters that compared BSJ alignment quality scores to alignment scores obtained when same reads were mapped to the genome and transcriptome references. The transcriptomic filter is only effective for predictions originating from annotated loci and not intergenic. A final filter was applied to exclude reads mapping suboptimally to the BSJ by removing reads with mismatches or indels within 4bp proximal to the BSJ (JSpan=8) and with sequence percent identity to the BSJ exceeding 85% (PID=0.85). Read counts were generated by counting reads recovered after these filters for each BSJ.

Parameter choice: Within PFv2, default parameter values for examining read mapping across putative BSJs were found to maximize the identification of true positives, whilst reducing false positive predictions using both simulated and real RNAseq data (13). However, to identify novel BSJs from intergenic, antisense, and intronic regions of the genome, PFv2 is unconstrained by existing transcript annotations.

13. Sailfish-cir

First, we identified all putative circRNAs using CIRI2 with default parameter settings. After circRNA identification, we run the main module of wf_profile_circRNA.py of Sailfish-cir with a configuration file, which transformed circRNAs to pseudo-linear transcripts and then specified Sailfish (version 0.10.0) to quantify the transcripts per million (TPM) values of all identified circular transcripts and known linear transcripts in the Ensembl human gene annotation (release 94) and human genome GRCh38 with default settings.

Parameter choice: In Sailfish-cir implementation, we used the CIRI2 package in identifying circular RNAs with the default parameters suggested in CIRI2. Therefore, Sailfish-cir implementation should have the same justification as CIRI2 in choosing the parameters.

14. segemehl

Fastq files were first trimmed (trimmomatic, v0.39; mean quality cutoff of 22 and a minimum read length of 18nt) and clipped (cutadapt v 2.10; default parameters). Subsequently, reads were corrected (rcorrector, v1.0.4; default parameters) and filtered for rRNAs (SortMeRNA, v2.1; default parameters). Afterward, reads were mapped with segemehl (v0.3.4) to the reference genome hg38 in split read mode (-S) using an overall accuracy of 95% (-A95) and a maximum insert size of 200kB (-I 200000) to search for paired ends using the regular alignment algorithm. Subsequently, the split read alignments were extracted and summarised using `haarz split.` To increase the pipeline's specificity, circular junctions annotated as type 'C' were extracted only if supported by at least five split read alignments using a custom script. Finally, all circular junctions supported by more than five circular split reads in at least two replicates were reported.

Parameter choice: As the parameters for the segemehl circ-mode are concerned, they were selected to ensure a low false positive rate. More specifically, all circ junctions with less than 5 supporting reads were discarded. Also, all junctions spanning more than 200kB were also discarded.

### Supplementary Data

#### Supplementary Data 1: Annotating circRNAs is not always straightforward

To generate data regarding circRNA length and exon composition, annotation of the linear host transcript is required. For this, the circRNA positions were compared with a list of canonical transcripts. Of note, not all circRNAs could be uniquely assigned to a specific host gene (Supplementary Figure 7).

#### Supplementary Data 2: Some detection tools filter on predicted circRNA length

As this study focuses on the evaluation of circRNA detection tools based on short-read RNA sequencing, the full circular transcript sequence and length remain unknown. Nevertheless, the theoretical length of a circRNA can be computed by considering either inclusion or exclusion of introns of canonical transcripts (Supplementary Figure 8). The median length (including all exons and introns) of a circRNA is 1,660 nucleotides (interquartile range (IQR) 454 - 9,925). Most tools show a similar circRNA length distribution, with circseq_cup and segemehl reporting generally shorter circRNAs than the other tools. Although the circRNA length calculated with introns is no measurement of the true circRNA size, the majority of circRNA detection tools filter circRNAs based on this estimation (Table 1). Overall, all reported circRNAs have a length of at least 31 nucleotides. When taking exon annotation into account, the 'exonic' circRNA length can also be computed, assuming all exons are retained, and all introns are removed (Figure 2C). The median length (excluding all introns) of a circRNA is 498 nucleotides (interquartile range (IQR) 286 – 917). This is in line with previous reports on circRNA length (26–28). Like the calculation with intron inclusion, circseq_cup and segemehl generally report shorter circRNAs compared to the other tools.

#### Supplementary Data 3: Most circRNAs are predicted to consist of 2-5 exons

Based on the circRNA exon annotation, the number of exons per circRNA was determined (assuming all exons are retained) (Supplementary Figure 9). Most tools have a similar distribution of the number of exons per circRNA, with the majority of circRNAs containing between 2 and 5 exons. PFv2 has a higher number of circRNAs for which no annotation match was found compared to the other circRNA detection tools.
Supplementary Figure 9 also shows if the circRNA start and end positions of a circRNA correspond to existing linear host gene annotations. Of note, just like circRNA length, the full circRNA sequence remains unknown, and the number of exons per circRNA is an upper bound estimation. Every tool also reports a small number of circRNAs (median 51 circRNAs, IQR 20-87 circRNAs) including the first exon of a canonical transcript. Only one circRNA containing the first exon was selected (by chance) for empirical validation. This circRNA could not be validated with amplicon sequencing.

#### Supplementary Data 4: CircRNAs are transcribed from both DNA strands

All tools except CirComPara2, circseq_cup, Sailfish-cir, and segemehl report the DNA strand from which the circRNA is transcribed. Of note, a more recent version of CirComPara does report strand information, and now circseq_cup as well). CircRNA strand information is not always acquired in the same way by the different tools (Table 1). All tools that provide circRNA strand information, report approximately 50% of the circRNAs on the positive strand, and 50% of the circRNAs on the negative strand (Supplementary Figure 10). Ecircscreen is the only tool that reports more circRNAs on the positive strand (~60%). In comparison, linear transcripts originate almost equally from the positive or the negative strand (based on Ensembl annotation, version GRCh38.103).

#### Supplementary Data 5: Pilot experiment to assess low abundant circRNA detectability with qPCR and establish count cut-off

To estimate the circRNA detection limit with our RT-qPCR method, 100 circRNAs detected by CIRCexplorer2 with BSJ counts in different bins were selected. More specifically, 50 circRNAs with BSJ count of 1 were selected, and 50 with BSJ count above 1. A circRNA was deemed detectable when the mean of the two qPCR replicates was lower than 32. This threshold was chosen to ensure it was still possible to measure circRNA degradation by Rnase R if there would be any. Based on the results of this experiment (Supplementary Figure 11), all analysis were split up between *low-abundance* circRNAs (BSJ count < 5) and *high-abundance* circRNAs (BSJ count ≥ 5).

**Supplementary Data 6: PCR primer design bias**

qPCR primers were designed for the selected circRNAs using CIRCprimerXL (29). As primer design can fail for multiple reasons (predicted off-target amplification, secondary structures, common SNPs, or limited design space), a larger set of circRNAs was selected for primer design. It is important to note that primer design introduces an inherent bias in our study, as the circRNAs for which no primers could be designed cannot undergo empirical validation. These can include circRNAs with problematic repeat sequences or too short circRNAs (CIRCprimerXL requires a minimum of 60 nucleotides for primer design). Overall, most circRNA detection tools have similar primer design success rates (median 86.8%, IQR 85.1%-87.7%), apart from PFv2 that has a primer design success rate of 51.4%. Remarkably, this lower design rate is only observed for the *high-abundance* circRNAs. When evaluating why primer design was not successful, PFv2 showed more primers (57.8%) discarded due to predicted off-target amplification compared to the other tools (median 38.2%, IQR 35.1%-42.2%) (Supplementary Figure 12). Of note, PFv2 is a discovery tool deliberately unconstrained by transcript annotation to identify novel circRNAs within poorly annotated/highly repetitive human lncRNAs, intergenic, antisense, and intronic regions. The higher rate of predicted off-target amplification for *high-abundance* circRNAs compared to *low-abundance* ones may result from overlap with highly repetitive, low complexity regions of the genome, explaining their high counts and the high number of predicted off-target effects.
Interestingly, for 47 circRNAs selected for validation (3.2%), the designed primer pairs were not unique for that circRNA (column *primer_unique* in Supplementary Table 3). This is primarily due to selected circRNAs with similar positions (often in regions with repeats). Without full-length circRNA sequencing, we cannot distinguish between these circRNAs. Furthermore, from the selected circRNAs, 35 circRNAs (2.4%) were selected for both strands (detected by different tools).

**Supplementary Data 7: Comparison to RNase R treated sequencing data**

RNase R is also often used in combination with RNA sequencing to validate circRNAs (30, 31). Upon RNase R treatment, a *bona fide* circRNA is expected to be enriched as fewer sequencing reads are taken up by linear RNA molecules. Often (but not always), the BSJ counts are first normalized by the total number of reads per sample, generating a BSJ CPM (counts per million reads). Then the CPM of the treated sample is divided by the CPM of the untreated sample to create an enrichment factor. Some researchers label a circRNA as enriched if the enrichment factor is ≥ 1, and some use enrichment ≥ 5. Often, a BSJ count threshold is used to calculate the fraction of enriched circRNAs in a specific sample.
We also included this analysis for the *high-abundance* circRNAs by computing the enriched factor as CPM treated / CPM untreated and labeling a circRNA as enriched when the enriched factor > 1 (Supplementary Table 5). Supplementary Figure 16 shows the effect of RNase R treatment on linear and circular reads for the TEAD1 transcript. Of note, many low abundant circRNAs are only present in one of the two samples (treated or untreated), which means a fraction of the data cannot be used (Supplementary Figure 17). However, this figure shows that the precision is mostly high and similar among tools (*i.e.*, most predicted circRNAs are enriched in RNA-seq data upon RNase R treatment), with PFv2 having the lowest precision value (similar to RNase R validation with RT-qPCR). Additionally, like RNase R validation in combination with RT-qPCR, there are significantly fewer enriched circRNAs in the *low-abundance* subgroup compared to the *high-abundance* group (Chi-squared = 1067, df = 1, p-value < 0.001, OR = 1.4). The distribution of the BSJ count enrichment factors can be seen in Supplementary Figure 18. Lastly, the cumulative enrichment factor in function of the BSJ count is represented in Supplementary Figure 19.

**Supplementary Data 8: The cell line effect on the results is limited**

To assess the effect of the different cell types, all metrics were recomputed per cell line (Supplementary Figures 27 and 28). Cell line NCI-H23 is associated with significantly lower precision (Chi-squared = 25.0, degrees of freedom = 2, p-value < 0.001), and several tools struggle particularly with this cell line. However, the overall ranking and the absolute precision values are highly similar among the tools over the cell lines, especially for the high abundance circRNAs (BSJ count ≥ 5). Of note, the metrics for the low abundance circRNAs (BSJ count < 5), are based on a small set of circRNAs (n = 20 per tool) and are therefore more variable.

Using two-way ANOVA tests to assess the effect of the cell line and the tool on the precision and sensitivity values, showed that the tool has a larger effect on each metric than the cell line, except for RNase R precision (sum of squares: 16.1% (cell line, p-value = 0.005) and 49.5% (tool, p-value = 0.008) for RT-qPCR precision, 64.5% (cell line, p-value = 0.001) and 28.0% (tool, p-value = 0.001) for RNase R precision, no significant effect (cell line) and 88.3% (tool, p-value < 0.001) for amplicon sequencing precision, 38.9% (cell line, p-value < 0.001) and 58.9% (tool, p-value < 0.001) for compound precision, and no significant effect (cell line) and 97.5% (tool, p-value < 0.001) for sensitivity).

**Supplementary Data 9: CircRNAs detected across different cancer cell lines**

Out of the 1,457 unique randomly selected circRNAs, 58 circRNAs were selected in more than one cell line. Of these, 57 circRNAs were tested in two cell lines, and one circRNA was selected to be validated in all three cell lines. This gives the opportunity to compare circRNA expression levels among cell lines and assess the agreement of the validation methods in different cell lines (Supplementary Figure 29). For 54/58 circRNAs (93.1%) all validation results were the same for all cell lines (these 53 circRNAs all happen to pass all validation methods or pass two validation methods and were not evaluated for amplicon sequencing). The 4 circRNAs (7.0%) that do not have the same validation results in different cell lines are all in the *high-abundance* count group (BSJ count ≥ 5). For these 4 circRNAs, the observed disagreement of validation results is consistently RNase R validation, whereas the other validation methods agree. Interestingly, the failed RNase R validation was always observed in the NCI-H23 cell line, while the same circRNAs could be validated in the HLF cell line (Supplementary Figure 30).

**Supplementary Data 10: CircRNA quantification correspondence analysis between RT-qPCR and total RNA sequencing**

As our study is the first to report matched Cq values and BSJ counts for an unprecedently large set of circRNAs, a correlation between Cq values and BSJ counts was computed for each tool (based on linear model, Supplementary Figure 31). Generally, a reasonable linearity is observed between these 2 orthogonal methods (median $r^2$ of 0.25, range 0.19-0.29, p<0.001).

**Supplementary Data 11: Comparison to simulated data reported by other studies**

An alternative method to benchmark circRNA detection tools is by using simulated data (2, 30, 32). One of the most comprehensive studies (30) compares 11 tools of which 9 tools were also included in our benchmarking study. This study uses a positive dataset of simulated reads, encompassing a total of 14,689 circRNAs detected in HeLa cells from CircBase (33). Their mixed dataset is the positive dataset in addition to a background dataset comprised of reads generated from mRNA sequences deposited in the NCBI Reference Sequence (RefSeq) database. A second study (32) uses three different simulated datasets: mix1 includes only circRNA and linear transcripts, mix2 includes mix1 and tandem RNAs, and mix3 includes circRNA and linear RNAs generated by CIRI-simulator and ART simulator.

Precision values are high and similar among the different studies, except for PFv2 (Supplementary Figure 35). This can easily be explained based on the chosen datasets. As the background dataset uses mRNA sequences, it does not contain a lot of repeat regions (these are mostly non-coding or intronic). However, as discussed in Supplementary Data 6, PFv2 is a discovery tool deliberately unconstrained by transcript annotation to identify novel

circRNAs within poorly annotated/highly repetitive human lncRNAs, intergenic, antisense, and intronic regions. This explains why PFv2 has such a low precision based on a real dataset compared to simulated data where repeated regions are mostly excluded.
For sensitivity, there are large differences among the tools, both for simulated data and our orthogonal validation (Supplementary Figure 36). The circRNA reads in the simulated data are mostly acquired from circRNAs in circBase, to which some of the circRNA detection tools have also contributed, which induces a bias. Therefore, it is not surprising that the absolute sensitivity values between both methods do not correlate for some of the tools. The simulated data and the real-world data are vastly different.

**Supplementary Data 12: Choice of performance metrics**
Based on the data generated in our study, for each circRNA detection tool T1, a complete contingency table can be generated, and all the corresponding performance metrics (such as sensitivity and specificity) can be calculated.

- TP (true positive): the circRNA was predicted by T1 and validated.
- FP (false positive): the circRNA was predicted by T1 but failed validation.
- FN (false negative): the circRNA was not predicted by T1 but was predicted by at least one other tool and was validated.
- TN (true negative): the circRNA was not predicted by T1 (but was predicted by at least one other tool T2, which implies T2 reported a FP) and failed validation.

It is essential to make a distinction between how the TP and FP values are obtained, in contrast to the FN and TN values. For determination of the TP and FP values, a random set of 100 circRNAs was selected for each tool (20 *low-abundance circRNAs* and 80 *high-abundance* circRNAs), allowing straightforward calculation of the precision value (TP/(TP+FP)) for each circRNA detection tool separately. For this, only the 100 randomly selected circRNAs per tool are used.
However, to determine the FN and TN values, we must rely on the predictions of other tools, which introduces an inherent bias, as a fixed set of circRNAs was randomly selected for each tool and not randomly based on the entire set of predicted circRNAs. Therefore, when all the circRNAs are grouped together to compute a FN and TN value for each tool, a bias is introduced.
This can be further demonstrated using an example: in total 1560 circRNAs were selected to be validated. From this set, 957 circRNAs were validated using three methods and are labeled 'true positive'. To calculate the FN value for tool T1, the set of 957 validated circRNAs is used, and the FN value is the number of circRNAs that were validated but not detected by T1. This is then used to calculate sensitivity. However, this set of 957 validated circRNAs is a biased pool of all the different tools. Therefore, tools with similar circRNA detection results, have a higher chance of reporting a larger subset of the 957 circRNAs and in general, tools that have more overlapping circRNAs with other tools have an (unwanted) positive bias for this analysis, depending on the number of similar tools (data not shown). A similar bias exists when determining the TN value for each tool in the same way. Additionally, the TN values are all very small numbers, as most circRNAs get validated. In summary, in our study, we mostly focus on validating the predicted circRNAs, and therefore use the TP and FP values (to determine precision), as these are straightforward and unbiased. The TN and FN are used as an alternative way to calculate sensitivity (compared to the theoretical number of true positive circRNAs or the extrapolated sensitivity).

### Supplementary Data 13: The standard error on the precision (FDR)

#### 1 General discussion on sample size for FDR estimation

To make a fair comparison between circRNA prediction tools, we strive for equal precision of the FDR estimates. The imprecision of an estimator is given by its variance (or standard error). In Section 2 we develop an expression for the variance of the FDR estimator. In particular, the variance is given by the expression

$$\text{FDR}(1-\text{FDR})\text{E}\left\{\frac{1}{P}\right\}$$

in which FDR is the true FDR (which we aim to estimate) and $P$ is the number of positives. This number of positives is distributed as a binomial distribution with parameters $N$ (number of sampled predicted circRNAs) and $\pi$ (the probability of a positive confirmation of the circRNA). The expression for $\text{E}\left\{\frac{1}{P}\right\}$ is quite complicated, but from the bimomial distribution we can see that it will decrease with increasing sample size $N$. In conclusion, the precision of the FDR estimator increases with increasing sample size $N$ (classical result in statistics), but it does not depend on the total number of predicted circRNAs. In other words, to make a fair comparison between the FDRs of different circRNA prediction tools, it is recommended to sample an equal number of predicted circRNAs for each tool, irrespective of the number of predicted circRNAs.

Two comments:

1. as the expression for the variance shows, the precision also depends on the true FDR, but this is of course unknown. For this reason, we can only compare precisions as if the FDRs are equal. Similar comment for the probability $\pi$ of a positive confirmation. This is also unknown and therefore we only compare precisions as if these probabilities are equal.
2. in the development of the development of the variance of the FDR estimator, we assumed that we sampled from an infinite population (i.e. as if the number of detected circRNAs is very very large for all tools). This is a wrong working assumption. However, when sampling from a finite population, the population size als matters. So here the number of predicted circRNAs will also affect the variance. The conventional correction factor (applied to the variance) is
$$\frac{M-N}{M-1}$$
with $M$ the population size (i.e. the number of predicted circRNAs), and, as before, $N$ the number of sampled circRNAs for validation. For our study, this correction factor ranges between $0.965$ and $0.999$. We believe it is fair enough to ignore this factor for the sample size calculation, and therefore we concluded that an equal sample size for each tool is the best option.

#### 2 Precision of FDR estimates

Let $N$ denote the total number of samples, of which $P$ are called positive and $FP$ are false positive. The FDR is then estimated as

$$\widehat{\text{FDR}} = \frac{FP}{P}.$$

We are looking for the variance of this FDR estimate.

$$\begin{aligned}\text{Var}\left\{\widehat{\text{FDR}}\right\} &= \text{Var}\left\{\frac{FP}{P}\right\} \\ &= \text{E}\left\{\text{Var}\left\{\frac{FP}{P} \mid P\right\}\right\} + \text{Var}\left\{\text{E}\left\{\frac{FP}{P} \mid P\right\}\right\}.\end{aligned}$$

Hence, we need

$$\text{E}\left\{\frac{FP}{P} \mid P\right\} = \frac{\text{E}\left\{FP \mid P\right\}}{P} = \frac{\text{FDR}P}{P} = \text{FDR}$$

and

$$\text{Var}\left\{\frac{FP}{P} \mid P\right\} = \frac{1}{P^2}\text{Var}\left\{FP \mid P\right\} = \frac{1}{P^2}P\text{FDR}(1-\text{FDR}) = \frac{\text{FDR}(1-\text{FDR})}{P}.$$

Since $\text{Var}\left\{FDR\right\} = 0$, we find

$$\text{Var}\left\{\widehat{\text{FDR}}\right\} = \text{E}\left\{\frac{\text{FDR}(1-\text{FDR})}{P}\right\} = \text{FDR}(1-\text{FDR})\text{E}\left\{\frac{1}{P}\right\}.$$

Since $P \sim \text{Bin}(N, \pi)$, with $\pi$ the probability of a positive, the expectation $\text{E}\left\{\frac{1}{P}\right\}$ becomes

$$\text{E}\left\{\frac{1}{P}\right\} = \sum_{x=0}^{\infty} \frac{1}{x}\binom{N}{x}\pi^x(1-\pi)^{N-x}.$$

### Supplementary Discussion

**Supplementary Discussion 1: Discrepancies between circRNA validation techniques**

As shown in Figure 3B, the validation results are sometimes conflicting for specific circRNAs. For example, 95.6% of putative circRNA chr1:90937484-90982370/- (hg38, 0-based) is degraded by RNase R, yet it is validated with amplicon sequencing (98.4% on-target amplification) and is detected by 14 different tools. Furthermore, the exact sequence of the BSJ could be confirmed in the amplicon sequencing data. It has been reported that long circRNAs are more susceptible to RNase R degradation (34). This specific circRNA is predicted to be 3705 nucleotides long (excluding introns), thus much longer than the median of 552 nucleotides for all tested circRNAs. In some cases, it could be that the primer pair is not specific enough and amplifies an off-target linear RNA. This would be visible as an increase in Cq upon RNase R treatment. However, this would then also have been visible in the amplicon sequencing as a lower percentage of on-target amplification. In total, 22 circRNAs are validated with qPCR and amplicon sequencing but are degraded by RNase R for at least 87.5% (*i.e.*, a difference of 3 cycles). The subset of these circRNAs for which annotation is available (n = 12), have a median length of 2382 nucleotides (IQR 2,110 - 3,483), respectively, compared to 498 nucleotides (IQR 286 - 917) for all circRNAs. An alternative hypothesis is that for these 22 cases, the primers amplify a mixture of circular and linear RNA, therefore showing a difference in Cq value upon RNase R treatment, but still including the expected BSJ sequence upon amplicon sequencing. Another subset of circRNAs with discrepant results is a set of 13 that can be detected by qPCR, but then fails RNase R and amplicon sequencing validation. In this case, RNase R can distinguish circRNAs from linear RNAs, and the results are confirmed with amplicon sequencing. We consider these as likely false positives, underscoring the fact that RT-qPCR validation on itself is not sufficient. A final subset of 92 circRNAs pass RT-qPCR validation and RNase R validation but fail amplicon sequencing. It is likely that (repetitive) RNAs have some resistance to RNAse R due to secondary structure, either internal or through base pairing with orthologs (34).

### Supplementary Tables

**Supplementary Table 1** Details of circRNA detection tools included in this study.

**Supplementary Table 2** A list of the circRNAs detected by the 16 different circRNA detection tools in the untreated samples. Because of large file size, only available on https://github.com/OncoRNALab/circRNA_benchmarking.

**Supplementary Table 3** A list of 1,560 selected circRNAs, with their initial detection information (tool, BSJ count), their primer information (including FWD and REV primer sequence), results from three validation methods (Cq value with and without RNase R, Cq difference, and amplicon sequencing percent on-target amplification), validation metrics, and annotation information.

**Supplementary Table 4** A list of the circRNAs detected by the 16 different circRNA detection tools in the RNase R treated samples. Because of large file size, only available on https://github.com/OncoRNALab/circRNA_benchmarking.

**Supplementary Table 5** RNase R enrichment factor calculated based on RNA sequencing data for each circRNA. Because of large file size, only available on https://github.com/OncoRNALab/circRNA_benchmarking.

**Supplementary Table 6** Precision values (RT-qPCR, RNase R, amplicon sequencing, and compound) and sensitivity per tool and BJS count group. The user can easily filter and order the circRNA detection tools based on their preferences.

**Supplementary Table 7** The number of circRNAs in the intersection and union of each combination of two tools, per cell line sample. Only for circRNAs with BSJ count ≥ 5.

**Supplementary Table 8** The number of circRNAs in the intersection and union of each combination of three tools, per cell line sample. Only for circRNAs with BSJ count ≥ 5.

**Supplementary Table 9** A list of the top-performing combinations of two tools. The list was composed by selecting the top 5 performing combinations in terms of the total number of detected circRNAs (union between both tools) and the weighted compound precision, for each cell line.

**Supplementary Table 10** Primer sequences for evaluation of RNAse R efficiency

| | forward sequence | reverse sequence |
|---|---|---|
| linear ACTB | CTGGAACGGTGAAGGTGACA | AAGGGACTTCCTGTAACAATGCA |
| circRNA chr1:117402185-117420649 | AGGTGTCTGTGTTTGAAGTC | TGCTCGAATTTCCTCTCTTG |

### Supplementary Figures

**Supplementary Figure 1** The number of predicted circRNA varies greatly among tools and most tools report a majority of circRNAs with a BSJ count below 5. Similar numbers of circRNAs are observed in the three different cell lines.

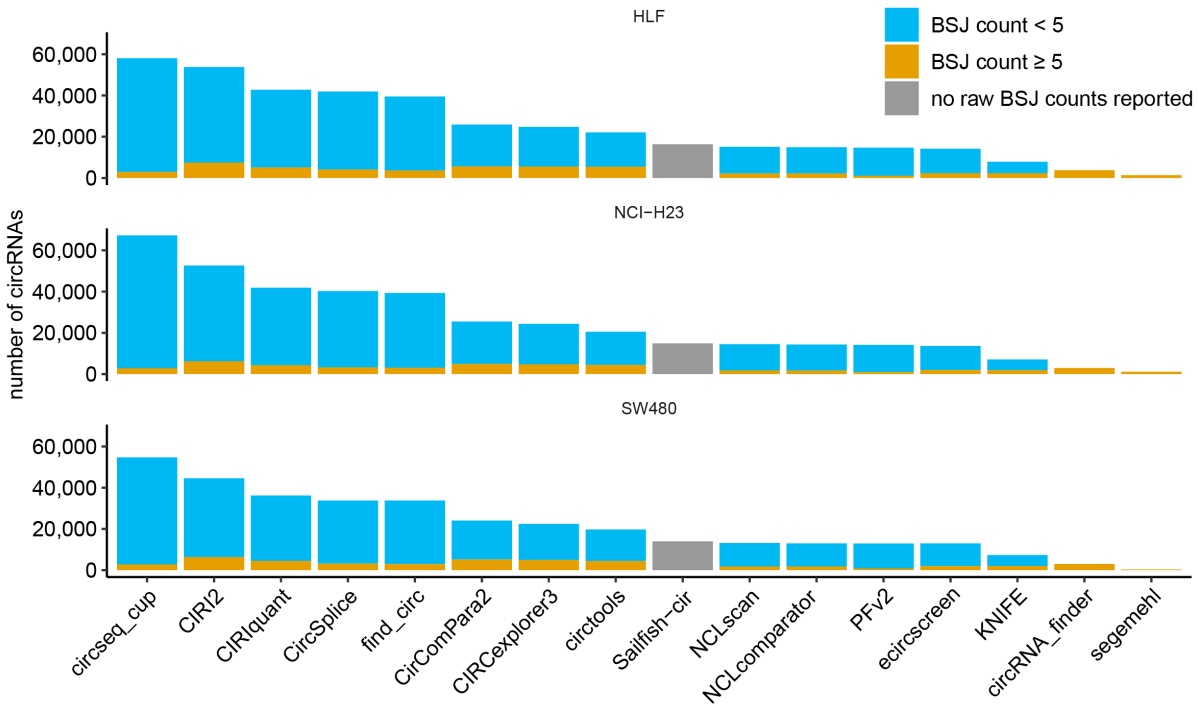


**Supplementary Figure 2** Most circRNAs are low abundant, as shown in these BSJ count boxplots per tool. CirComPara2 and KNIFE only report circRNAs with a BSJ count of at least 2, and circRNA_finder and segemehl only report circRNAs with a BSJ count of at least 5. Sailfish-cir does not report raw BSJ counts but reports transcripts-per-million (TPM) instead. The upper and lower hinges of the box correspond to the first and third quartile (interquartile range, IQR), respectively. The upper and lower whiskers extend from the hinge to the highest and lowest value that is within 1.5 * IQR of the hinge, respectively. Data beyond the end of the whiskers are outliers and plotted as points.

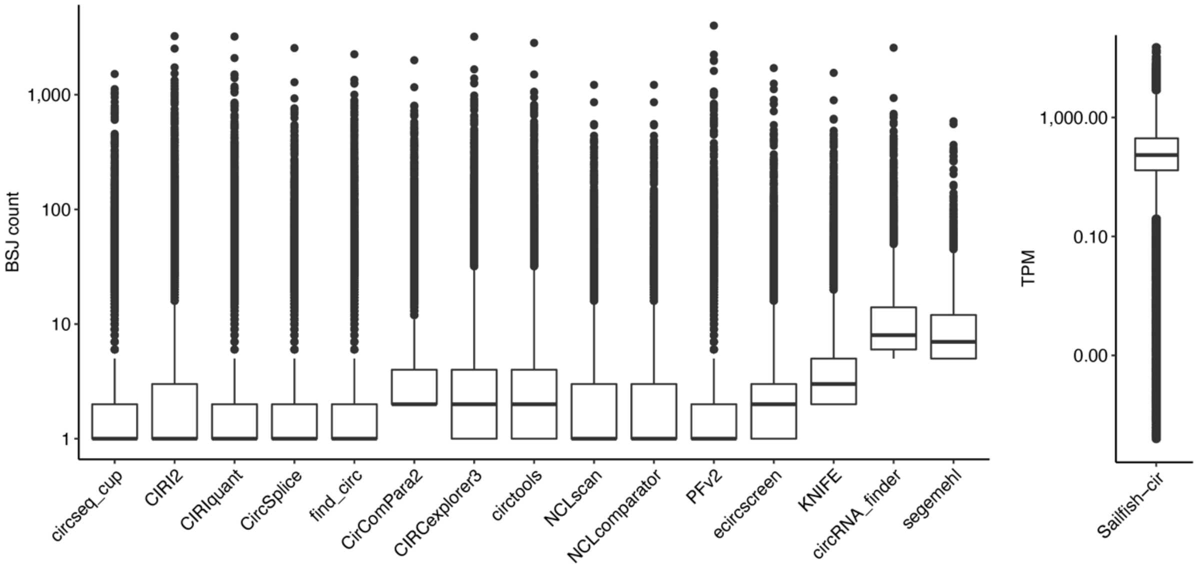

**Supplementary Figure 3** Correlation between log-scaled BSJ counts of the same circRNA detected by different tools is visualized by plotting the slope and R-squared value of the linear regression line (one value for each pair) (based on linear model). Perfect quantitative correspondence would result in slope and an R-squared of 1 (grey dashed lines). As NCLcomparator reports a subset of circRNAs detected by NCLscan with the same BSJ count, these two tools show a slope and an R-squared of 1.

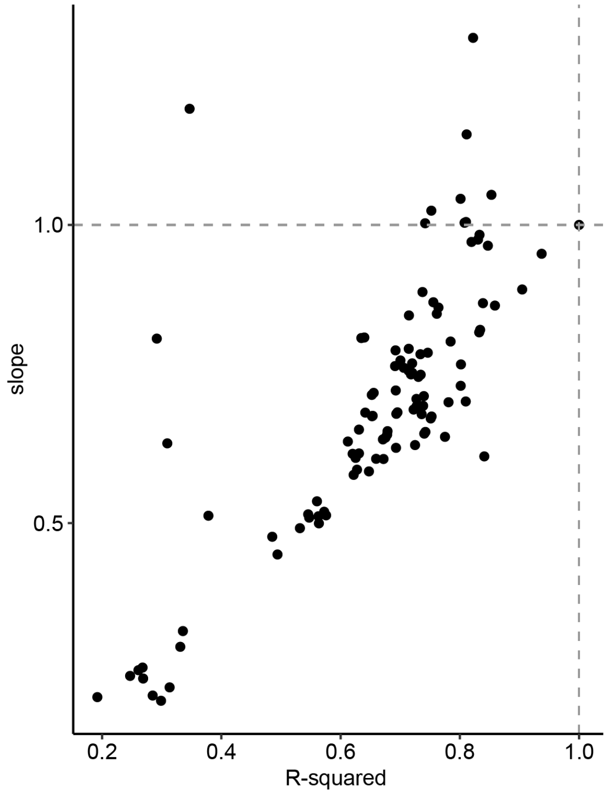


**Supplementary Figure 4** Most circRNAs are detected by more than one tool (when circRNA strand information is not taken into account). circseq_cup reports a high number of unique circRNAs. The same distribution is observed for all three cancer cell lines.

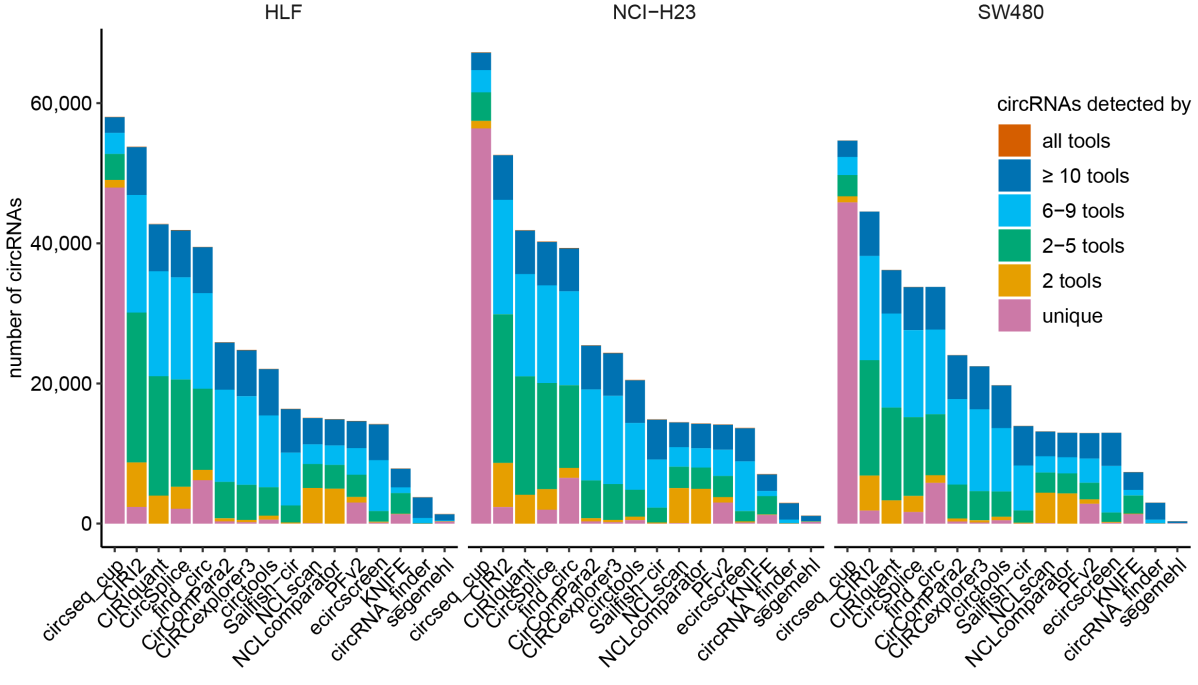

**Supplementary Figure 5** CircRNA detection tools cluster into multiple subgroups based on the Jaccard distance. As NCLcomparator is a subset of NCL_scan, it is no surprise that these two tools cluster together and have a very low Jaccard distance. Similarly, CIRI2 and CIRIquant (which is based on CIRI2) have a low Jaccard distance. Of note, the overlap among tools is highly dependent on the total number of circRNAs detected by a given tool. All three cell line samples show similar results.

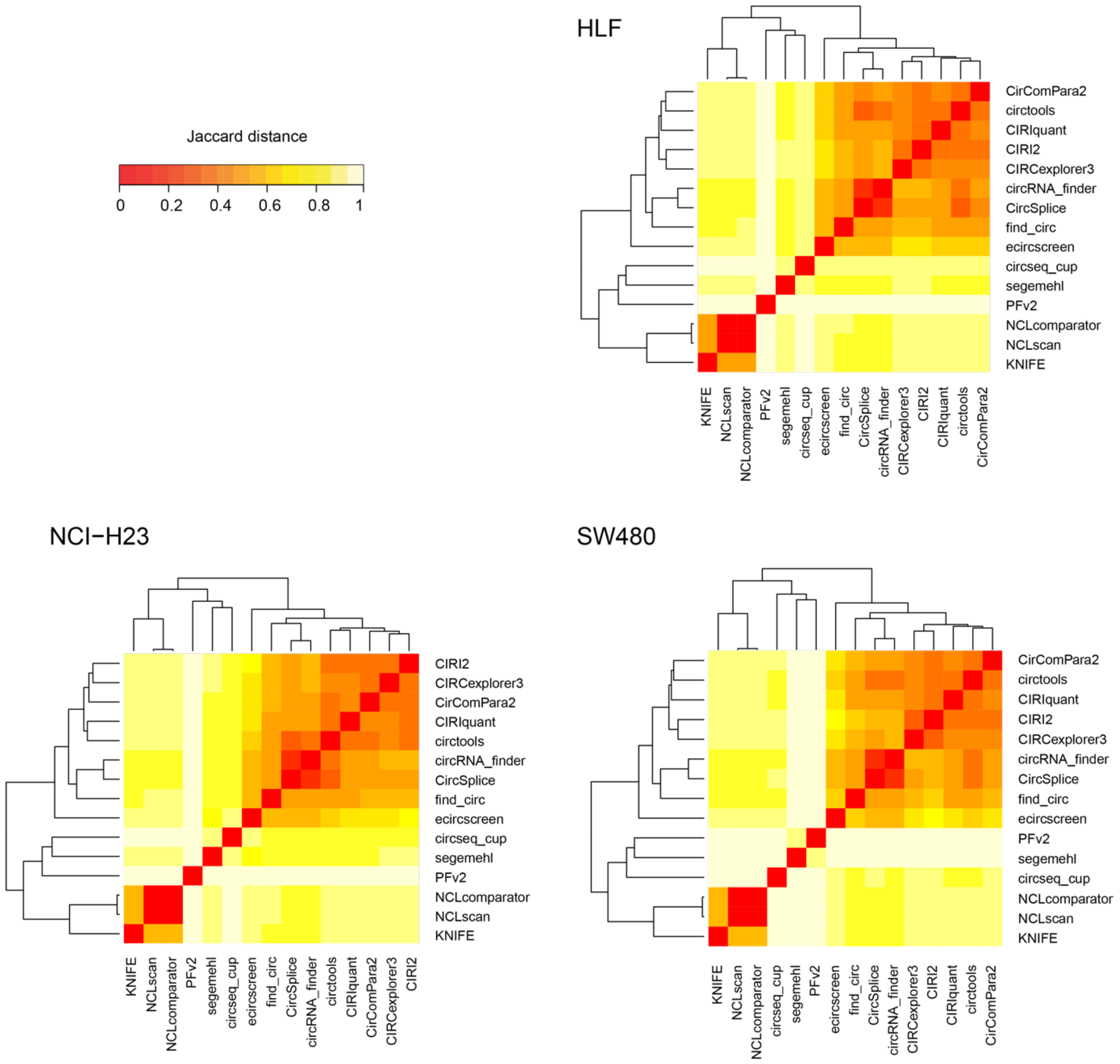

**Supplementary Figure 6** Most detected circRNAs have previously been described in circRNA databases (ignoring circRNA strand information). circseq_cup, KNIFE, NCLscan, and NCLcomparator predict many novel circRNAs.

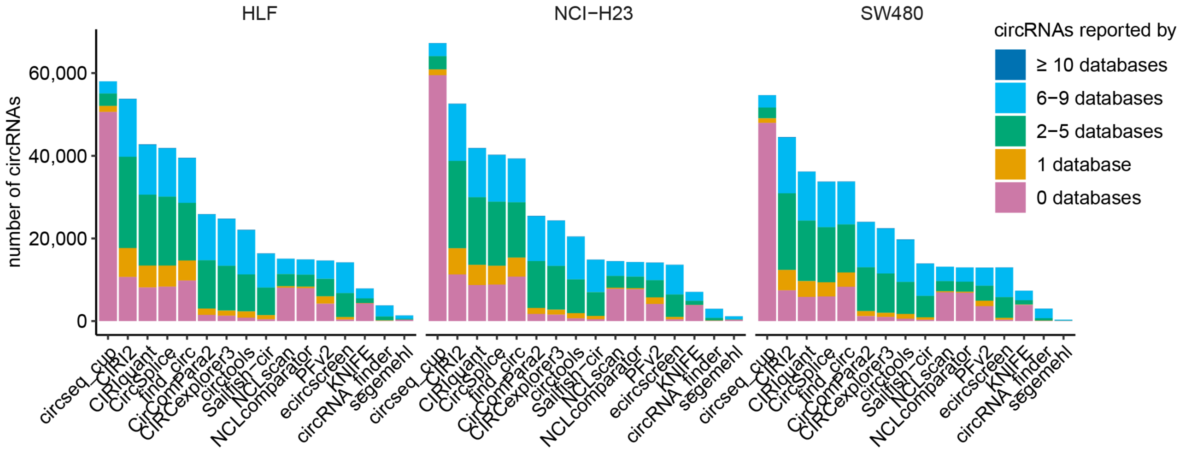


**Supplementary Figure 7** Most tools report a similar fraction of circRNAs that match linear annotation, except circseq_cup and PFv2.

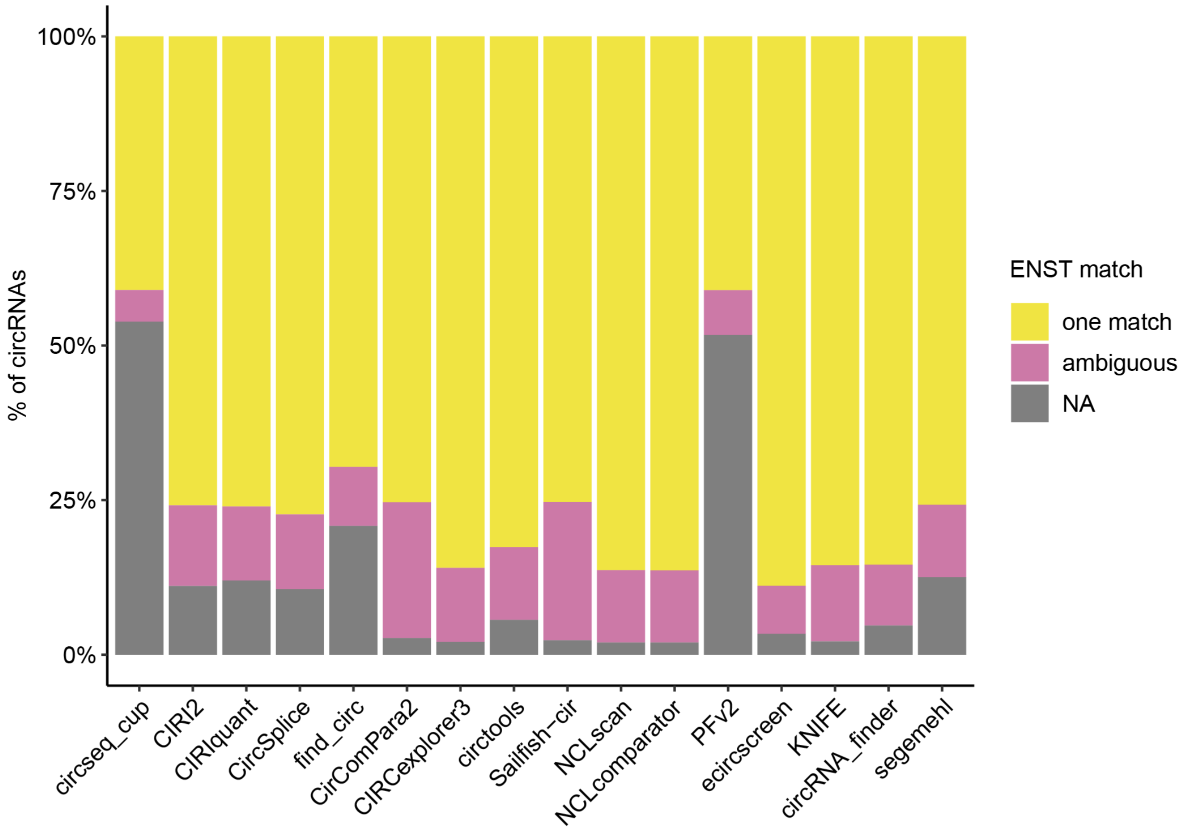

**Supplementary Figure 8** The circRNA predicted length distribution is similar among different tools. The upper and lower hinges of the box correspond to the first and third quartile (interquartile range, IQR), respectively. The upper and lower whiskers extend from the hinge to the highest and lowest value that is within 1.5 * IQR of the hinge, respectively. The outliers are not shown.

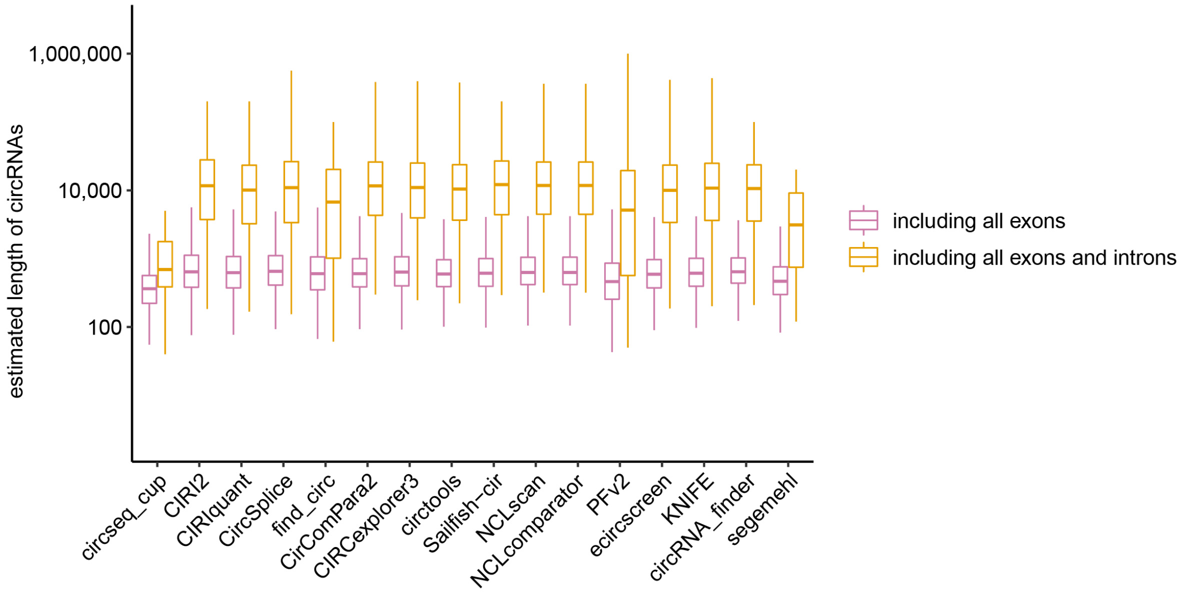


**Supplementary Figure 9** The distribution of the number of exons per circRNA is generally similar among tools. Circseq_cup and PFv2 result in a larger number of circRNAs with no canonical annotation match. Of note, it is more challenging to unambiguously annotate circRNAs without provided strand information, such as the circRNAs reported by circseq_cup, CirComPara2, Sailfish-cir, and segemehl.

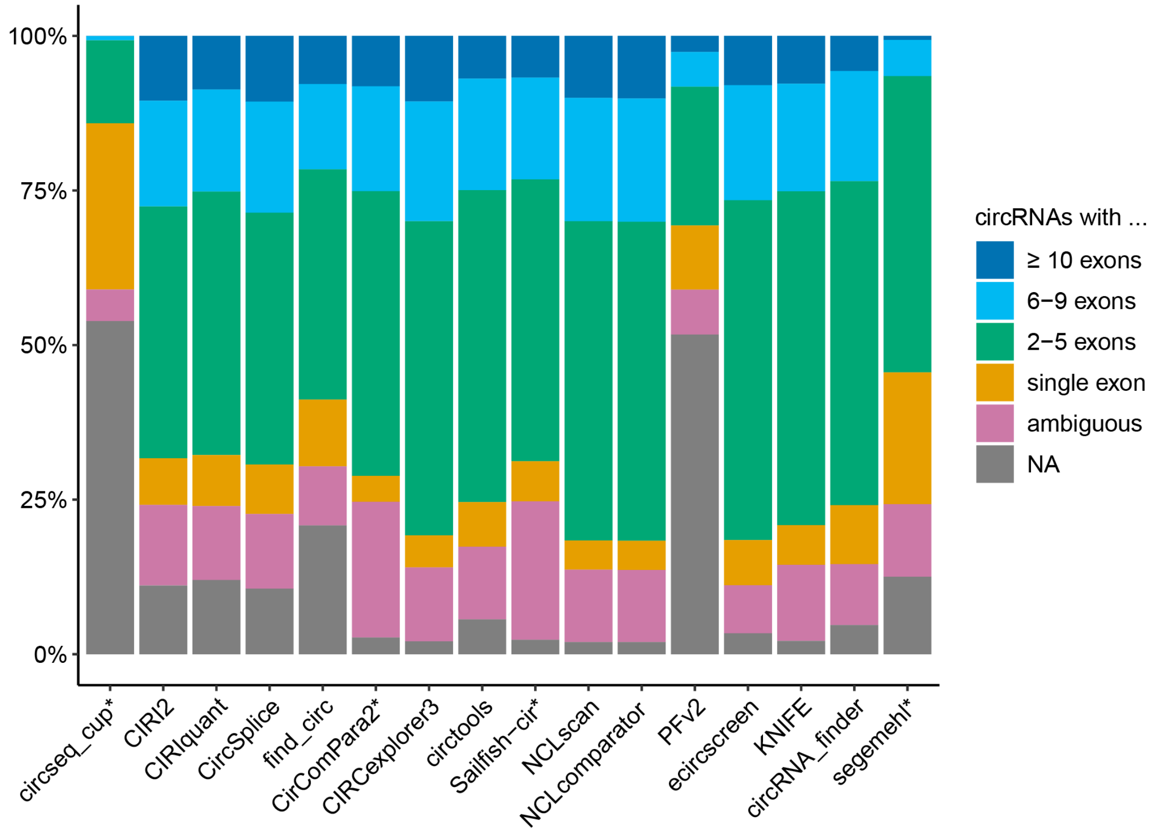


*tool does not report strand information

**Supplementary Figure 10** Most tools report approximately half of the circRNAs on the positive strand (median of 51.1%, IQR 50.9%-51.5%), similar to linear transcripts. ecircscreen reports more circRNAs (59.9%) on the positive strand. Ecircscreen is an integrative tool that reports structures predicted by at least three methods using different aligners, and then assigns strand information by consensus, potentially resulting in the strand bias observed. Of note, CirComPara2, circseq_cup, Sailfish-cir, and segemehl do not report circRNA strand information.

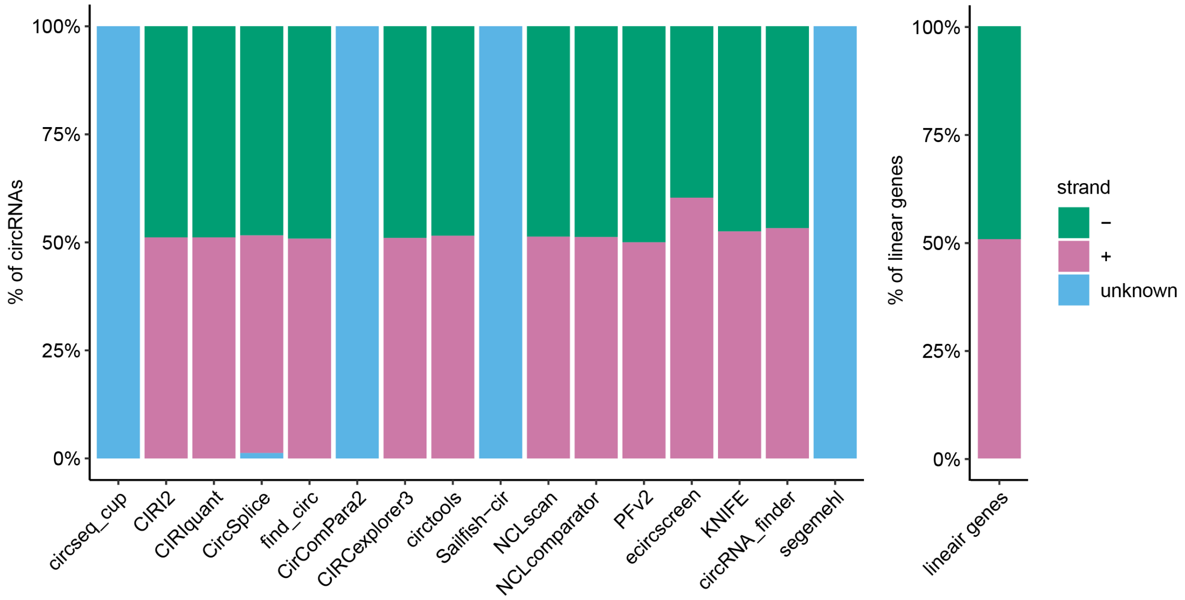


**Supplementary Figure 11** CircRNAs with lower BSJ counts are more difficult to detect with RT-qPCR due to their low abundance. Here, a threshold of Cq 32 is used assuming circRNAs under this threshold are high abundant enough to detect potential degradation upon RNase R treatment. The cumulative plot shows that above a certain BSJ count, circRNAs have a higher chance of being detected by RT-qPCR. Therefore, all data and validation rates were separately considered for circRNAs with a BSJ count lower than 5 (*low-abundance*), and circRNAs with a BSJ count of 5 or higher (*high-abundance*) (vertical grey dashed line).

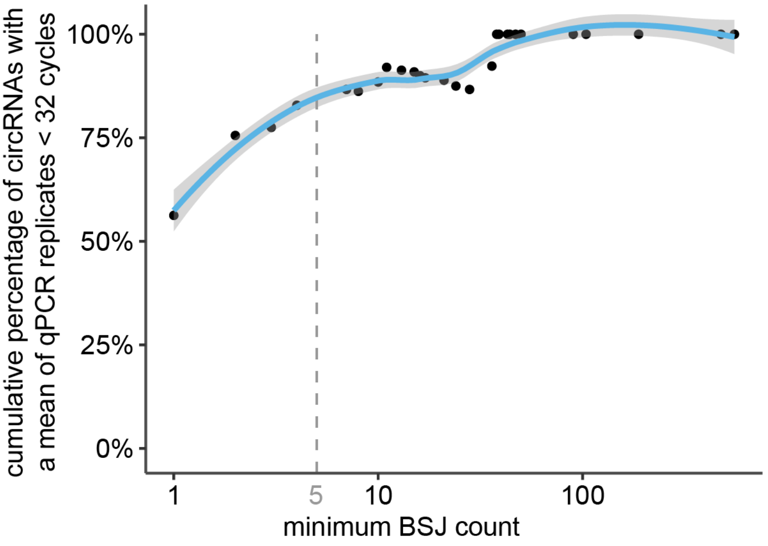

**Supplementary Figure 12** CircRNA primer design impacts final circRNA selection. Here, a random set of 1000 circRNAs per tool was selected (200 from circRNAs with a BSJ count < 5, 800 from circRNAs with a BSJ count ≥ 5), and primers were designed using CIRCprimerXL. Overall, most circRNA detection tools have similar primer design success rates (median 86.8%, IQR 85.1%-87.7%), apart from PFv2 with a primer design success rate of 51.4%. Remarkably, this lower design rate is only observed in the *high-abundance* circRNAs. When investigating why primer design was not successful, PFv2 showed a higher percentage (57.8%) of primers discarded due to predicted off-target amplification compared to the other tools (median 38.2%, IQR 35.1%-42.2%).

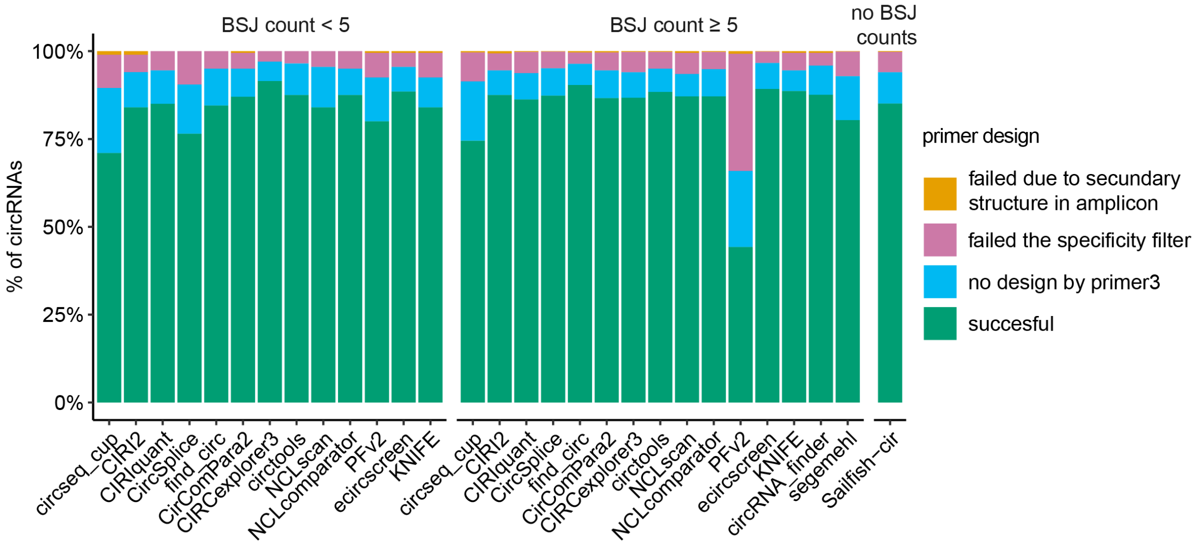


**Supplementary Figure 13** The BSJ count distribution of the 1,560 selected circRNAs. The upper and lower hinges of the box correspond to the first and third quartile (interquartile range, IQR), respectively. The upper and lower whiskers extend from the hinge to the highest and lowest value that is within 1.5 * IQR of the hinge, respectively. Data beyond the end of the whiskers are outliers and plotted as points.

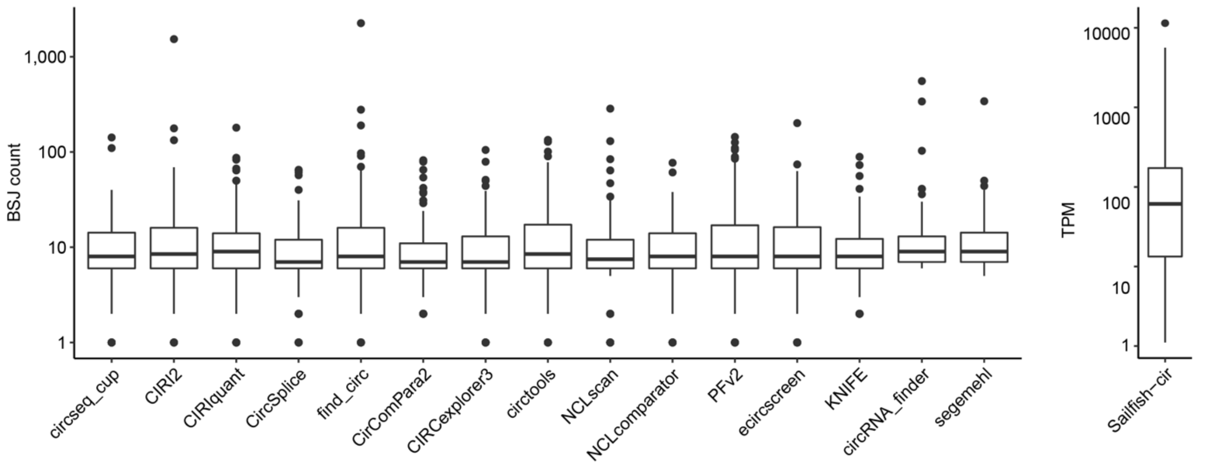

**Supplementary Figure 14** In total, 1,560 circRNA/tool/cell line tuples were randomly selected. As some circRNAs were selected more than once (for different tools) the total number of unique circRNA/cell line pairs is 1,516, and the number of unique circRNAs (not taking into account the originating DNA strand) is 1,457.

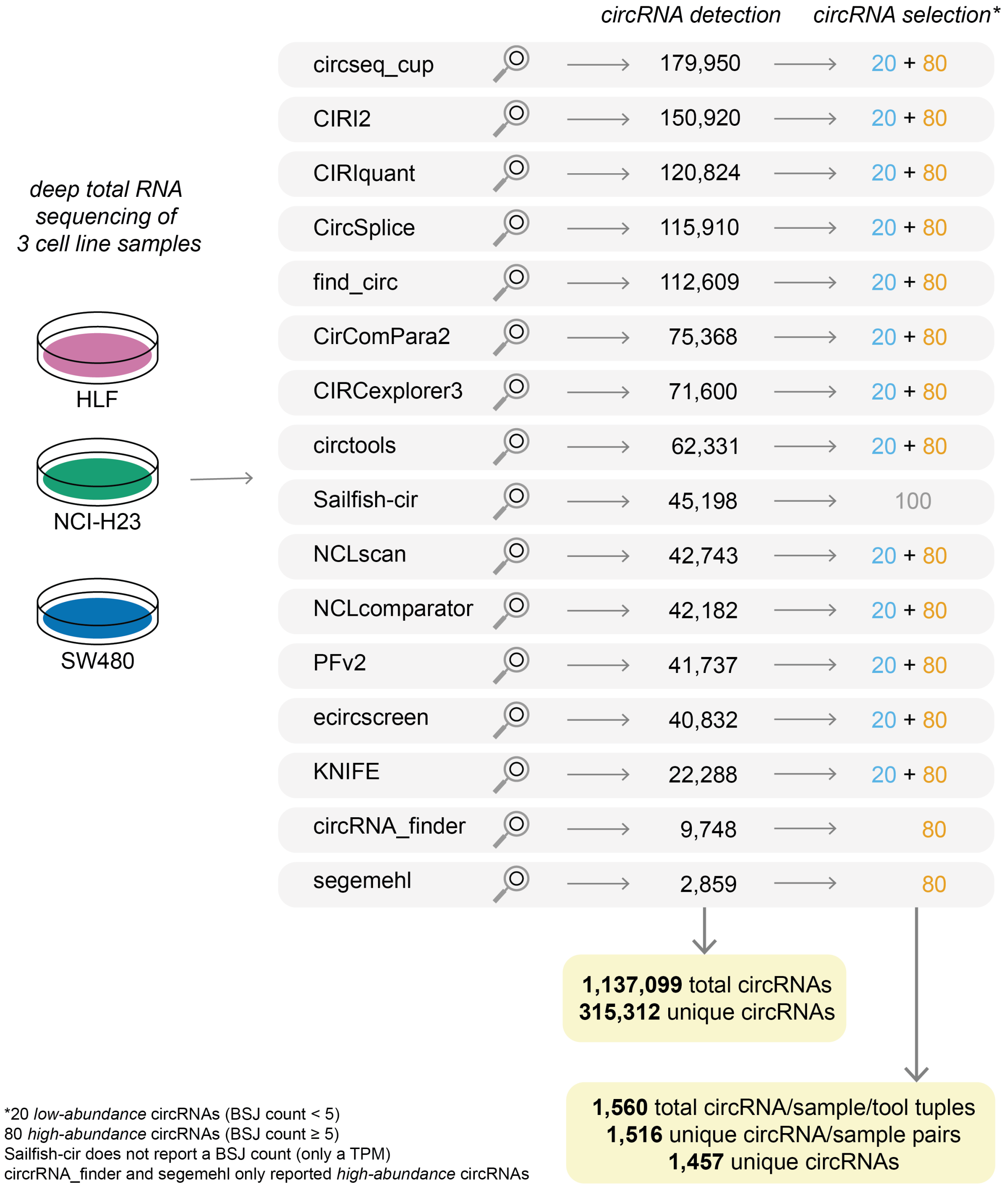

**Supplementary Figure 15** Cumulative plots of precision for each of the three empirical validation methods and the compound validation metric in function of the BSJ count level. The precision for each of the three validation methods is slightly lower for lower BSJ counts, but the differences are modest. The cumulative precision values were calculated by descending BSJ count, meaning that each dot shows the cumulative precision for all circRNAs with a BSJ count equal to or larger than the value shown on the x-axis. In other words, the most-left point of each graph represents the total precision value.

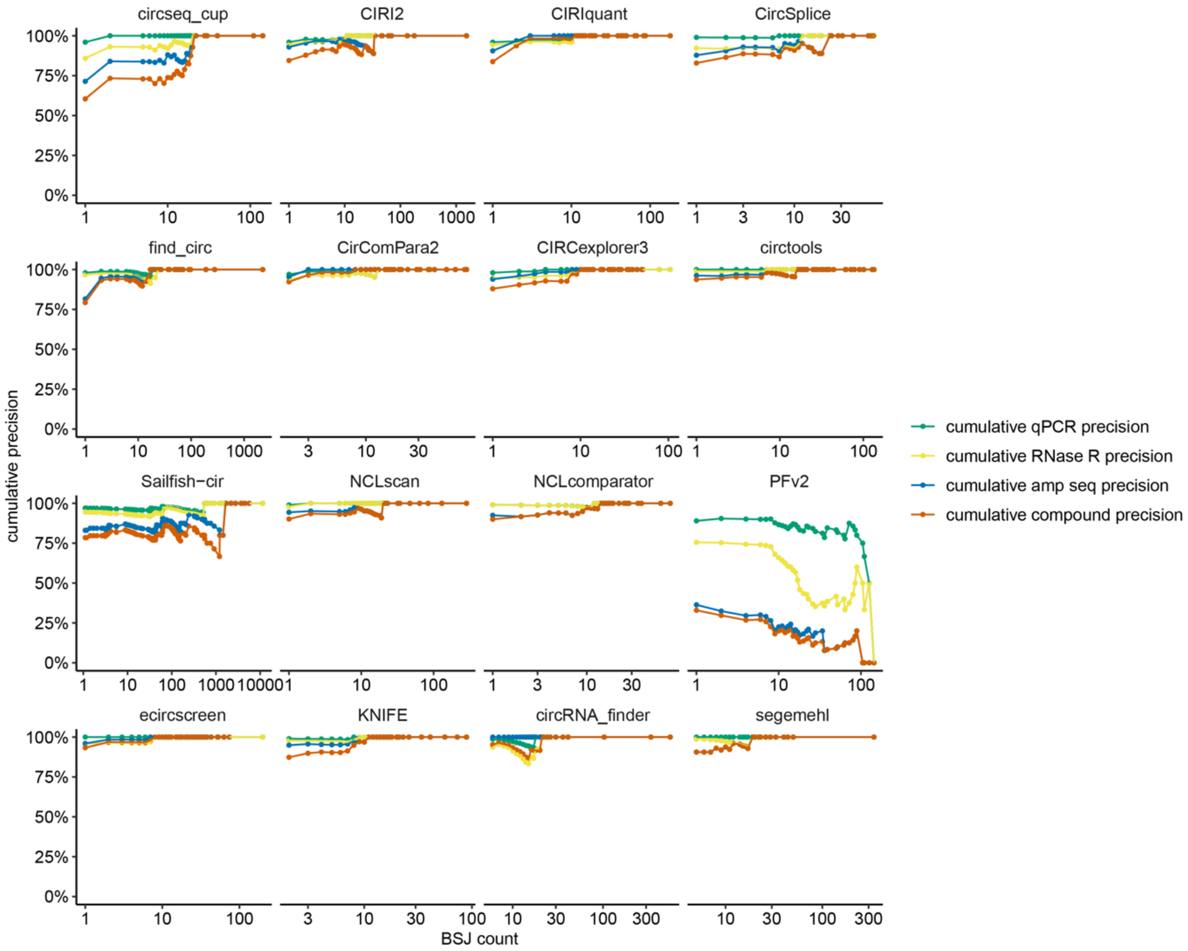

**Supplementary Figure 16** A Sashimi plot of linear and circular RNA sequencing counts (and counts per million (CPM)) after RNase R treatment of the NCI-H23 cell line shows the enrichment of the single-exonic circRNA circTEAD1(3) (chr11:12764178-12764434), with an enrichment factor (CPM_treated / CPM_untreated) of 4.1 (dashed lines). Furthermore, the degradation of linear RNA is visible (full lines). For example, there is an 8-fold increase in the circRNA to linear RNA ratio on the 3' end after RNase R treatment (comparing the circTEAD1(3) BSJ and the linear junction between TEAD1 exon 3 and 4). Of note, not all linear RNA is degraded as a relatively mild RNase R treatment was used. Of note, this circRNA example was purposely selected from a region with low-complexity splicing patterns to better illustrate the effect of RNase R treatment.

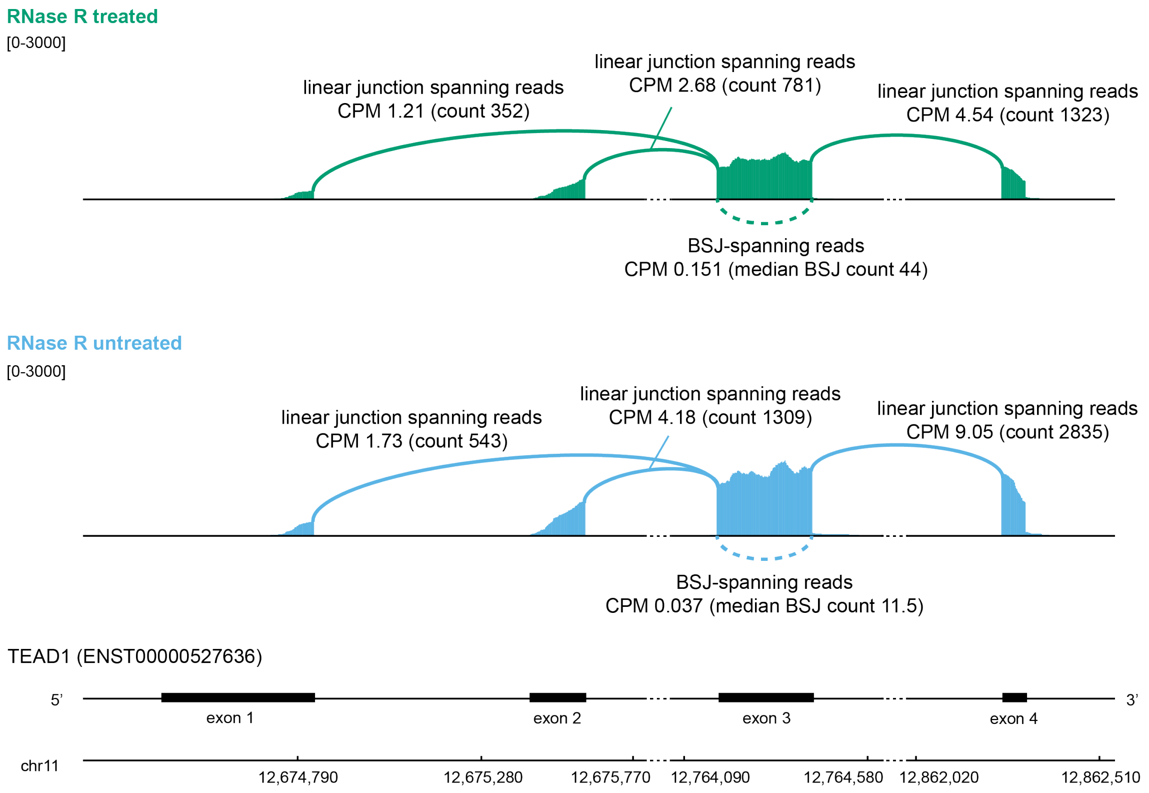

**Supplementary Figure 17** CircRNA enrichment in RNA sequencing CPM upon RNase R treatment (for *high-abundance* circRNAs). Some candidate circRNAs are only detected in the untreated sample; hence, no enrichment factor can be computed for these. For some tools, these represent a large fraction of the circRNAs (for example, PFv2, circseq_cup, Sailfish-cir).

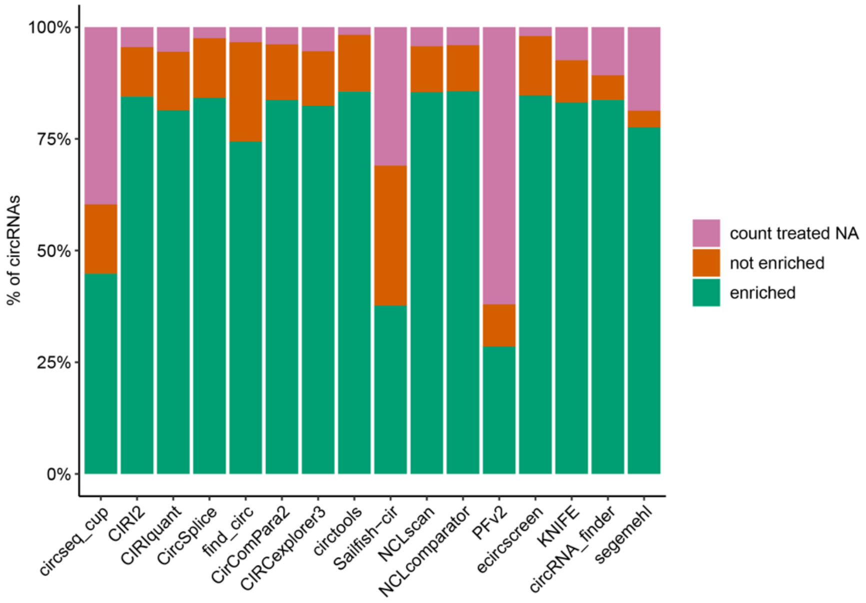


**Supplementary Figure 18** Distribution of the RNase R enrichment factors per tool (for *high-abundance* circRNAs). For some tools, this is an overestimation of the enrichment factor, as a substantial number of its predicted circRNAs are not detected in the RNase R treated sample and therefore no enrichment factor can be computed for these circRNAs. The short horizontal lines and the number above indicate the median enrichment for a given circRNA detection tool. The dashed grey horizontal line indicates an enrichment of 1, which is used as a threshold.

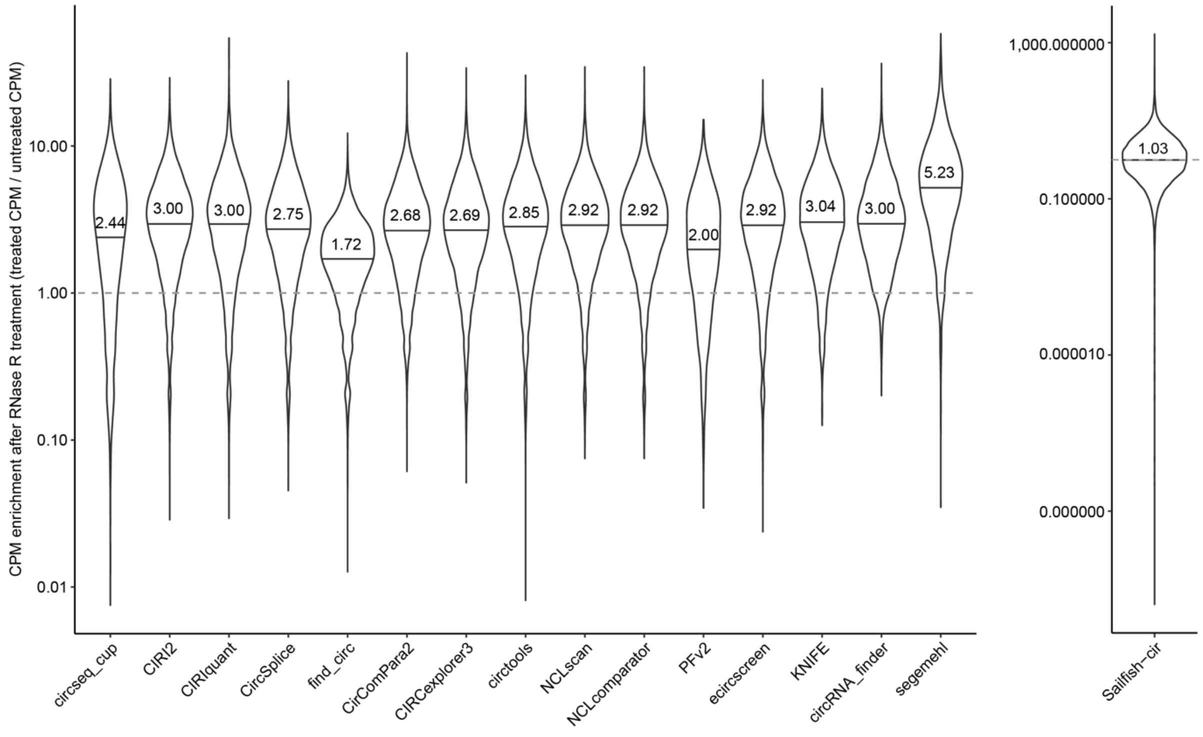

**Supplementary Figure 19** Cumulative percentage of enriched circRNAs after RNase R treatment, measured via RNA sequencing (for *high-abundance* circRNAs). The dashed grey vertical line indicates an enrichment of 1, which is used as a threshold to determine a circRNA as a true positive result. Curves more to the right indicate higher overall enrichment.

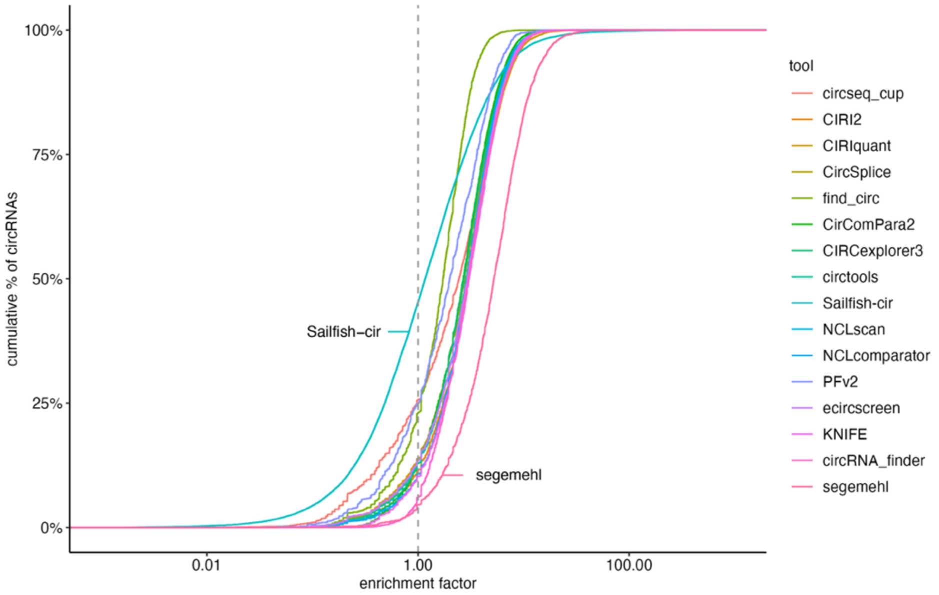


**Supplementary Figure 20** For each circRNA detection tool, a cumulative distribution plot showing the percentage of on-target amplification after amplicon sequencing was generated. A vertical grey dashed line at 50% on-target amplification is added to show the amplicon sequencing validation cut-off that was used. For example, for circRNA_finder, all selected circRNAs had an on-target amplification of at least 50%, as the curve only starts above the 50% threshold. It is noticeable that circRNAs with a BSJ count ≥ 5 have generally higher validation rates than circRNAs with a BSJ count < 5. Interestingly, for KNIFE and circtools, both BSJ count groups have a similar curve. For PFv2, the curve for BSJ counts ≥ 5 is lower than the one with BSJ counts < 5.

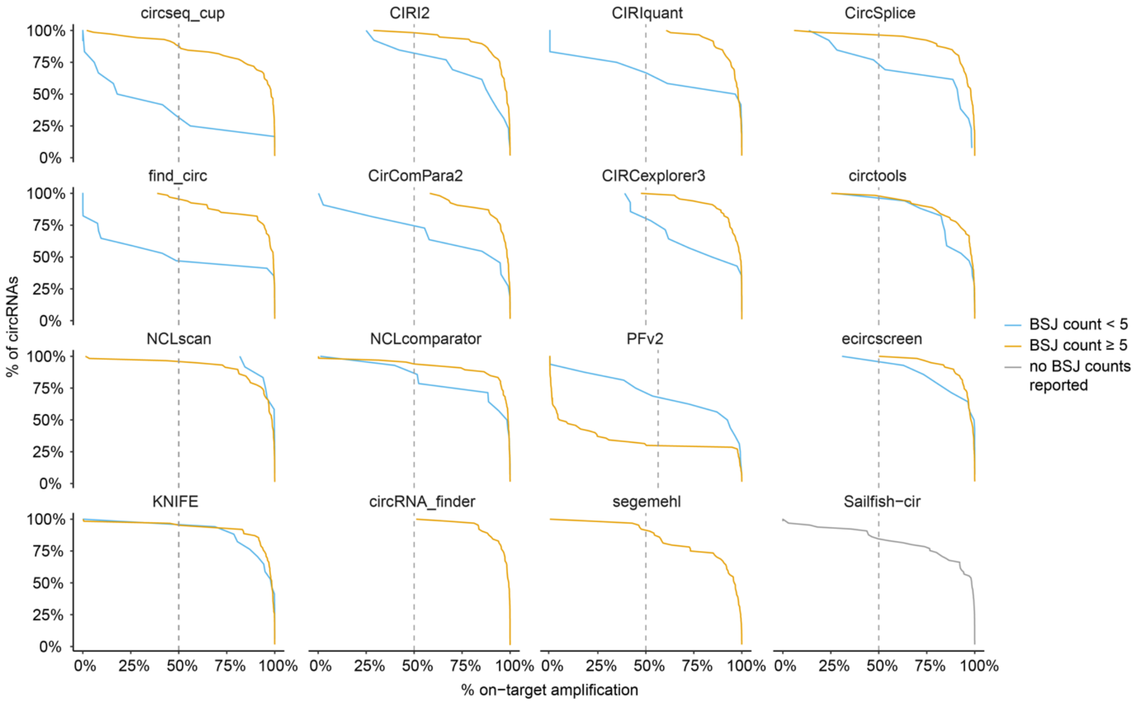

**Supplementary Figure 21** Simple display of the circRNA benchmarking validation results. For each tool, 80 circRNAs with a BSJ count ≥ 5, and 20 circRNAs with a BSJ count < 5 (if available) are represented. For each circRNA, three horizontal rectangles are shown, representing the three validation methods. This overview shows that the majority of selected circRNAs are validated by all three orthogonal validation methods. It also shows how some tools have a large number of circRNAs failing validation, especially in the subset of circRNAs with a BSJ count < 5. (The tools are sorted based on Figure 2A.)

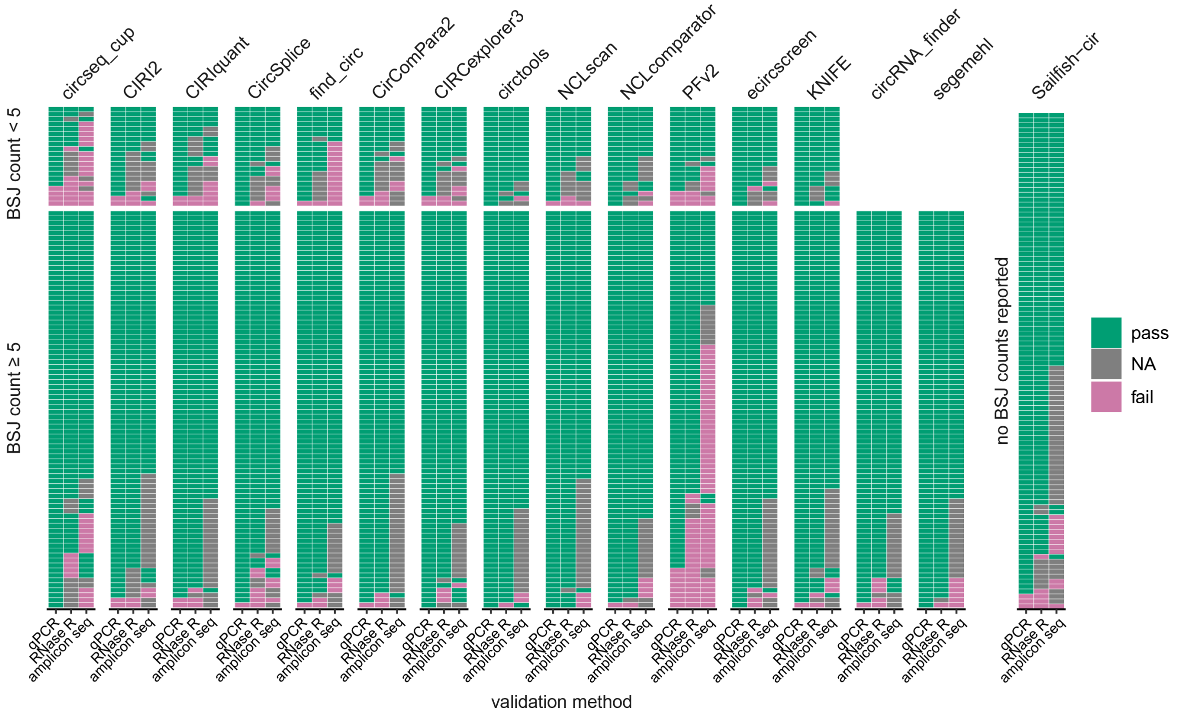


**Supplementary Figure 22** The compound precision value of most tools is high for *high-abundance* circRNAs (in orange).

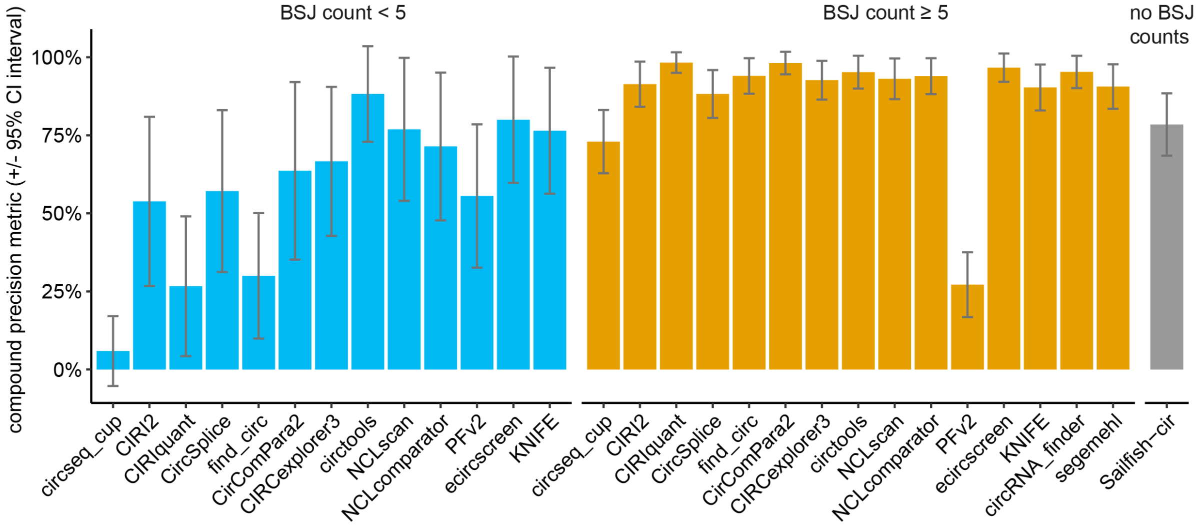

**Supplementary Figure 23** CircRNA tools differ in their sensitivity. circRNA_finder and segemehl filtered their results on circRNAs with a BSJ count ≥ 5, which explains why these tools have very low sensitivity in the BSJ count < 5 subset. This sensitivity metric is calculated based on a biased selection of circRNAs and should be used with caution (Supplementary Data 12).

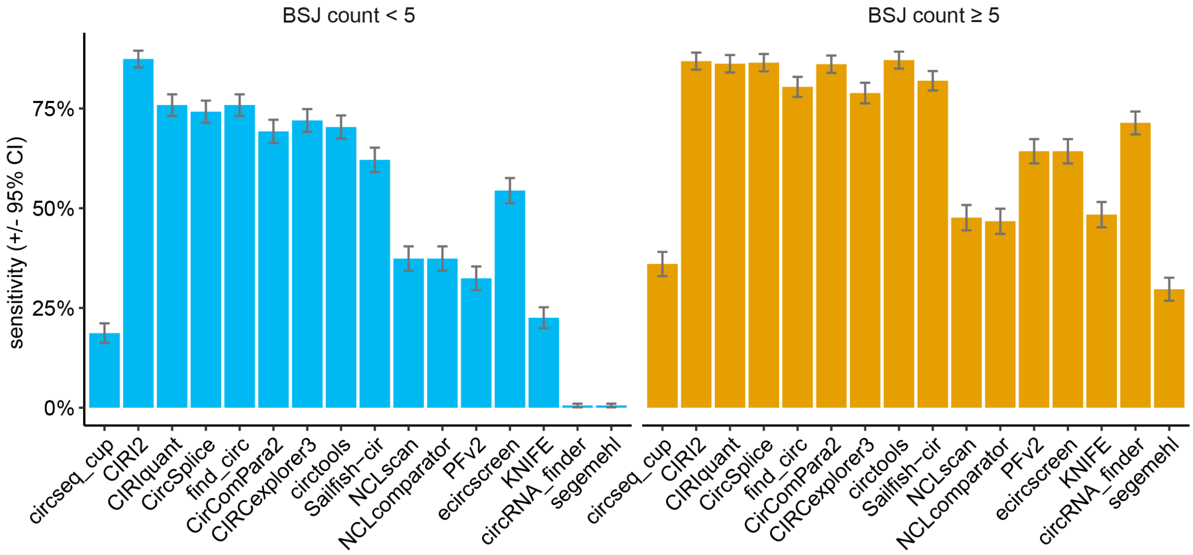


**Supplementary Figure 24** Cumulative sensitivity plots in function of the BSJ count level. There is a slight increase in sensitivity when the BSJ counts are higher, but the effect size is small. Somewhat surprisingly, this analysis demonstrates that some very high abundant circRNAs are not detected by certain tools. This is thus a tool-specific issue and not a general relationship between circRNA abundance and detection sensitivity. The cumulative sensitivity values were calculated by descending BSJ count, meaning that each dot shows the cumulative sensitivity for all circRNAs with a BSJ count equal to or larger than the value shown on the x-axis. In other words, the most-left point of each graph is the global sensitivity value.

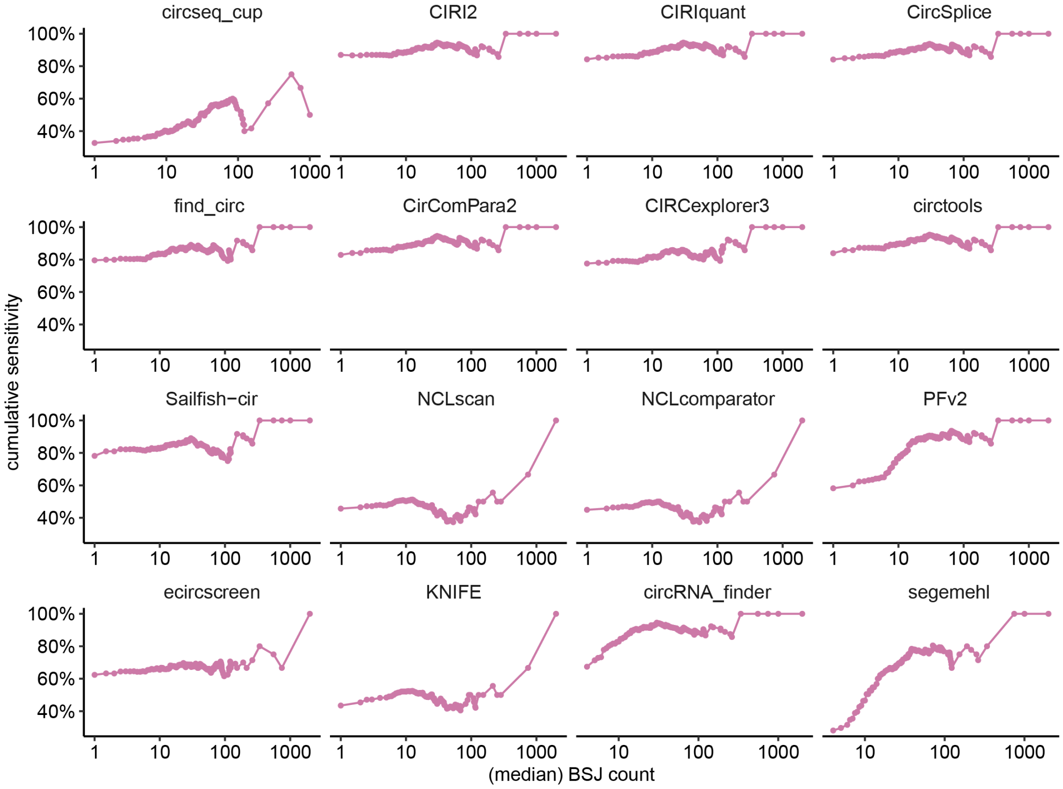

**Supplementary Figure 25** Using the compound precision value for each tool and BSJ count group, the theoretical number of true positive circRNAs based on the original number of predicted circRNAs is computed (*i.e.* the extrapolated sensitivity). A similar pattern is visible across all three cell lines.

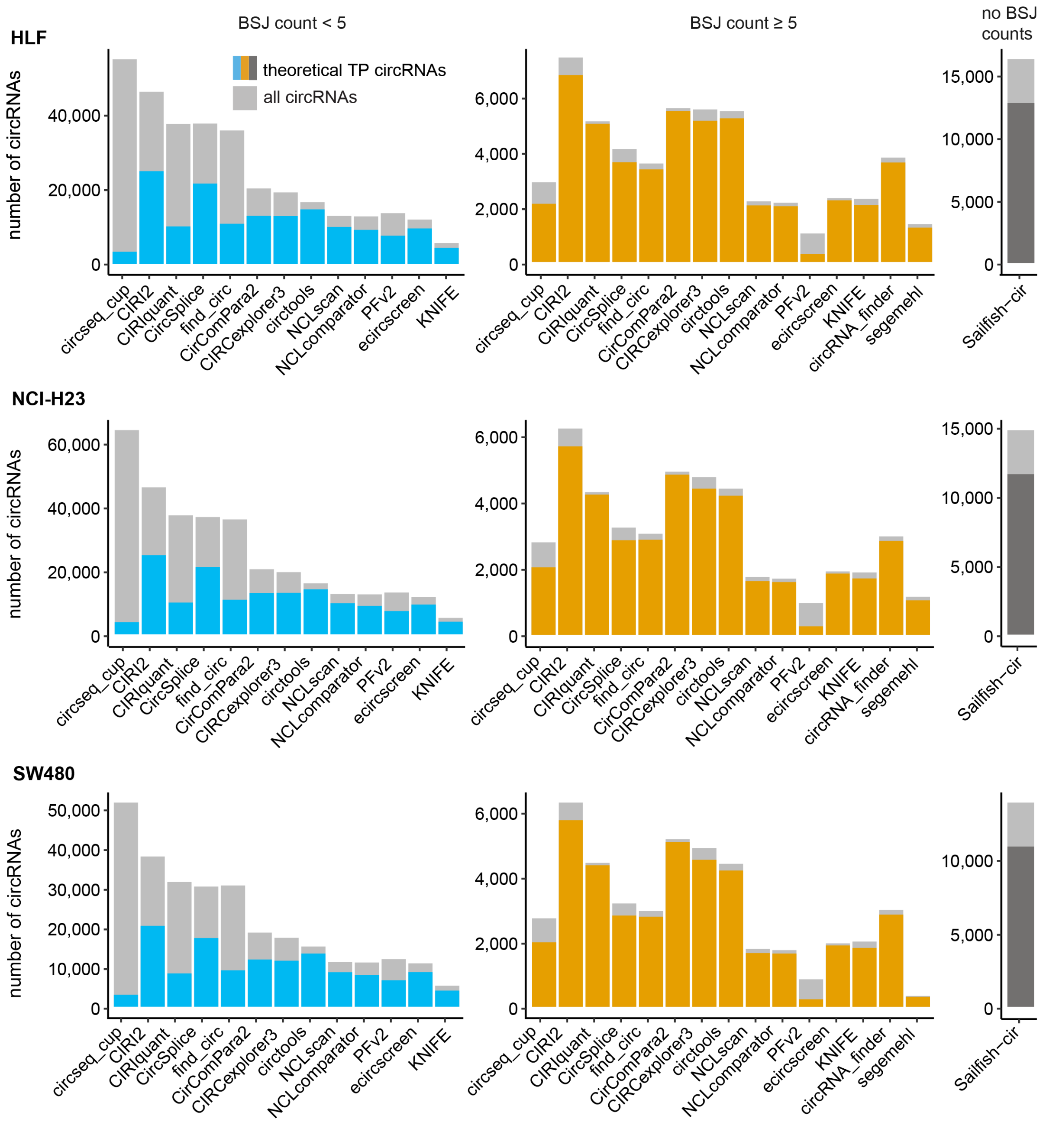

**Supplementary Figure 26** Precision-recall (sensitivity) dot plot for the 16 tools (for *high-abundance* circRNAs). The dashed grey diagonal line represents the y = x line.

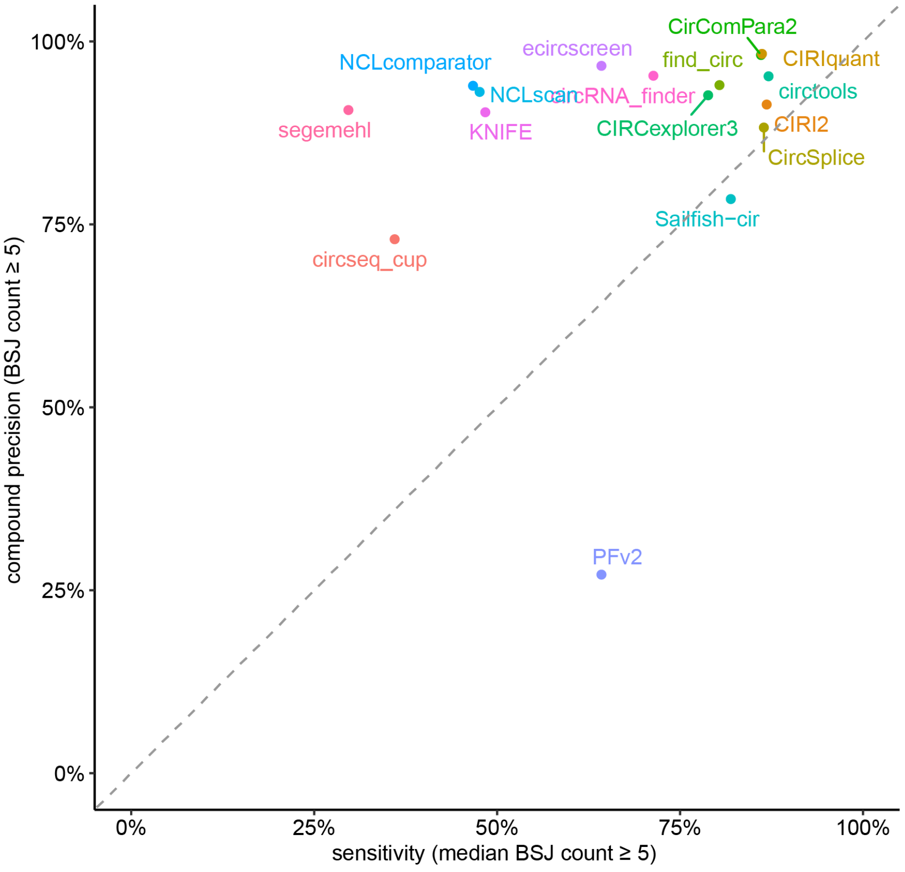

**Supplementary Figure 27** The circRNA detection tool precision (qPCR, RNase R, amplicon sequencing, and compound) is similar for the three separate cell lines (for *high-abundance* circRNAs).

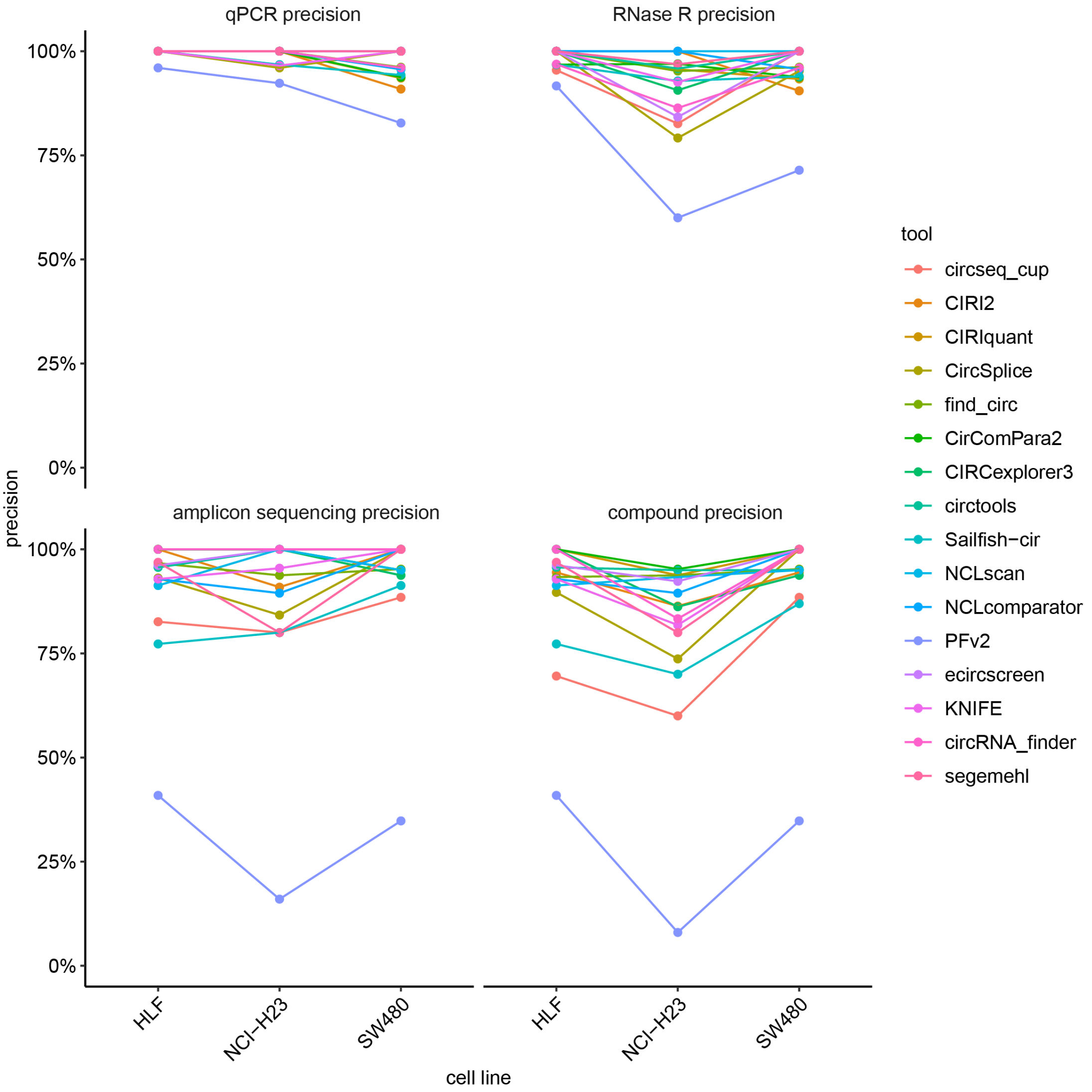

**Supplementary Figure 28** The circRNA detection tool sensitivity is similar for the three separate cell lines.

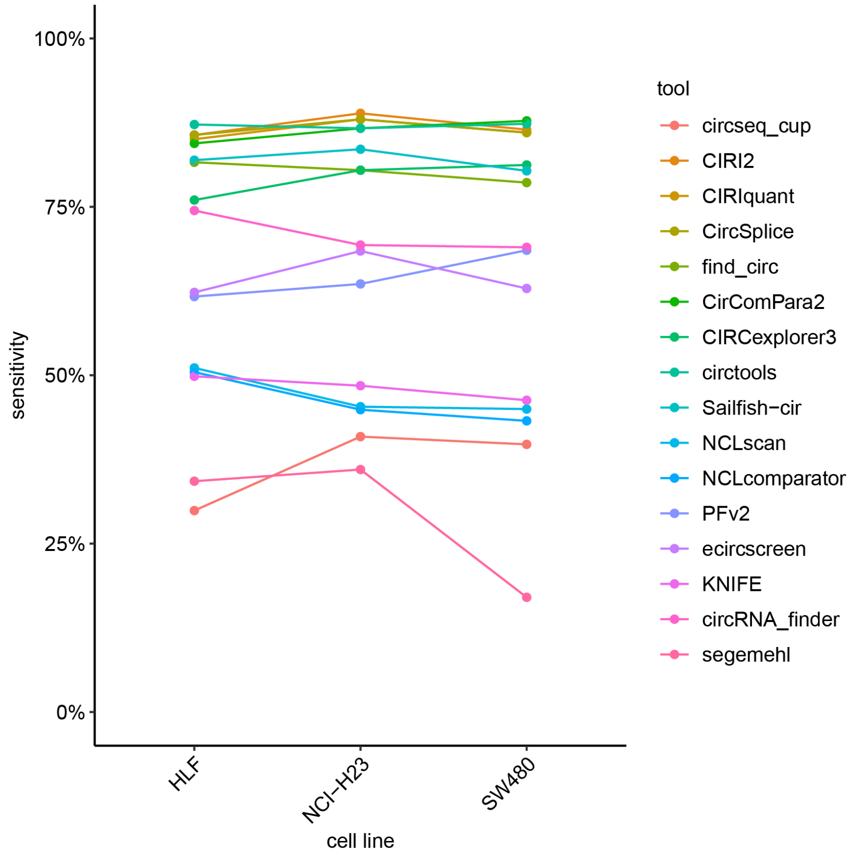


**Supplementary Figure 29** Correlation (based on linear model) of circRNA RT-qPCR values counts determined in different cell lines.

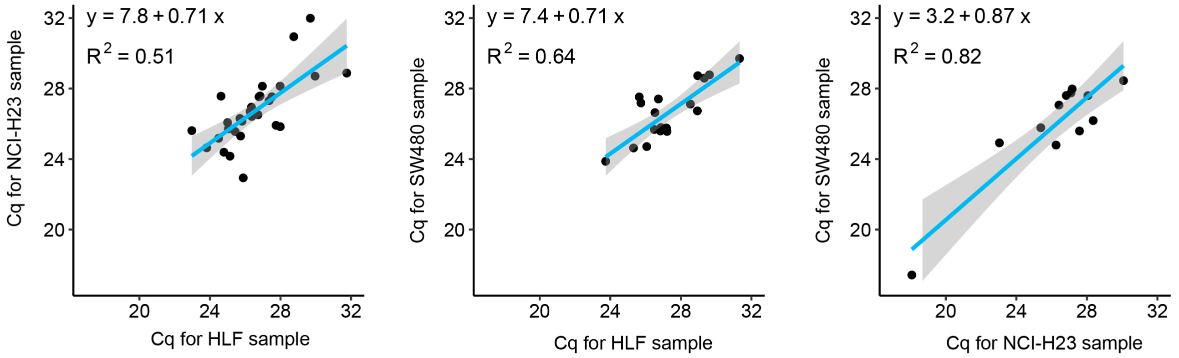

**Supplementary Figure 30** For 4/58 circRNAs detected in multiple cell lines, the RNase R validation was not the same for both cell lines. Comparison of RNase R treatment results of circRNAs in different cell lines.

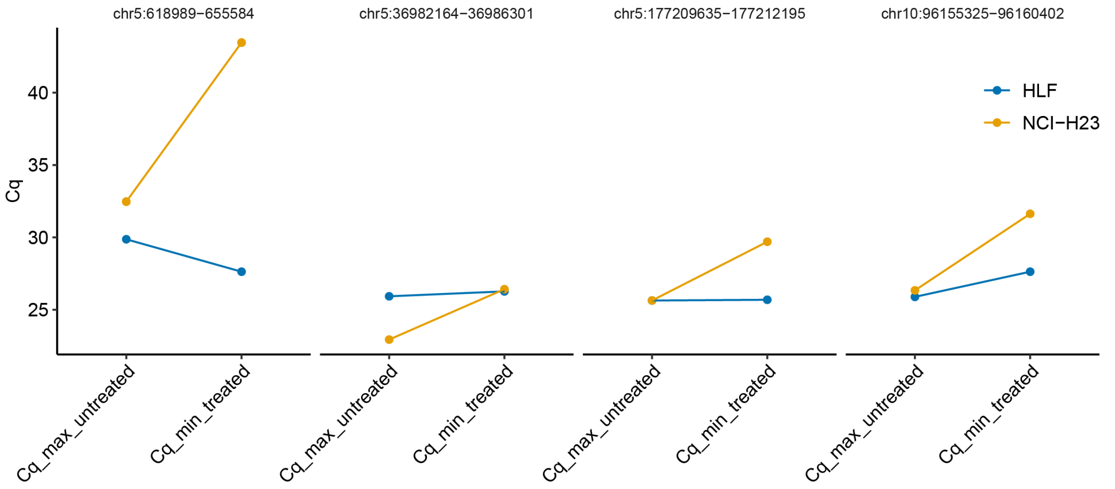


**Supplementary Figure 31** Linear regression of BSJ count (x-axis) and Cq values (y-axis).

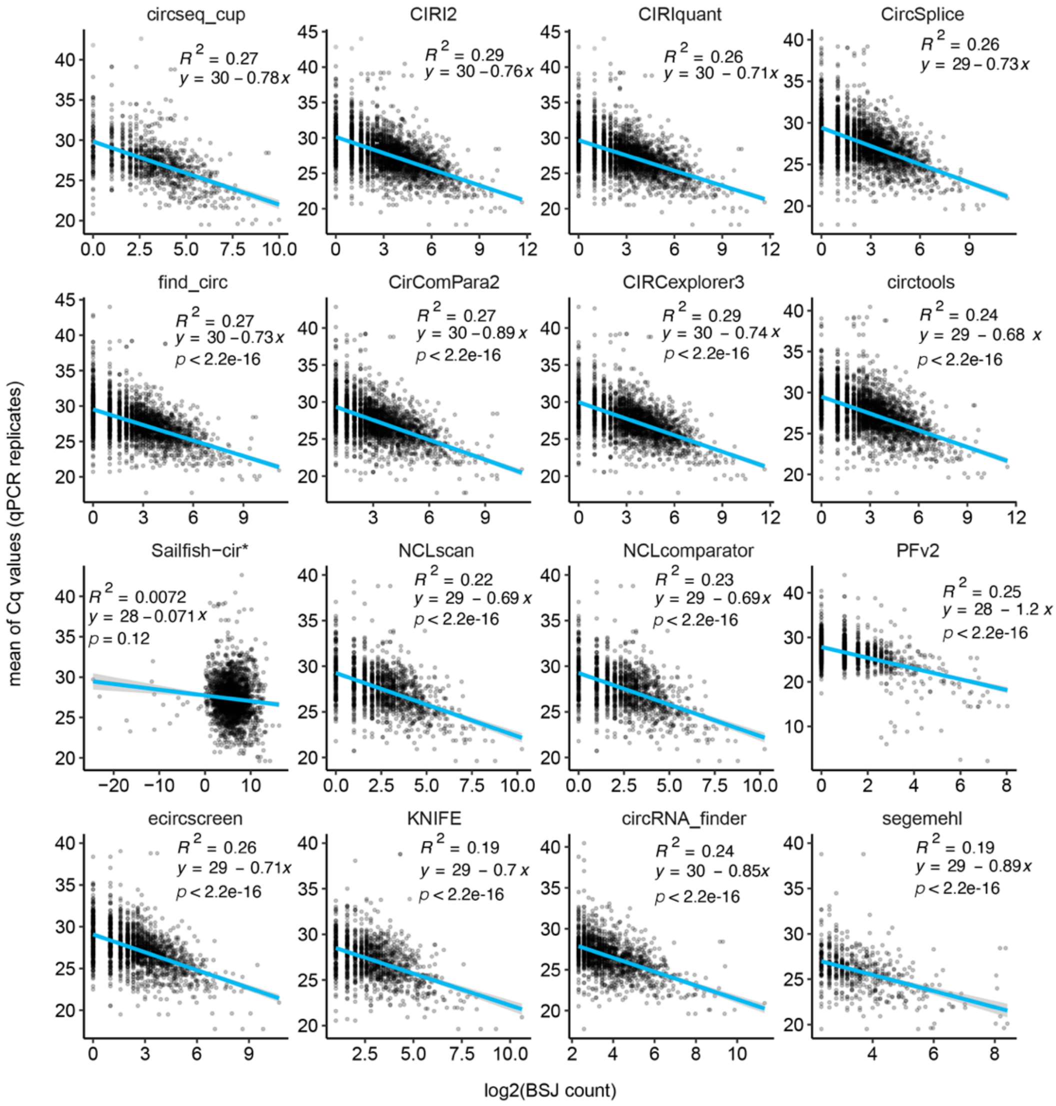


*Sailfish-cir, does not report raw BSJ counts, but transcripts per million (TPM) instead.

**Supplementary Figure 32** CircRNAs previously described in databases are more often validated compared to novel circRNAs.

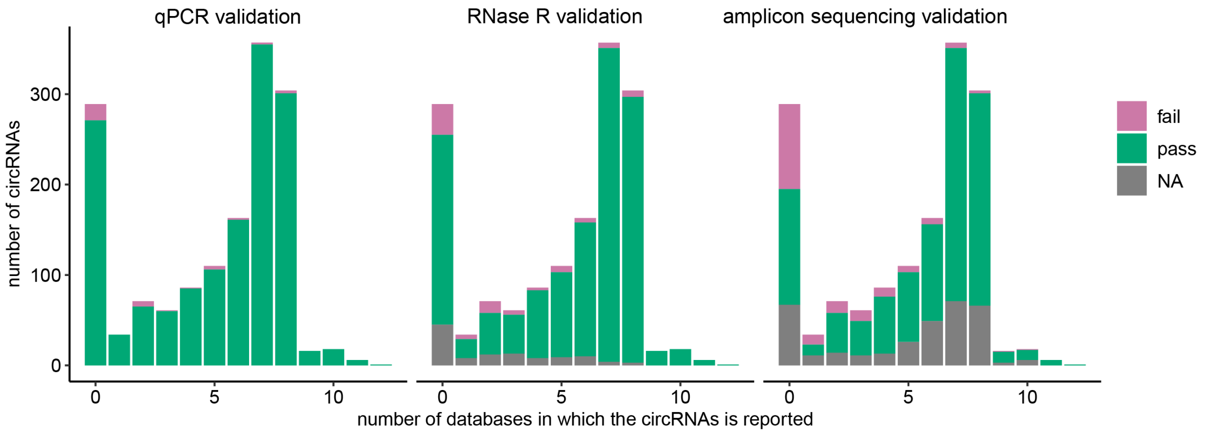

**Supplementary Figure 33** CircRNA detection tools can be combined by using the union of two tools, resulting in a larger set of detected circRNAs. Here, all combinations with a minimum 7.5% increase in the number of circRNAs are represented (percentage calculated with respect to the total number of circRNAs predicted for that cell line). For the y-axis, the percentage of detected circRNAs is calculated by dividing the number of circRNA detected by that tool combination by the total number of predicted circRNAs for that sample taking the union of all tools (13,087, 11,272, and 11,026 for the HLF, NCI-H23, and SW80 sample, respectively).

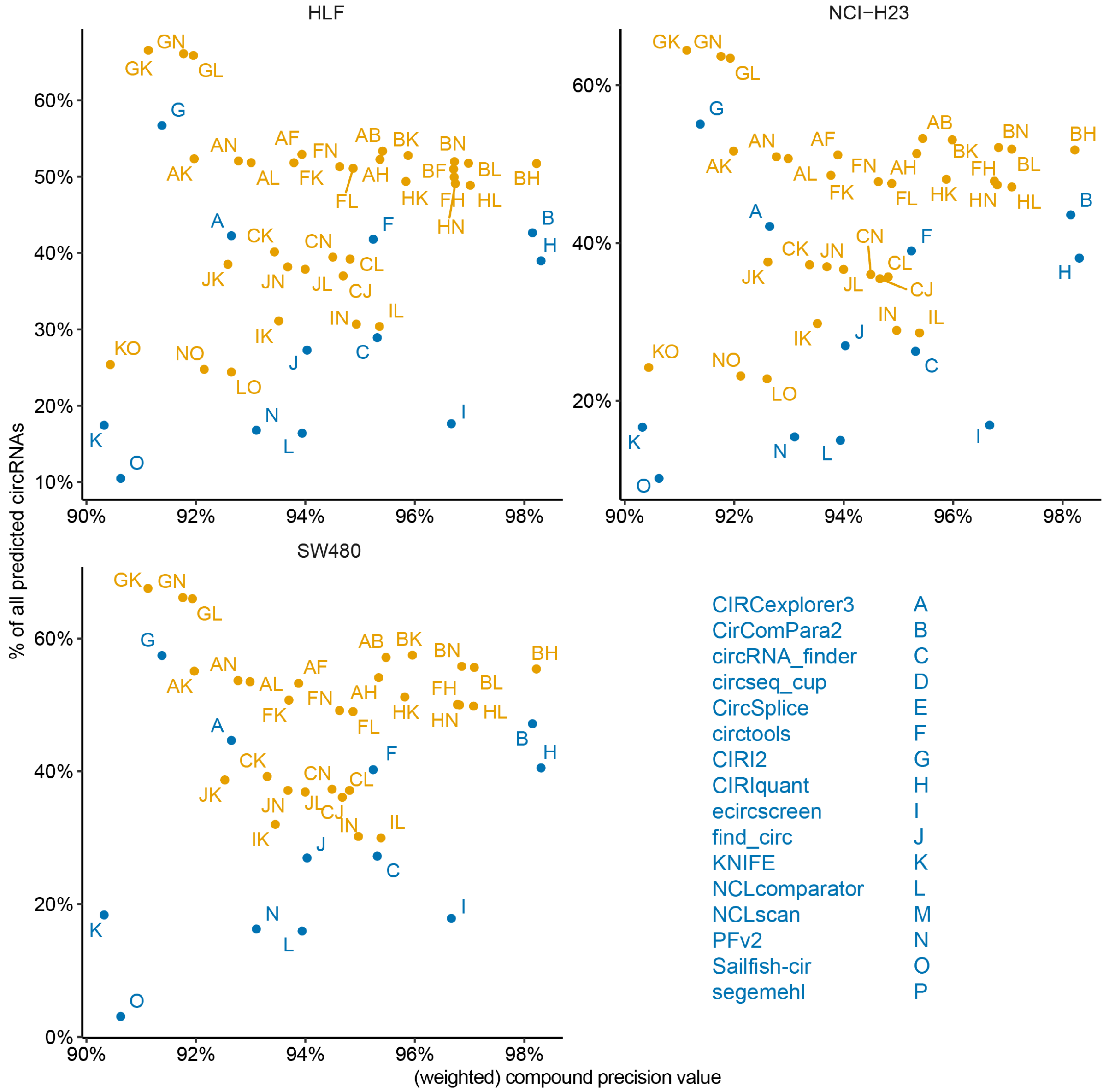

**Supplementary Figure 34** The union of the results of three circRNA detection tools can be used to increase the number of detected circRNAs (for visualization purposes, only combinations for which at least 9,000/13,087 possible circRNAs are present in the union of the three tools; only shown for HLF).

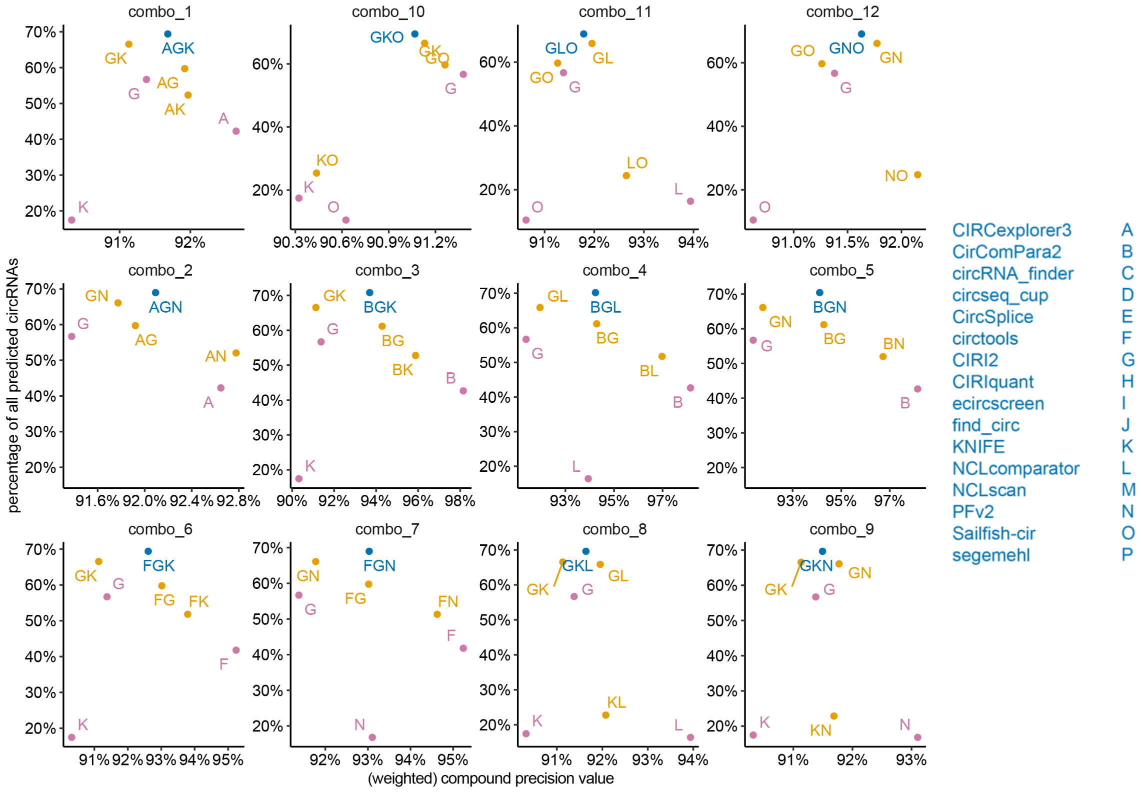

**Supplementary Figure 35** The precision values based on our validated dataset are comparable to previous studies with simulated data (30, 32). The dashed grey diagonal line represents the y = x line. PFv2 is an outlier due to its tendency to predict circRNAs in regions with repeat sequences, which are not present in the simulated datasets.

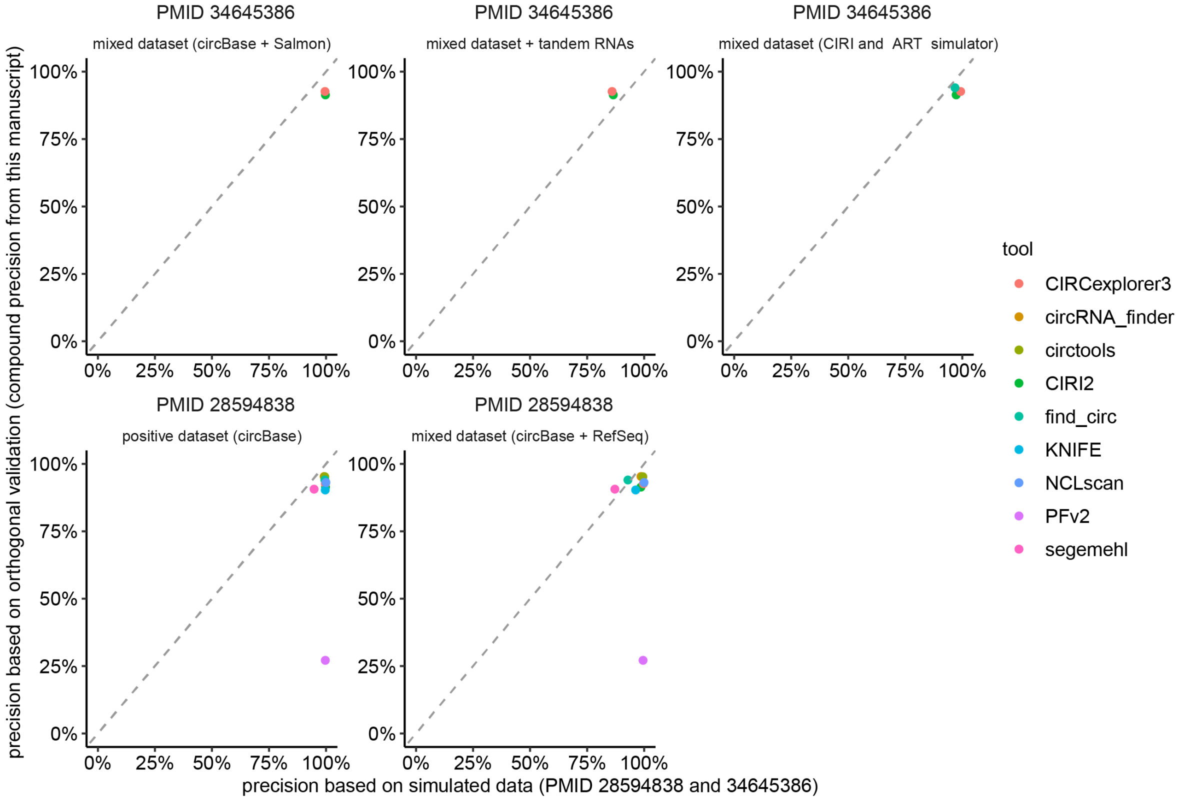

**Supplementary Figure 36** Sensitivity is highly variable among tools, both based on our validated dataset and on previously published simulated data (30, 32). The dashed grey diagonal line represents the y = x line.

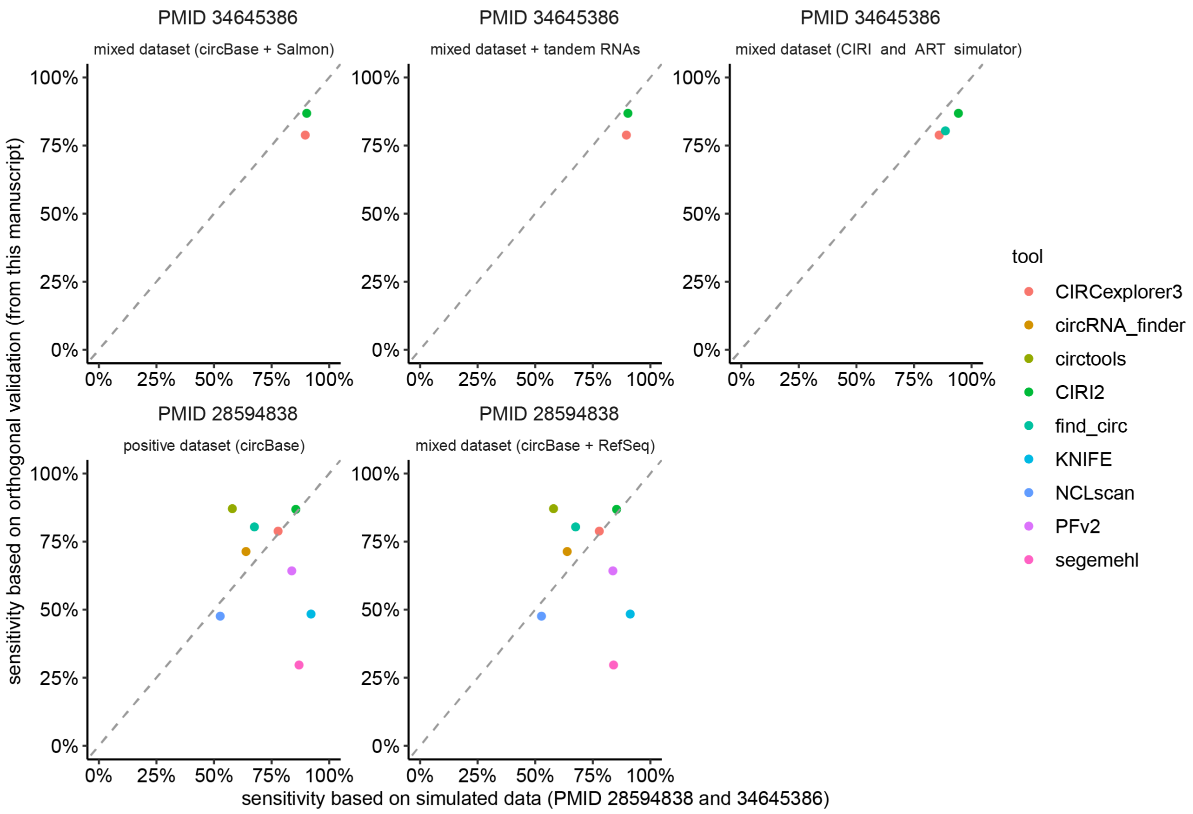
